# Establishing a Role of the Semantic Control Network in Social Cognitive Processing: A Meta-analysis of Functional Neuroimaging Studies

**DOI:** 10.1101/2021.04.01.437961

**Authors:** Veronica Diveica, Kami Koldewyn, Richard J. Binney

## Abstract

Most leading models of socio-cognitive processing devote little discussion to the nature and neuroanatomical correlates of cognitive control mechanisms. Recently, it has been proposed that the regulation of social behaviours could rely on brain regions specialised in the controlled retrieval of semantic information, namely the anterior inferior frontal gyrus (IFG) and posterior middle temporal gyrus. Accordingly, we set out to investigate whether the neural activation commonly found in social functional neuroimaging studies extends to these ‘semantic control’ regions. We conducted five coordinate-based meta-analyses to combine results of over 500 fMRI/PET experiments and identified the brain regions consistently involved in semantic control, as well as four social abilities: theory of mind, trait inference, empathy and moral reasoning. This allowed an unprecedented parallel review of the neural networks associated with each of these cognitive domains. The results confirmed that the anterior left IFG region involved in semantic control is reliably engaged in all four social domains. This suggests that social cognition could be partly regulated by the neurocognitive system underpinning semantic control.

## 1. Introduction

The ability to comprehend and respond appropriately to the behaviour of others is essential for humans to survive and thrive. A major challenge for the cognitive sciences, therefore, is to characterise *how* we understand others and coordinate our behaviour to achieve mutually beneficial outcomes, and what can cause this ability to break down (Frith, 2007). There is an indubitable requirement for systems that control, or regulate, the cognitive processes underpinning social interactions. This is because social interactions are intricate and fraught with the potential for misunderstandings and faux pas; first, the everyday social signals to which we are exposed are typically complex, often ambiguous and sometimes conflicting. This is compounded by the fact that the meaning of a given gesture, expression or utterance can vary across contexts (Barrett et al., 2011; Rodd, 2020). Moreover, once we have settled upon an interpretation of these signals, we are then faced with the additional challenge of selecting an appropriate response, and inhibiting others which might, for example, be utilitarian but socially insensitive or even damaging. In order to undergo social interactions that are coherent, effective and context-appropriate, we must carefully regulate both our comprehension of, and response to, the intentions and actions of others (Binney and Ramsey, 2020; Fujita et al., 2014; Gilbert and Burgess, 2008; Ramsey and Ward, 2020).

Despite there being a wealth of literature describing executive functions involved in general cognition (Assem et al., 2020; Diamond, 2013; Duncan, 2013, 2010; Fedorenko et al., 2013; Petersen and Posner, 2012), prominent models of socio-cognitive processing are under-specified in terms of the contribution and neural basis of cognitive control mechanisms (e.g., Adolphs, 2009, 2010; Frith & Frith, 2012; Lieberman, 2007). For example, Adolphs (2009; 2010) only very briefly refers to cognitive processes involved in ‘social regulation’ and largely within the limited context of emotional regulation. Likewise, Frith and Frith (2012) refer to a “supervisory system” which has the characteristic features of executive control, but its functional and anatomical descriptions lack detail important for generating testable hypotheses. However, research into specific social phenomena, such as prejudice (Amodio, 2014; Amodio and Cikara, 2021) and automatic imitation (Cross et al., 2013; Darda and Ramsey, 2019) has recently begun to give the matter of cognitive control greater attention. Of particular interest has been the contribution of the domain-general multiple-demand network (MDN), a set of brain areas engaged by cognitively-challenging tasks irrespective of the cognitive domain (Assem et al., 2020; Duncan, 2010; Fedorenko et al., 2013; Hugdahl et al., 2015). MDN activity increases with many kinds of general task demand, including working memory load and task switching, and it has been suggested that this reflects the implementation of top-down attentional control and the optimal allocation of cognitive resources to meet immediate goals (Duncan, 2013, 2010). The MDN is comprised of parts of the precentral gyrus, the middle frontal gyrus (MFG), the intraparietal sulcus (IPS), insular cortex, the pre-supplementary motor area (pre-SMA) and the adjacent cingulate cortex (Assem et al., 2020; Fedorenko et al., 2013), some of which have been implicated in controlled social processing such as, for example, working memory for social content (Meyer et al., 2012), social conflict resolution (Zaki et al., 2010), inhibition of automatic imitation (Darda and Ramsey, 2019) and mental state inference or theory of mind (ToM) (e.g. Rothmayr et al., 2011; Samson et al., 2005; Van der Meer et al., 2011). However, there are at least three key unresolved questions regarding the role of cognitive control in social cognition. First, it remains to be seen whether there could be multiple, distinguishable mechanisms of, and neural systems for, control. Second, it is unclear whether there exists a subset of control systems that are specialised towards processing social information and, third, we have little understanding as to whether certain types of control are necessary for all or only select social behavioural phenomena. Shedding light on these issues has the potential to generate important new hypotheses regarding social behaviour both in the context of health and injury/disease.

It has recently been proposed that a relatively specialised form of cognitive control, termed *semantic control*, could be particularly important for social cognitive processing (Binney and Ramsey, 2020). This follows a broader claim that social cognition and its neural correlates can be understood as a nuanced form of *semantic cognition* which itself is defined as a set of processes involved in extracting meaning from the environment and using it to guide purposeful and context-appropriate behaviour (Binney and Ramsey, 2020; Lambon Ralph et al., 2017). This framework contrasts with approaches that look upon social processing as a distinct or even special case of cognition (i.e., domain-specific models; Barrett, 2012; Saxe, 2006; but also see Amodio, 2019; Amodio and Cikara, 2021; Schaafsma et al., 2015; Spunt and Adolphs, 2017) and, instead, posits that it is underpinned by two, more domain-general neurocognitive systems. The first system is representational in nature and supports the acquisition and long-term storage of conceptual-level knowledge about objects, people, abstract concepts, and events. The anterior temporal cortices act as a central, supramodal semantic store through interaction with modality-specific and lower-order heteromodal association cortices (Binney et al., 2010; Kuhnke et al., 2021; Lambon Ralph et al., 2017; Patterson et al., 2007; Pobric et al., 2010). The second system, the semantic control system, modulates activation of semantic knowledge to bring to the fore aspects of conceptual information that are relevant to the situational context or the task at hand while inhibiting irrelevant aspects (Chiou et al., 2018; Jefferies, 2013; Lambon Ralph et al., 2017).

The reasons why semantic control should be critical for social cognition and interaction are uncomplicated; we retain a vast amount of socially-relevant knowledge including knowledge about familiar people (Greven et al., 2016; Hassabis et al., 2014), about the structure of and relationship between social categories and their associated stereotypes (Freeman and Johnson, 2016; Quinn and Rosenthal, 2012), and an understanding of abstract social concepts, norms and scripts (Frith and Frith, 2003; Van Overwalle, 2009). But only a limited portion of this information is relevant in a given social instance and it would be computationally inefficient to automatically retrieve it all. For example, there is no need to retrieve information about someone’s personality traits, or personal interests and hobbies, if the only task is to pick them out from within a crowd. Moreover, the types and the scope of information we need to retrieve to understand and respond appropriately to certain social signals change according to the context, and irrelevant information could potentially interfere. Therefore, semantic control is essential for limiting semantic retrieval according to the circumstances and avoiding potential social errors.

There is a growing body of convergent computational modelling, patient, neuroimaging and neuromodulation evidence that the semantic control system is supported by a neural network that is distinct from that underpinning semantic representation (e.g., Corbett et al., 2009; Davey et al., 2016, 2015; Jackson, 2021; Jefferies et al., 2008; Jefferies and Lambon Ralph, 2006; Teige et al., 2018). Specifically, semantic control engages regions of the MDN, as well as the semantic control network (SCN) which comprises the anterior IFG and the posterior middle temporal gyrus (pMTG) (Badre et al., 2005; Davey et al., 2016; Jackson, 2021; Noonan et al., 2013). Moreover, while the domain-general MDN is engaged by semantic tasks, and particularly those with high control demands (Jackson, 2021; Thompson et al., 2018), there is evidence that both the anatomy of the SCN and MDN and their functional contributions to controlled semantic processing are at least partially distinct (Gao et al., 2020). In particular, fMRI studies reveal that the mid-to posterior IFG (pars triangularis and pars opercularis), nodes of the MDN, have been shown to increase activity in response to increased ‘semantic selection’ demands, a process that is engaged when automatic retrieval of semantic knowledge results in competition between multiple representations which must be resolved (for example, hearing the word *bank* might elicit retrieval of the concept of a riverside and a financial institution)(Badre and Wagner, 2007; Nagel et al., 2008; Thompson-Schill et al., 1997). However, this mid-to posterior IFG region is also engaged by other non-semantic forms of response competition (Badre et al., 2005; Barredo et al., 2015) and tests of inhibitory function such as the Stroop task (Huang et al., 2020; January et al., 2009; Nee et al., 2007). In contrast, activation of the anterior IFG (pars orbitalis) appears to be more selective to semantic control demands and driven specifically by an increased need for ‘controlled semantic retrieval’, a mechanism that is engaged when automatic semantic retrieval fails to activate semantic information necessary for the task at hand, and a further goal-directed semantic search needs to be initiated (Badre and Wagner, 2007; Krieger-Redwood et al., 2015).

To date, there have been but a few neuroimaging investigations that have directly questioned the involvement of the SCN in social cognitive processing. Two recent fMRI studies compared activation during semantic judgements made on social and non-social stimuli and found that the IFG and pMTG were engaged by both stimulus types (Binney et al., 2016; Rice et al., 2018). Further, Satpute et al., (2014) found that controlled retrieval, but not selection of social conceptual information engages the anterior IFG. However, we are not aware of any prior studies that attempt to examine the engagement of the SCN specifically during tasks that are commonly viewed as social in nature (e.g., ToM tasks). As a starting point, rather than conducting a novel individual experiment, the present study adopted a meta-analytic approach to extract reliable trends from large numbers of studies. Meta-analyses of functional neuroimaging data overcome the limitations of individual studies (Cumming, 2014; Eickhoff et al., 2012), which are frequently statistically underpowered (Button et al., 2013) and vulnerable to effects of idiosyncratic design and analytic choices (Botvinik-Nezer et al., 2020; Carp, 2012) so that it becomes difficult to distinguish between replicable and spurious findings and to generalize the results. Our principal aim was to determine whether the distributed neural activation commonly associated with functional neuroimaging studies of social cognition extends to the neural networks underpinning semantic control (i.e., SCN and MDN). In order to localise the brain network sensitive to semantic control demands (i.e., semantic retrieval and/or selection), and then compare and contrast it to networks implicated in social cognition, we performed an update of Noonan et al.’s (2013) meta-analysis of semantic control (also see Jackson, 2021a).

We took the approach of investigating multiple sub-domains of social cognition in parallel because this should allow an assessment of the extent to which inferences are generalisable, rather than specific to certain types of social tasks and/or abilities. We chose to focus on four particular areas of research that target abilities frequently identified as key facets of the human social repertoire - ToM, empathy, trait inference, and moral reasoning (Lieberman, 2007; Van Overwalle, 2009) – and, for each, we conducted separate meta-analyses of the available functional imaging data to determine the brain regions consistently implicated. In the case of trait inference, this was the first neuroimaging meta-analysis to include studies that used stimuli other than faces (see Section 2, and also Bzdok et al., 2011, and Mende-Siedecki et al., 2013, for contrasting approaches). In the other three cases, we performed updates of prior meta-analyses (Eres et al., 2018; Molenberghs et al., 2016; Timmers et al., 2018).

Further, we conducted an exploratory conjunction analysis aimed at identifying brain areas reliably implicated in all four social sub-domains and, thus, a core network for social cognitive processing (Bzdok et al., 2012; Schurz et al., 2020; Van Overwalle, 2009). We hypothesised that this core network would include parts of the MDN and the SCN. It is of note that, across all four social sub-domains, we took a different approach to study inclusion and exclusion criteria than that taken by some prior meta-analyses of general social cognition (e.g., Van Overwalle, 2009). In particular, we excluded studies investigating processes associated primarily with the self because social cognition is, although perhaps only in the strictest sense, about understanding other people. We also excluded studies in which tasks could be completed based on relatively simple perceptual processing and without a need for deeper cognitive and inferential processes (e.g., emotion discrimination tasks, automatic imitation). This was done in an attempt to constrain our inferences to be about the neurobiology underpinning cognitive rather than primarily perceptual social processes (for further detail on this distinction, see Adolphs, 2010, and Binney & Ramsey, 2020).

Finally, as a secondary aim, the present study used the meta-analytic approach to assess whether there are differences in the neural networks engaged by implicit and explicit social processing (also see Dricu & Frühholz, 2016; Eres et al., 2018; Fan et al., 2011; Molenberghs et al., 2016; Timmers et al., 2018). This was aimed at addressing a pervasive distinction in the social neuroscientific literature between automatic and controlled processes (Adolphs, 2010; Happé et al., 2017; Lieberman, 2007), and followed an assumption that implicit paradigms engage only automatic processes, whereas controlled processes are recruited during explicit paradigms (Sherman et al., 2014); automatic processes are described as unintentional, effortless, and fast, whereas controlled processes are deliberate, effortful, and thus slower (Lieberman, 2007; Shiffrin and Schneider, 1977). Some authors have argued that automatic and controlled social processes are mutually exclusive of one another and draw upon distinct cortical networks, with the former engaging lateral temporal cortex, the amygdala, ventromedial frontal cortex and the anterior cingulate, and the latter engaging lateral and medial prefrontal and parietal cortex (Forbes and Grafman, 2013; Lieberman, 2007). However, these dual-process models have been criticised for over-simplifying both the distinction and the relationship between automatic and controlled processes (Amodio, 2019; Cunningham and Zelazo, 2007; Ferguson et al., 2014; Fidler and Hütter, 2014; Fujita et al., 2014; Melnikoff and Bargh, 2018). An alternative proposal, that we describe above, makes a different distinction - one between representation and control. This neurocognitive model proposes that social processing relies on a single-route architecture wherein the degree to which cognitive processing has certain attributes (e.g., speed or effort) does not reflect one system versus another. Instead, it is proposed that it reflects the degree to which the control system needs to exert influence, upon otherwise automatic activation within the representational system, in order to meet the demands of a task in an appropriate and efficient manner (Binney and Ramsey, 2020; Jefferies, 2013). If the dual route model is correct, explicit but not implicit social paradigms should differentially engage brain regions associated with cognitive control demands, including the SCN and MDN. If the single-route model is correct, then there should be no qualitative difference in terms of the network of regions activated by implicit paradigms (ergo automatic processing) and explicit paradigms (ergo controlled processing), although there may be differences in the magnitude of regional activation.

To summarise, the aims of the present study were as follows: 1) explore the involvement of domain-general control systems in social cognition; more specifically, determine whether social cognitive processing reliably engages brain areas implicated in the controlled retrieval and selection of conceptual knowledge; and 2) examine the evidence for dual-route and single-route models of controlled social cognition.

## 2. Methods

*Preregistration and Open Science statement:* Following open science initiatives (Munafò et al., 2017), the current study was pre-registered via the Open Science Framework (OSF; osf.io/fktb8/). We adhered to our pre-registered protocols with a few minor exceptions (see Section S1 of Supplementary Information (SI) 1 for details). All the raw datasets are openly-available on the OSF project page and are accompanied by a range of study characteristics including details that are not the focus of the present study but may be of interest in future research (please see Section S1 of SI 1 for a detailed description). Moreover, the input data and output files of all analyses can be accessed via the OSF page.

In accordance with our pre-registered aims, we performed a comprehensive review of published functional neuroimaging studies investigating four social abilities – Theory of mind (ToM), trait inference, empathy and moral reasoning - and independent coordinate-based meta-analyses aimed at characterising the brain-wide neural networks underpinning each. In the case of three of these domains (ToM, empathy and moral reasoning), we updated earlier meta-analyses (Eres et al., 2018; Molenberghs et al., 2016; Timmers et al., 2018), capitalizing on additional data, and also implementing recommendations for best practice that became available in a year subsequent to these prior studies (Müller et al., 2018). In the case of trait inference, as far as we are aware, this was the first neuroimaging meta-analysis to include studies that explored potential sources of information beyond face stimuli (see Bzdok et al., 2011; Mende-siedlecki et al., 2013, for contrasting approaches). To localise the brain areas underpinning semantic retrieval and selection, we also updated a meta-analysis of functional imaging studies of semantic control by Noonan et al. (2013). This involved the inclusion of additional data, and improvements in meta-analytic tools which corrected previous implementation errors that led to the use of liberal statistical thresholds (Eickhoff et al., 2017).

To directly address our first aim, we assessed the degree of overlap between the neural networks supporting semantic control and those involved in social information processing via a set of formal conjunctions and contrasts analyses. To address our second aim, where possible, we contrasted brain-wide activation associated with explicit versus implicit social cognitive paradigms. Tasks that drew the participant’s attention to the behaviour/cognitive process of interest were categorised as explicit, while tasks that used non-specific instructions (e.g., they involved passive observation of stimuli) or employed orthogonal tasks (e.g., age judgement) were categorised as implicit. Finally, where sufficient relevant information was available, we explored the influence of task difficulty on patterns of brain activation.

All of the meta-analyses reported below were conducted following best-practice guidelines recommended by Müller et al. (2018). This, as well as several refinements to inclusion/exclusion criteria, contributed to methodological differences between the present meta-analyses and those prior meta-analyses upon which the ‘updates’ were based. A summary of similarities and differences is provided in Table S1 (SI1) and further details are given in the sections below.

### 2. 1. Literature Selection and Inclusion Criteria

#### 2. 1. 1. General Approach and Criteria

Where possible, relevant functional neuroimaging studies were initially identified based on their inclusion in a recent prior neuroimaging meta-analysis. These lists were supplemented via a search on the Web of Science (WoS) online database (www.webofknowledge.com) for original reports published in the years subsequent, and by searching through reference lists of said articles. Each WoS search used the terms [‘fMRI’ or ‘PET’], as well as terms uniquely chosen for a given cognitive domain (see Table 2).

**Table 2.** Terms used to search the Web of Science database for relevant articles.

| Cognitive domain | Search terms |
| --- | --- |
| Semantic control | 'semantic', 'comprehension', 'conceptual knowledge', 'selection', 'retrieval', 'inhibition', 'control', 'elaboration', 'fluency', 'ambiguity', 'metaphor', 'idiom' |
| ToM | 'theory of mind', 'ToM', 'mentalising', 'mentalizing' |
| Trait inference | 'social judgement', 'social evaluation', 'social attribution', 'trait inference', 'impression formation' |
| Empathy | 'empathy', plus 'empath*' - corresponding variations (e.g. 'empathic') |
| Moral cognition | 'morality', 'moral', 'moral decision making', 'moral emotion', 'harm', 'guilt' |

*N.b.*, For all five cognitive domains, the search followed the following format: [fMRI OR PET] AND [term1 OR term2 OR ... OR termX].

A general set of inclusion criteria applied to all our analyses were as follows:

1. Only studies that employed task-based fMRI or PET to obtain original data were included. Studies employing other techniques (e.g., EEG/MEG), meta-analyses and review articles were excluded.
2. Studies were only included if they reported whole-brain activation coordinates that were localised in one of two standardised spaces – Talairach (TAL) or Montreal Neurological Institute (MNI) – or these coordinates were made available on request (see Section 1 of SI1). Coordinates reported in TAL space were converted into MNI space using the Lancaster transform (tal2icbm transform (Lancaster et al., 2007) embedded within the GingerALE software (version 3.0.2; http://brainmap.org/ale). Studies exclusively reporting results from region-of-interest or small volume correction analyses were excluded because these types of analysis violate a key assumption of coordinate-based meta-analyses (Eickhoff et al., 2012; Müller et al., 2018).
3. Studies were only included if they reported activation coordinates that resulted from univariate contrasts clearly aimed at identifying the process of interest (e.g., ToM). We included contrasts between an experimental task and either a comparable active control task or a low-level baseline such as rest or passive fixation. Contrasts against low-level baselines were included in the primary analyses because they can reveal activity associated with domain-general cognitive processes that is subtracted out by contrasts between active conditions. This could include semantic processes that are common to both social and non-social tasks. However, contrasts against low-level baselines also yield activity associated with differences in perceptual stimulation and attentional demand. To address this caveat, we repeated the analyses whilst excluding this subset of contrasts. The results can be found on the project’s OSF page (osf.io/fktb8/). We excluded contrasts that make comparisons between components of the process of interest (e.g., affective vs. cognitive ToM; utilitarian vs. deontological moral judgements) because we were interested in the common, core processes that would be subtracted out by these contrasts (but see the following paragraph).
4. Multiple contrasts from a single group of participants (e.g., separate contrasts against one of two different baseline conditions) were included in a single meta-analysis as long as they independently met all other inclusion criteria for the primary analyses. This allowed maximum use of all available data and enabled us to evaluate the effect of using different types of baseline, for example (see above). However, it is important to adjust for this (Müller et al., 2018), and accordingly, we adopted an approach to controlling for within-group effects (Turkeltaub et al., 2012); specifically, sets of activation coordinates from different contrasts, but the same participant group, were pooled. This means that when we refer to the numbers of experiments, we have counted multiple contrasts from a single participant sample as one single experiment. This partially explains why the number of experiments in our analyses is lower than in those of some prior meta-analyses. However, in formal contrast analyses that compare different conditions (e.g., instructional cue, task difficulty), contrasts like these would be separated, and care was also taken to minimize the difference in the number of experiments on either side of the contrast. For example, if a study reported two contrasts – one implicit and one explicit - based on the same participant group, only the peaks from the implicit task would be included in the contrast/conjunction analyses if there were a greater number of explicit than implicit tasks overall (see Figure S8).
5. Only studies that tested healthy participants were included. Contrasts including clinical populations or pharmacological interventions were excluded.
6. Only research articles published in English were included.

#### 2.1.2. Theory of Mind

This meta-analysis was built upon that of Molenberghs et al. (2016) and only included studies that were specifically designed to identify the neural network underpinning ToM processes (i.e., they employed tasks involving inferences about the mental states of others, including their beliefs, intentions, and desires). Therefore, studies that looked at passive observation of actions, social understanding, mimicry or imitation were not included, unless tasks included a ToM component. Unlike Molenberghs et al., (2016), we excluded studies investigating irony comprehension (e.g., Wang et al., 2006) because ToM might not always be necessary to detect non-literal meaning in language (Ackerman, 1983; Bosco et al., 2018; Pexman, 2008) and studies that employed interactive games (e.g., Rilling et al., 2008). These latter studies are commonly designed to investigate the degree to which ToM is engaged under different task conditions rather than distinguish activation associated with ToM from that related to other processes. Moreover, unlike Molenberghs et al. (2016), we excluded studies that employed trait inference tasks as these were considered separately (see Section 2.1.3).

Molenberghs et al.’s (2016) search was inclusive of fMRI studies published prior to July 2014 and yielded 144 independent experiments (1789 peaks) contributing to their analysis. We performed a WoS search for further original fMRI and PET studies conducted between August 2014 and March 2020, and a search of PET studies published prior to July 2014. We then applied our inclusion criteria to both newly identified studies and those analysed by Molenberghs and colleagues (see Table S1 for further differences in criteria). In the end, we found 136 experiments with a total number of 2158 peaks and 3452 participants that met our criteria for inclusion (see Figure S1of SI1 for more details regarding the literature selection process; and Table S1 of SI2 for a full list of the included experiments).

#### 2.1.3. Trait inference

Studies were included in the meta-analysis if they employed tasks that required the participants to infer the personality traits of others based on prior person knowledge or another’s appearance and/or behaviour. Whereas the types of mental states typically inferred in ToM tasks are transitory in nature (e.g., relating to immediate goals or the intentions behind a specific instance of behaviour), traits are coherent and enduring dispositional characteristics of others (i.e., personality traits; Van Overwalle, 2009). Previous meta-analyses (Molenberghs et al., 2016; Schurz et al., 2014) of ToM have included tasks requiring trait inferences. However, it has been suggested that personality trait inferences differ from mental state inferences in terms of likelihood and speed of processing, and hold a higher position in the hierarchical organisation of social inferential processes (Korman and Malle, 2016; Malle and Holbrook, 2012). In line with this proposal, we maintained a distinction and performed separate analyses. Moreover, previous imaging meta-analyses of trait inference were limited to studies that used face stimuli (Bzdok et al., 2011; Mende-siedlecki et al., 2013). However, trait inferences can be made on the basis of many different sources of information, including physical appearance, behaviour and prior knowledge about others (Uleman et al., 2007). To our knowledge, the present attempt is the first to include studies that required participants to make trait inferences based on facial photographs, behavioural descriptions *or* prior person knowledge. We excluded any studies that asked participants to make inferences about transitory mental states, including basic emotions. We also excluded studies that did not use a subtraction approach, but rather investigated brain activity that varied parametrically with the levels of a pre-defined trait dimension (e.g. Engell et al., 2007). Finally, we excluded studies that included emotional face stimuli to avoid conflating brain activity related to trait inference with that associated with emotion recognition and processing.

We performed a WoS search of studies published before March 2020 and reference-tracing to identify relevant studies for inclusion in the meta-analysis. A total of 40 experiments with 523 peaks and 732 participants were found to meet the criteria for inclusion (Figure S2 - SI1; Table S2 - SI2).

#### 2.1.4. Empathy

This meta-analysis was built upon that of Timmers et al. (2018) and only included studies that were specifically designed to identify the neural network underpinning empathy by employing tasks asking participants to observe, imagine, share and/or evaluate the emotional or sensory state of others. The task definition was kept identical to previous meta-analyses on empathy (Fan et al., 2011; Timmers et al., 2018). We also made a distinction between tasks eliciting empathic responses to other people’s pain and those investigating empathic responses to others’ affective states.

Timmers et al. (2018) included studies published before December 2017, totalling 128 studies with 179 contrasts (1963 peaks). We identified additional original studies conducted between January 2018 and March 2020 via a WoS search and subsequently applied our inclusion criteria to all, including those analysed by Timmers et al. (2018) (see Table 1 for further differences in criteria). This resulted in a yield of 164 experiments with a total number of 2704 peaks and 4423 participants (Figure S3 - SI1; Table S3 – SI2). Empathy for pain was independently investigated in 93 of these experiments, empathy for affective states was independently explored in 70 experiments, and 9 experiments concurrently explored both empathy for pain and emotions in the same contrasts.

#### 2.1.5. Moral reasoning

This analysis updated a previous meta-analysis conducted by Eres et al., (2018) and included studies that employed tasks designed to investigate judgements and decision-making based on moral values. In line with Eres et al., (2018), studies that did not specifically have a morality component were not included. For example, studies investigating judgements regarding adherence to social expectations but not moral values (e.g., Bas-Hoogendam et al., 2017) were excluded.

Eres et al., (2018)’s search was restricted to fMRI studies and covered the period before February 2016 yielding 123 contrasts (989 peaks). We expanded this list via a WoS search for original fMRI and PET studies published between March 2016 and March 2020, and a search for PET studies published before March 2016, and then applied our inclusion criteria (see Table 1 for differences in criteria). This resulted in a yield of 69 experiments with a total number of 909 foci and 1609 participants (Figure S4 - SI1; Table S4 – SI2).

#### 2.1.6. Semantic Control

In this meta-analysis, we sought to extend an earlier meta-analysis conducted by Noonan et al. (2013). In line with theirs, this analysis only included studies that were specifically investigating semantic processing, and that reported contrasts that reflected high > low semantic control, or comparisons between a task requiring semantic control and an equally demanding executive decision in a non-semantic domain. We excluded studies with a focus upon priming without an explicit semantic judgment (e.g., primed lexical decision), bilingualism, episodic memory, or sleep consolidation.

Noonan et al., (2013)’s search covered the period between January 1994 and August 2009 and yielded 53 studies (395 peaks) that met their criteria for inclusion in their analysis. We performed a WoS search for original studies published between September 2009 and March 2020, and reference-tracing, and then applied our inclusion criteria to both newly identified studies and those analysed by Noonan et al. (2013). This produced a yield of 96 experiments with a total number of 981 peaks and 2052 participants that met the criteria for inclusion in our analysis (Figure S5 - SI1; Table S5 – SI2).

### 2.2. Data Analysis

We performed coordinate-based meta-analyses using the revised activation likelihood estimation (ALE) algorithm (Eickhoff et al., 2012, 2009; Turkeltaub et al., 2012) implemented in the GingerALE 3.0.2 software (http://brainmap.org/ale). We used the GingerALE software to conduct two types of analysis. The first were independent dataset analyses, which were used to identify areas of consistent activation across particular sets of experiments. These analyses were performed only on the experiment samples with a recommended minimum of 17 experiments in order to have sufficient power to detect consistent effects and circumvent the possibility of results being driven by single experiments (Eickhoff et al., 2016). The ALE meta-analytic method treats reported activation coordinates as the centre points of three-dimensional Gaussian probability distributions which take into account the sample size (Eickhoff et al., 2009). First, the spatial probability distributions of all coordinates reported were aggregated, creating a voxel-wise modelled activation (MA) map for each experiment. Then, the voxel-wise union across the MA maps of all included experiments was computed, resulting in an ALE map that quantifies the convergence of results across experiments (Turkeltaub et al., 2012).The version of GingerALE used in the present study tests for above-chance convergence between experiments (Eickhoff et al., 2012) thus permitting random-effects inferences.

Following the recommendations of Eickhoff et al. (2016), for the main statistical inferences, the individual ALE maps were thresholded using cluster-level family-wise error (FWE) correction of p < 0.05 with a prior cluster-forming threshold of p < 0.001. Cluster-level FWE correction has been shown to offer the best compromise between sensitivity to detect true convergence and spatial specificity (Eickhoff et al., 2016). This was complemented by a highly conservative voxel-level FWE correction of p < 0.05 which, despite the decreased sensitivity to true effects, allows the attribution of significance to each voxel above the threshold, offering increased spatial specificity (Eickhoff et al., 2016). The FWE-corrected cluster-level and voxel-height thresholds were estimated using a permutation approach with 5000 repetitions (Eickhoff et al., 2012). None of the meta-analyses that we updated had used the recommended cluster-level FWE or the FWE height-based correction methods.

The second set of analyses, conjunction and contrast analyses, were also performed in GingerALE and were aimed at identifying similarities and differences in neural activation between the different sets of studies. The conjunction images were generated using the voxel wise minimum value (Nichols et al., 2005) of the included ALE maps to highlight shared activation. Contrast images were created by directly subtracting one ALE map from the other to highlight unique neural activation associated with each dataset (Eickhoff et al., 2011). Then, the differences in ALE scores were compared to a null-distribution estimated via a permutation approach with 5000 repetitions. The contrast maps were thresholded using an uncorrected cluster-forming threshold of p < 0.001 and a minimum cluster size of 200 mm^3^.

In addition, we performed post-hoc analyses to investigate if the clusters of convergence revealed by the ALE analyses were driven by experiments featuring specific characteristics of interest (i.e., type of instructional cue, task difficulty). To this end, we examined the list of experiments that contributed at least one peak to each ALE cluster and compared the number of contributing experiments featuring the characteristic of interest (e.g., explicit vs implicit processing) by conducting Fisher’s exact tests of independence and post-hoc pairwise comparisons (using False Discovery Rate correction for multiple comparisons) in RStudio Version 1.2.5001 (RStudio Team, 2020).

A full list of the confirmatory and exploratory analyses we conducted can be found in Section 3 of SI1.

## 3. Results

### 3.1. The “Social Brain”

#### 3.1.1. Theory of Mind

Convergent activation across all 136 ToM experiments was found in 13 clusters (see Figure 1a and Table S1.1.1 – SI3) located within the bilateral middle temporal gyrus (MTG) (extending anteriorly towards the temporal poles and also in a posterior and superior direction towards the superior temporal gyrus (STG) and angular gyrus (AG) in both hemispheres), bilateral IFG, bilateral dorsal precentral gyrus, ventromedial prefrontal cortex (vmPFC), dorsomedial prefrontal cortex (dmPFC), pre-SMA, precuneus, left fusiform gyrus and left and right cerebellum. All these clusters survived both the height-based and extent-based thresholding. A cluster in the posterior cingulate cortex (PCC) survived height-based thresholding but did not survive extent-based thresholding. These results are largely consistent with those of Molenberghs et al. (2016), with the difference being that they did not find activation in SMA, left fusiform gyrus or cerebellum. In order to address concerns regarding the validity of some other popular ToM tasks (Heyes, 2014; Quesque and Rossetti, 2020), we conducted a separate supplementary meta-analysis that was limited to the subset of ToM experiments that employed false belief tasks (see Section 3.1 of SI1, Table S1.1.2). This analysis revealed convergent activation in similar temporo-parietal and medial frontal regions to the inclusive ToM analysis but did not implicate the lateral frontal cortex.

**Figure 1.**
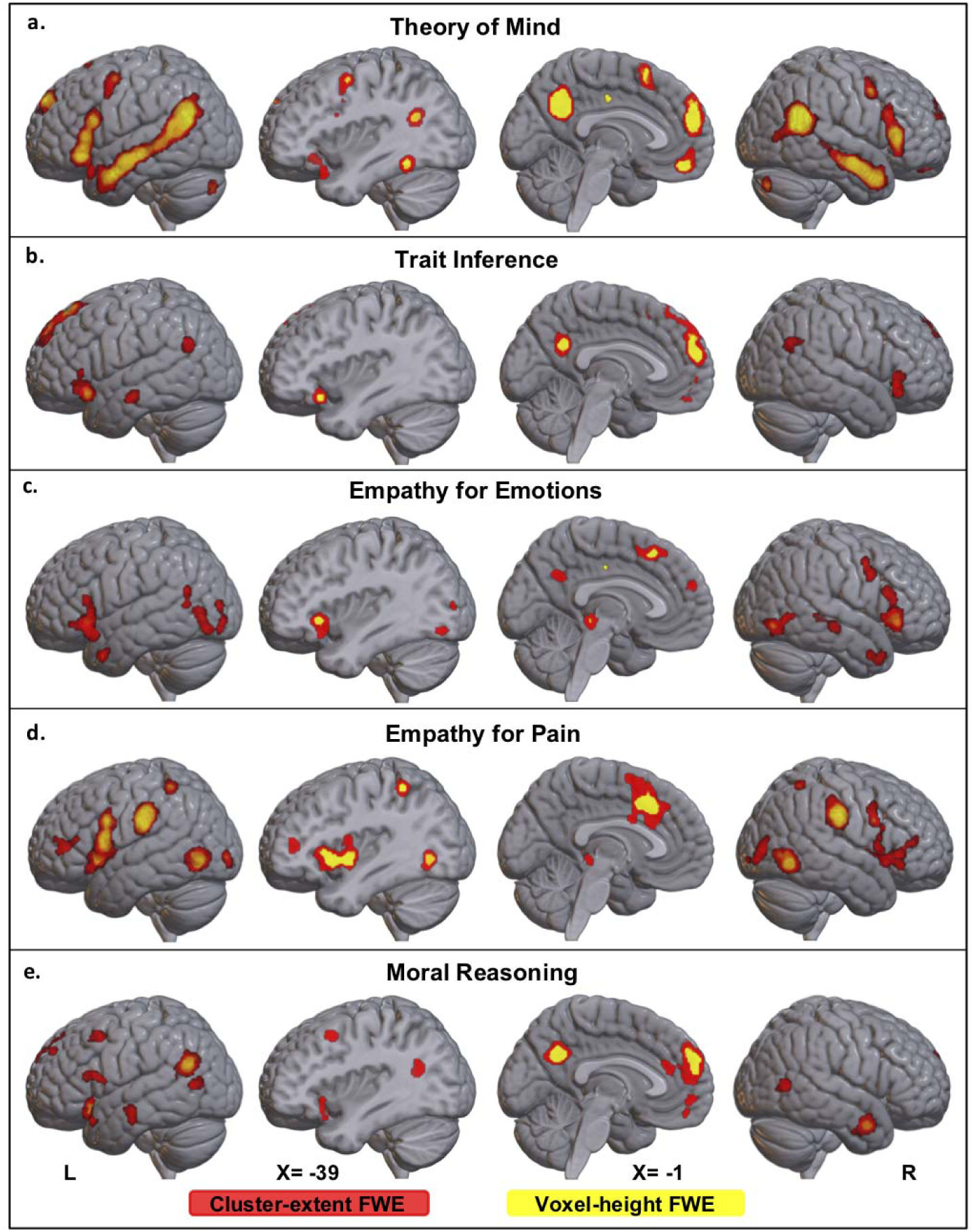
Binary whole-brain ALE maps showing statistically significant convergent activation resulting from independent meta-analyses of ToM studies (N=136), trait inference (N= 40), empathy for pain (N=80) and emotions (N=75) and moral reasoning (N=69). The ALE maps were thresholded using an FWE corrected cluster-extent at *p* < .05 with a cluster-forming threshold of *p* < .001 (red) and an FWE corrected voxel-height threshold of *p* < .05 (yellow). The lateral views, which show projections on the cortical surface, are accompanied by brain slices at the sagittal midline and also coplanar with the peak of the left IFG cluster observed across all social domains (X = -39; Table S1.5).

#### 3.1.2. Trait inference

The ALE meta-analysis revealed convergent activation across 40 experiments in 8 clusters (Figure 1b, Table S1.2) implicating the bilateral IFG, dmPFC, vmPFC, PCC, right pMTG (extending to AG), left AG and left anterior MTG. Voxels from all clusters, except for those in the right pMTG and vmPFC, survived the more conservative height-based thresholding.

#### 3.1.3. Empathy

The ALE meta-analysis of all 164 empathy experiments revealed 16 clusters of convergent activation (Figure S6a; Table S1.3.1), including in the bilateral IFG (extending towards the insula), SMA, dmPFC, bilateral posterior inferior temporal gyrus (ITG), right pMTG, bilateral supramarginal gyrus (SMG), left inferior parietal lobule (IPL), bilateral occipital cortex, left amygdala, left thalamus, left caudate and brainstem. These clusters survived both the height-based and extent-based thresholding, except for the anterior dmPFC, right pMTG and brainstem clusters, which survived extent-based thresholding only. Two clusters, one in the right cerebellum and one in the right hippocampus survived height-based thresholding but did not survive cluster extent-based thresholding. These areas were also implicated by Timmers et al. (2018). In contrast, however, we did not find convergent activation in the left posterior fusiform gyrus, left SMG (although we found a cluster slightly more posterior and inferior), left anterior ITG, right TP, precuneus, middle cingulate gyrus, right superior parietal lobule, and right amygdala.

The separate ALE maps for empathy for pain and empathy for affective states are displayed in Figure 1c and d. A conjunction analysis found activation common to empathy for pain (Table S1.3.2) and empathy for affective states (Table S1.3.3) in the bilateral insula (extending to the IFG), SMA, right precentral gyrus, right ITG, bilateral occipital cortex and the brainstem (Figure S6b; Table S1.3.4). Formal contrasts revealed that empathy for pain and empathy for emotions also engage highly distinct brain areas (Figure S6b; Table S1.3.4). Clusters with increased convergence for empathy for pain were found in left IFG (pars triangularis), right MFG, bilateral insula, middle cingulate gyrus, bilateral SMG, right IPL and bilateral pITG. In contrast, increased convergence in empathy for affective states was revealed in left IFG (pars orbitalis), PCC, left pMTG, right temporal pole and left anterior MTG. Given these significant differences in their underlying neural networks, empathy for pain and empathy for emotions were considered separately for all subsequent analyses.

#### 3.1.4. Moral reasoning

Convergent activation across all 69 experiments studying moral reasoning was found in 11 clusters (Figure 1e, Table S1.4) located in the left IFG, left insula (extending towards the superior temporal pole), mPFC, medial orbitofrontal cortex (OFC), precuneus, bilateral pMTG, and the bilateral anterior MTG. Only four clusters - left insula, mPFC, precuneus and left pMTG - survived height-based thresholding. These results are mostly consistent with those obtained by Eres (2018), with the difference that we did not find convergent activation in the left amygdala and right AG, and found additional clusters of convergent activation in left MFG, bilateral anterior MTG, and right pMTG.

#### 3.1.5. A common network for multiple sub-domains of social cognition

To identify brain areas consistently activated across multiple sub-domains of social cognition, we performed an overlay conjunction analysis of the cluster-extent FWE-corrected ALE maps associated with ToM, trait inference, empathy (for pain and/or emotions) and moral reasoning (see Figure 2a, Table S1.5). Convergent activation across all four socio-cognitive sub-domains was found in the left IFG (pars orbitalis), mPFC, precuneus and left pSTG. Overlapping areas of activation across three of four social sub-domains included right IFG, left IFG (pars triangularis and pars opercularis), SMA, medial OFC, left MTG, right anterior MTG and right pMTG. Overlap between two of four maps was found in right posterior IFG, bilateral precentral gyrus, right AG, right pMTG, left TP and left pMTG. Because the conservative thresholding used in this analysis could have excluded smaller clusters that nonetheless overlap across the sub-domains, we repeated the conjunction using ALE maps treated with a more liberal statistical threshold of p<.001 uncorrected. This revealed additional overlapping activation for all four social domains in the right IFG (pars orbitalis) and bilateral ATL (Figure S7). These brain areas have been implicated in a variety of social-cognitive abilities by multiple previous meta-analyses (Alcalá-López et al., 2018).

**Figure 2.**
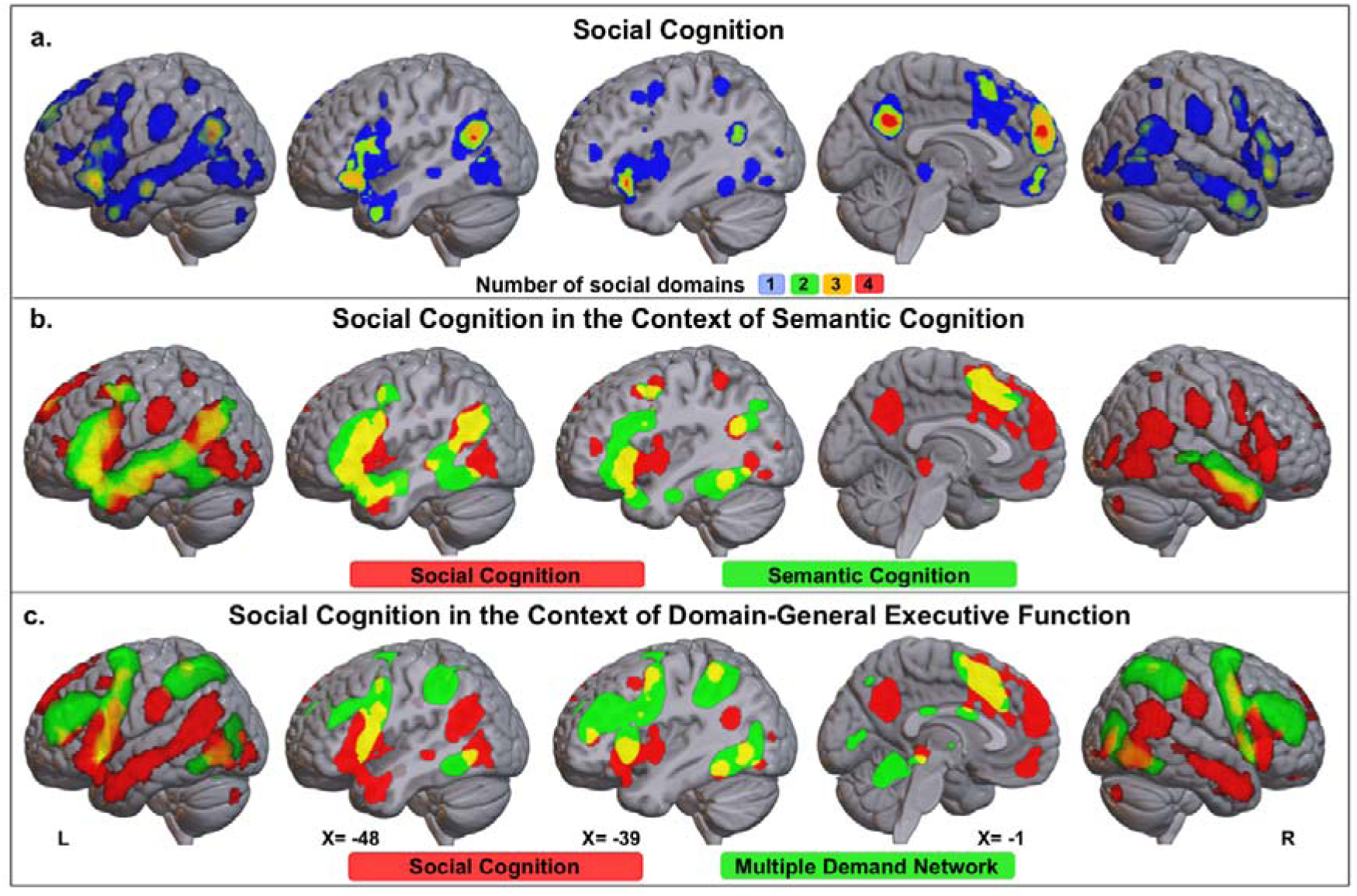
The neural network engaged in social cognitive processing: (a) An overlay conjunction of the ALE maps resulting from independent meta-analyses of ToM studies, trait inference, empathy for pain/emotions, and moral reasoning. The map displays the number of social domains showing convergent activation in each voxel. The ALE maps were thresholded using an FWE corrected cluster-extent threshold at *p* < .05 with a cluster-forming threshold of *p* < .001. (b) The binarized social cognition map (red) generated by the overlay conjunction is displayed overlaid with a binarized ALE map of convergent activation across N = 415 semantic > non-semantic contrasts generated in Jackson, 2021 (green); overlap is shown in yellow. (c) The binarized social cognition map (red) generated by the overlay conjunction is displayed overlaid with a mask of the multiple-demand network (MDN) generated in Federenko et al., 2013 (green) by contrasting hard > easy versions of seven diverse cognitive tasks; overlap is shown in yellow. The lateral views, which show projections on the cortical surface, are accompanied by brain slices at the sagittal midline and also coplanar with the peak of the left STG (X = -48) and left IFG (X = -39) clusters that overlapped across all four social domains (Table S1.5).

The extent to which brain regions engaged in social cognition overlap with those engaged in general semantic cognition (including both representation and control processes) is illustrated in Figure 2b. Figure 2c shows that the brain regions engaged in social cognition are largely non-overlapping with the network engaged by domain-general executive processes (i.e., the MDN).

### 3.2. The semantic control network

The ALE meta-analysis of all 96 semantic control experiments revealed convergent activation in a distributed network consisting of frontal, temporal and parietal areas (Figure 3a, Table S2). The largest cluster was located in the left frontal lobe and extended from the IFG (pars orbitalis) to MFG. In the right frontal lobe, convergent activation was limited to two clusters with peaks in pars orbitalis and pars triangularis of the IFG. Consistent activation was also found in the medial frontal cortex with the peak in SMA. The left temporal cluster extended from the posterior portion of the MTG, which showed the highest level of convergence, to the fusiform gyrus. All these clusters survived both the height-based and extent-based thresholding. In addition, two left IPL clusters survived only the cluster-extent FWE correction. In contrast to Noonan et al., (2013), we did not find convergent activation in ACC, bilateral SFG, left AG, right IPL/SPL, and left anterior MTG.

**Figure 3.**
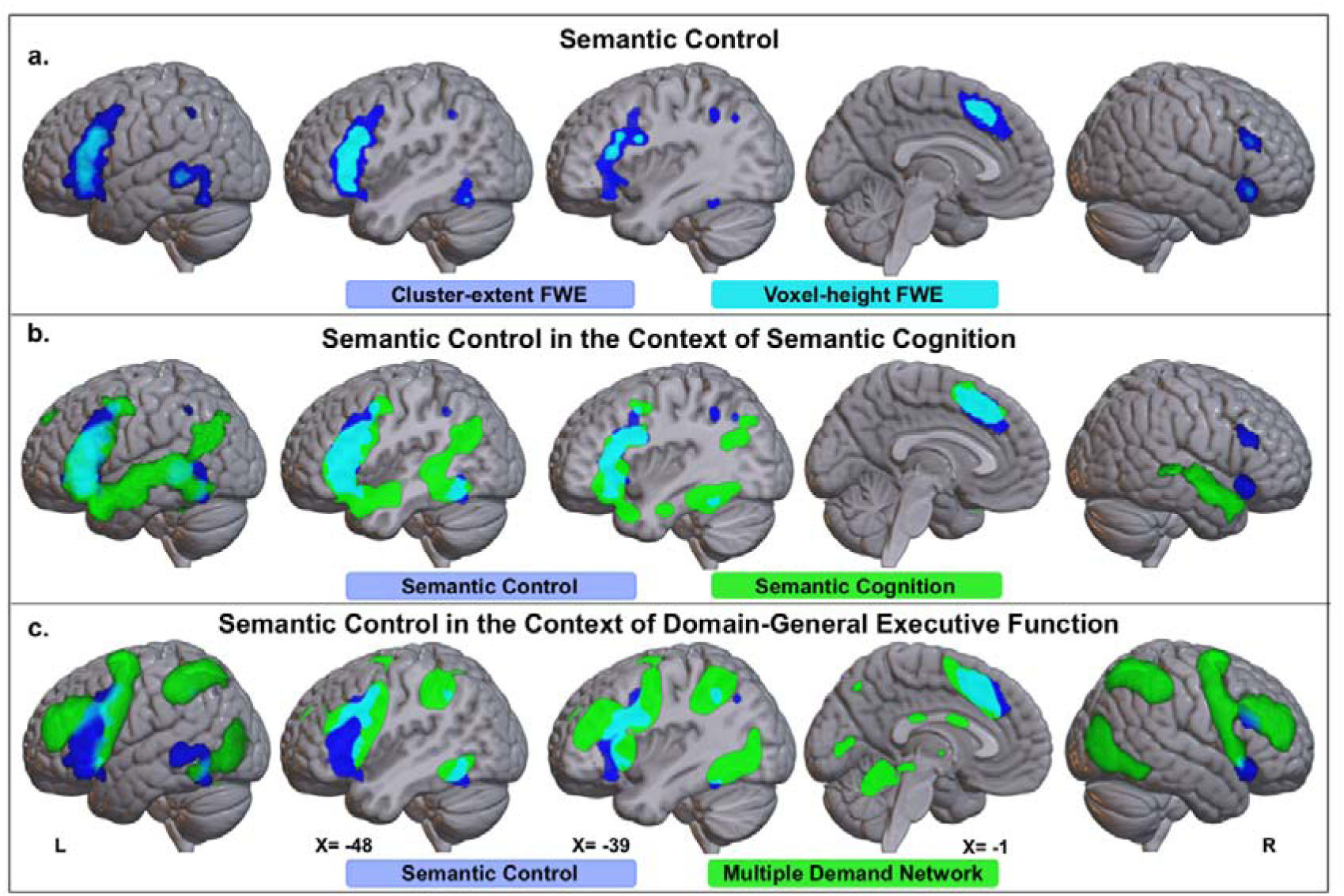
The neural network engaged in semantic control: (a) Binarized ALE maps showing statistically significant convergent activation across 96 experiments contrasting high > low semantic control thresholded using cluster-extent FWE correction of p < .05 with a cluster-forming threshold of p < .001 (blue) and voxel-height FWE correction of p < .05 (cyan). (b) The binarized semantic control map (blue) overlaid with a binarized ALE map of convergent activation across N = 415 semantic > non-semantic contrasts generated in Jackson, 2021 (green); overlap is shown in cyan. (c) The binarized semantic control map (blue) overlaid with a mask of the multiple-demand network (MDN) generated in Federenko et al., 2013 (green) by contrasting hard > easy versions of seven diverse cognitive tasks; overlap is shown in cyan. The lateral views, which show projections on the cortical surface, are accompanied by brain slices at the sagittal midline and also coplanar with the peak of the left STG (X = -48) and left IFG (X = -39) clusters that overlapped across all four social domains (Table S1.5).

Figure 3b illustrates the extent to which brain regions engaged in semantic control overlap with those engaged in general semantic cognition (including both representation and control processes), while Figure 3c illustrates their overlap with the network engaged by domain-general executive processes (i.e., the MDN).

### 3.3. Neural substrates shared by semantic control and social cognition

#### 3.3.1. ToM

Overlap between the neural network underpinning semantic control (i.e., SCN & regions of the MDN) and the ToM network was found in 8 clusters located in the left IFG (including pars orbitalis and triangularis and extending to the precentral gyrus) and, to a smaller extent, the right IFG, the left dorsal precentral gyrus, SMA, left pMTG, left superior temporal pole and the left fusiform gyrus (Figure 4a, Table S3.1.1). The results of the conjunction between semantic control and false belief reasoning can be found in Section 3.1 of SI1 and Table S3.1.2. This analysis revealed overlapping activation in the pMTG, but not in the SMA or lateral frontal cortex.

**Figure 4.**
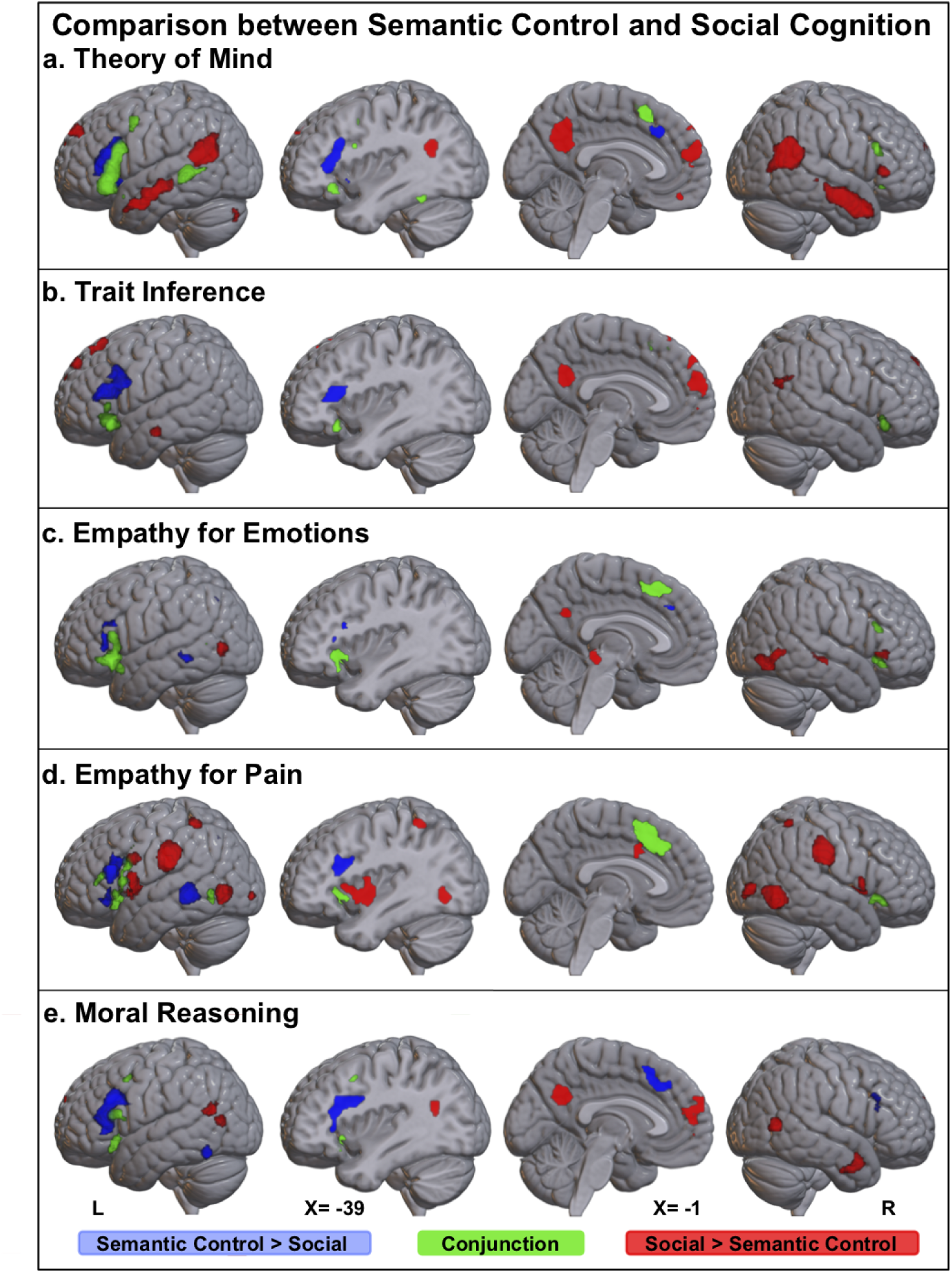
Results of the contrast (blue, red) and conjunction (green) analyses between the ALE maps associated with semantic control and each social domain: a) Theory of Mind b) Trait Inference c) Empathy for Emotions d) Empathy for Pain and e) Moral Reasoning. The contrast maps were thresholded with a cluster-forming threshold of *p* < .001 and a minimum cluster size of 200 mm^3^. The lateral views, which show projections on the cortical surface, are accompanied by brain slices at the sagittal midline and also coplanar with the peak of the left IFG cluster (X = -39) that overlapped across all four social domains (Table S1.5).

#### 3.3.2. Trait Inference

Brain areas involved in both semantic control and trait inference included bilateral IFG (pars orbitalis), SMA and dmPFC (Figure 4b, Table S3.2).

#### 3.3.3. Empathy for emotions

The neural network underpinning semantic control overlapped with the areas engaged in empathy for emotions in bilateral IFG (pars orbitalis and pars triangularis), SMA, left pMTG and right insula (Figure 4c, Table S3.3).

#### 3.3.4. Empathy for pain

Overlapping activation between empathy for pain and semantic control was revealed in left IFG (pars orbitalis and pars triangularis), right IFG (pars orbitalis), left precentral gyrus, bilateral insula, SMA and left posterior ITG (extending towards pMTG) (Figure 4d, Table S3.4).

#### 3.3.5. Moral reasoning

Overlapping activation in response to semantic control and moral reasoning included left insula (extending to pars orbitalis of the IFG), pars opercularis of the left IFG and the left precentral gyrus (Figure 4e, Table S3.5).

Overall, the neural network engaged in semantic control overlapped with the neural networks underpinning all four social domains in the left IFG and, in particular, pars orbitalis. Except for moral reasoning, overlapping activation was also found in the right IFG (pars orbitalis) and SMA. In the left pMTG, we found a large area of overlap between semantic control and ToM and some evidence of overlap between semantic control and empathic processing.

### 3.4. Explicit versus implicit social cognition

Further to the meta-analyses above, we compared activation associated with implicit and explicit paradigms for studying empathy for emotions, empathy for pain and moral reasoning. The results of independent analyses are displayed in Figure 5 a-c and Tables S4.1.1 – S4.1.6). Conjunctions and formal contrasts are displayed in Figure 5 d-f and Tables S4.2.1 – S4.2.3). The only difference between activation associated with explicit and implicit paradigms, as identified by these formal comparisons, was in the case of empathy, with a small cluster in the dmPFC showing increased convergence for explicit as compared to implicit empathy (see Section 3.3.1. of Supplementary Information). In addition, we conducted exploratory cluster analyses to investigate whether the explicit and implicit experiments contributed similarly to each of the significant ALE clusters found for each social domain. In summary, these analyses (Figure S8) revealed that in the case of all social domains, implicit and explicit experiments contributed equally to most clusters (see Section 3.3.2. of Supplementary Information for a more detailed description).

**Figure 5.**
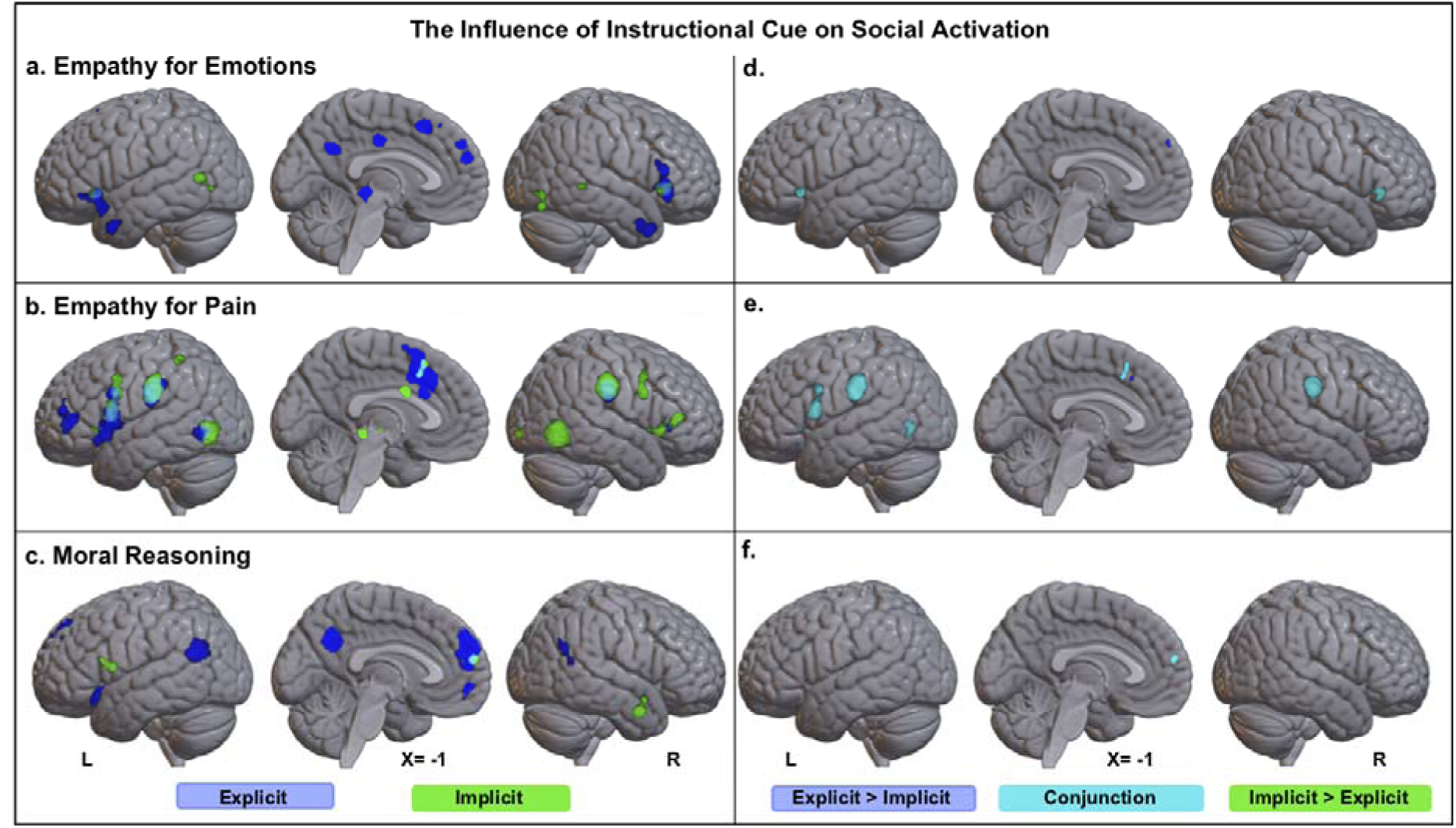
The left panel displays the binary ALE maps showing statistically significant convergent activation resulting from independent meta-analyses on explicit (blue) and implicit (green) studies on a) Empathy for Emotions, b) Empathy for Pain and c) Moral Reasoning. The ALE maps were thresholded using an FWE corrected cluster-extent threshold of *p* < .05 with a cluster-forming threshold of *p* < .001. The right panel displays the results of the contrast (dark blue, green) and conjunction (cyan) analyses between the ALE maps associated with explicit and implicit instructions. The contrast maps were thresholded at *p* < .001 and using a minimum cluster size of 200 mm^3^. The lateral views, which show projections on the cortical surface, are accompanied by brain slices at the sagittal midline.

### 3.5. The relationship between cognitive effort and brain regions engaged during social cognitive tasks

The above-reported conjunction analyses suggest that social cognition engages regions associated with semantic control. In these analyses, we took a pooled approach which involved collapsing over many different comparisons between social and non-social tasks and ignoring subtler differences between experimental and baseline conditions. The key advantage of this approach is that it identifies activation that is generalisable across highly variable experimental conditions. However, ignoring experimental differences precludes a determination of more specific factors driving a given region’s involvement. In particular, it is not possible to directly infer from the above results that semantic control regions are specifically being engaged by the cognitive control demands of social tasks. Therefore, to address this issue, we performed a set of exploratory analyses to determine whether the IFG and pMTG regions are sensitive to the degree of cognitive effort required to complete social tasks. While these analyses cannot disentangle semantic control from other forms of control, they represent a further initial step towards confirming a role of semantic control regions in social regulatory processes. To this end, we took experiments that used explicit paradigms and, on the basis of reported inferential statistics regarding participants’ reaction/decision times, categorised them according to whether the experimental condition was more difficult than the control condition (E>C), experimental and control conditions (E=C) were equally difficult, or the experimental condition was easier than the control condition (C>E). In the subsequent set of analyses we worked with the premise that in the case of E=C experiments and C>E experiments, activation associated with cognitive effort that is common to both the experimental and control conditions is subtracted away (along with activation specific to the control condition). In contrast, E>C experiments preserve activation associated with cognitive effort that is specific to the experimental condition. Therefore, a contrast analysis pitting E>C experiments against either C>E or E=C experiments will reveal activation associated with cognitive effort specific to the social domain. A conjunction will reveal activation associated with social processing irrespective of task difficulty.

There was only enough information regarding behavioural data to allow for sufficiently powered analyses in the case of ToM (Figure S9) where there were 26 E>C ToM experiments and 25 E=C ToM experiments. The results of the independent ALE analyses are reported in Tables S5.1 – S5.3. A conjunction analysis of E>C and E=C experiments yielded common activation in the left IFG (pars orbitalis and pars triangularis), dmPFC, precuneus, bilateral anterior MTG, right pMTG and left SMG (cyan in Figure 6a; Table S5.3) which we interpret as regions engaged in ToM irrespective of task difficulty. Interestingly, a contrast of E>C with E=C ToM experiments revealed differential activation in the left pMTG, an area implicated in semantic control. The full reports of these analyses, including prerequisite independent ALE analyses on the E>C ToM and E=C ToM experiments, can be found in Tables S5.1 – S5.4. For completeness, we also analysed C>E ToM experiments, but the sample size (N=14) was smaller than required to be sufficiently powered (Eickhoff et al., 2016) and therefore the result should be interpreted with caution (Figure 6a, Table S5.4).

**Figure 6.**
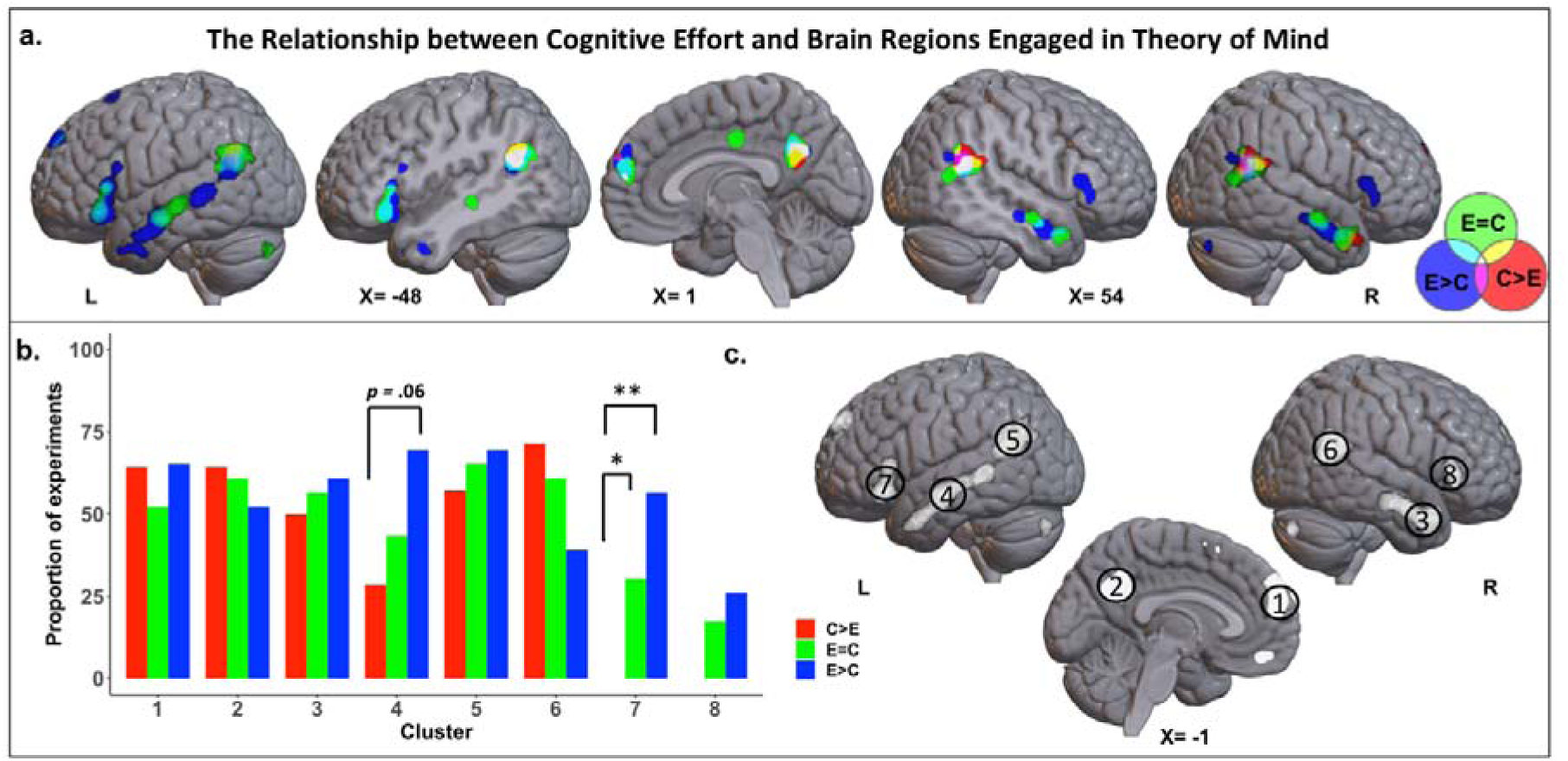
Results of exploratory analyses investigating the effect of task difficulty on ToM activation: (a) Binary ALE maps showing statistically significant convergent activation resulting from independent meta-analyses of three subsets of explicit ToM studies characterized by experimental conditions that were harder than the control task (E>C; N=26; blue), experimental and control conditions that were equally difficult (E=C; N=27; green) and control conditions that were harder than the experimental condition (C>E; N=14; red) as indexed by participant reaction times. The ALE maps were thresholded using an FWE corrected cluster-extent threshold at *p* < .05 with a cluster-forming threshold of *p* < .001. The lateral views, which show projections on the cortical surface, are accompanied by brain slices at the sagittal midline and also coplanar with the peak of the left IFG cluster (X = -39) that overlapped across all four social domains (Table S1.5) and the right pSTG cluster from the ToM meta-analysis (Table S1.1.1). (b) The results of the cluster analyses where bars represent the proportion of experiments in each difficulty category contributing to clusters of interest resulting from an ALE analysis of N = 60 ToM meta-analysis which included E>C, E=C and C>E experiments. (c) Binary ALE map showing statistically significant convergent activation across ToM experiments limited to those for which the behavioural information was known – this map represented the basis of the cluster analysis. The ALE map was thresholded using an FWE corrected cluster-extent threshold at *p* < .05 with a cluster-forming threshold of *p* < .001; ** *p* < .001 * *p* < .05.

Secondly, we conducted exploratory analyses to assess whether E>C, E=C or C>E ToM experiments were equally likely to contribute to each activation cluster (Figure 6b). The clusters were identified in an independent ALE analysis of ToM experiments limited to those for which the behavioural information was known (Figure 6c; Table S5.5). We expected clusters within brain areas that have a cognitive control function to have a disproportionate contribution from experiments in which the experimental task was more difficult than the control condition. To assess this, we conducted Fisher’s exact tests and then interrogated significant main effects through post-hoc pairwise comparisons and using false-discovery-rate adjustments for multiple comparisons. This cluster analysis revealed that E>C, E=C and C>E experiments contributed equally to mPFC (*p* = 0.67), precuneus (*p* = 0.8), right anterior MTG (*p* = 0.85), left pMTG (*p* = 0.74), right pMTG (*p* = 0.15) and right IFG (*p* = 0.15). Contributions to the left IFG cluster depended on the difficulty category (*p* < .001) and pairwise comparisons indicated that the C>E experiments contributed significantly less peaks compared to E>C (*p* = .001) and E>C (*p* = .046) experiments. Contributions to the left anterior MTG cluster also depended on the difficulty category (*p* = .043) and pairwise comparisons indicated that the C>E experiments contributed fewer peaks compared to E>C, but this effect did not survive correction for multiple comparisons (*p* = .06). These results suggest that the left IFG is particularly sensitive to cognitively-challenging ToM processing.

## 4. Discussion

Although many contemporary theories of social cognition acknowledge the importance of control, or regulatory processes (Adolphs, 2010; Amodio and Cikara, 2021; Frith and Frith, 2012), many key questions remain about their exact nature and neural underpinnings. These include a) whether multiple forms of cognitive control contribute to social cognition, b) whether these mechanisms are best understood in terms of domain-general processes or systems specialised for social information processing and, c) whether they are ubiquitously involved or selectively engaged according to certain task demands (Binney and Ramsey, 2020). In the present study, we set out to specifically investigate whether brain regions implicated in the controlled retrieval and selection of conceptual knowledge - namely the IFG and pMTG comprising the SCN (Jefferies, 2013; Lambon Ralph et al., 2017) - contribute to social processing. We simultaneously applied this question to multiple sub-domains of social cognition so that we could assess the extent to which involvement is general, or specific to certain types of social tasks and/or abilities. And we adopted a formal meta-analytic approach to extracting reliable trends from across a large number of functional neuroimaging studies and overcome the limitations of individual experiments (Cumming, 2014; Eickhoff et al., 2012). We found that theory of mind, trait inference, empathy, and moral reasoning commonly engage a core social network that includes the left IFG, left pMTG/AG, mPFC and precuneus. Moreover, the left IFG (particularly the pars orbitalis) region greatly overlapped with that implicated in an independent meta-analysis of neuroimaging studies of semantic control. Further, exploratory analyses suggest that both the left anterior IFG and the left posterior MTG (at a position just anterior to the ‘temporoparietal junction’) are sensitive to executive demands of social tasks. We interpret our overall findings as supportive of the hypothesis that the SCN supports social cognition via a process of controlled retrieval of conceptual knowledge. This aligns with a broader proposal in which social cognition is described as a flavour of domain-general semantic cognition and relies on the same basic cognitive and brain systems (Binney & Ramsey, 2020).

### 4.1. Cognitive control in social cognition

#### 4.1.1. The contribution of semantic control

A form of cognitive control known as semantic control could be crucial for effective goal-directed social behaviour (Binney and Ramsey, 2020; Satpute et al., 2014). In a broad sense, semantic control refers to a set of executive processes involved in the attribution of meaning to stimuli and experiences, and in the production of meaning-imbued behaviour (Corbett et al., 2015; Lambon Ralph et al., 2017). However, a key distinction has been drawn between a) top-down goal-directed retrieval and b) post-retrieval selection of goal-relevant semantic knowledge (Badre et al., 2005; Jefferies, 2013; Thompson-Schill et al., 1997), and it has been suggested that both of these two semantic control mechanisms contribute significantly to interpersonal interactions (Binney and Ramsey, 2020; Satpute and Lieberman, 2006). Studies of semantic cognition suggest that ‘selection’ is engaged when bottom-up, automatic activation of conceptual knowledge results in multiple competing semantic representations and/or responses. Social interactions frequently involve subtle or ambiguous cues, such as neutral facial expressions and bodily gestures, and/or conflicting cues (e.g., sarcasm). This causes semantic competition that can only be resolved by taking into account the wider situational and linguistic context and/or prior knowledge about the speaker (Aviezer et al., 2008; Pexman, 2008). Controlled retrieval processes, on the other hand, are engaged when automatic semantic retrieval fails to activate the semantic information necessary for the task at hand. This may occur frequently in social interactions, and particularly with less familiar persons, because of a preponderance of surface features (e.g., physical characteristics) over less salient features (e.g., personality traits, preferences, and mental states). To avoid exchanges that are deemed superficial at best, controlled retrieval must be used to bring to the fore person-specific but also context-relevant semantic information.

There is now over a decade’s worth of multi-method evidence that semantic control is underpinned by the left IFG and the left pMTG (Jefferies, 2013; Lambon Ralph et al., 2017). Research is now aimed at understanding the neural mechanisms by which these regions modulate semantic processing. One recent proposal is that it involves coordination of spreading activation across the semantic representational system (Chiou et al., 2018). According to the hub-and-spoke theory of semantic representation (Lambon Ralph et al., 2017), coherent concepts are represented conjointly by a central supramodal semantic ‘hub’ located in the ATLs, as well as multiple distributed areas of association cortex (i.e. ‘spokes’) that represent modality-specific information (e.g. visual features, auditory features, verbal labels, etc). Chiou et al., (2018) showed that the left IFG could be imposing cognitive control by flexibly changing its effective connectivity with the hub and spoke regions according to task characteristics; the IFG displayed enhanced functional connectivity with the ‘spoke’ region that processes the most task-relevant information modality. A similar proposal has been made for the contribution of domain-general cognitive control systems to social information processing. Zaki et al. (2010) found that, in the presence of conflicting social cues, IFG activity becomes functionally coupled with the brain areas associated with processing the particular cue type the participant chose to rely on to make inferences about emotional states. On this basis, they proposed that cognitive control areas upregulate activation within systems that represent social cues that are currently most relevant to the task. Consistent with this, a further study found evidence to suggest that the left IFG downregulates neural activation associated with task-irrelevant self-referential information when the task required reference to others (and vice versa) (Soch et al., 2017).

An important feature of semantic processing is the ability to accommodate new information that emerges over extended periods of time and update our internal representation of the current “state of affairs” in the external world according to contextual changes. This is particularly important for navigating social dynamics which are liable to abrupt and sometimes extreme changes in tone. For instance, imagine being in a bar and having your attention drawn to someone standing suddenly and picking up a glass. One might reasonably infer that this person is thirsty. That is until they proceed to walk towards a group of noisy sports fans rather than the bartender. In this case, you will likely adapt your interpretation and engage in a pre-emptive defensive stance. Recent research suggests that this ability to update depends, at least in part, on the IFG, as well as the mPFC and ventral IPL (also see Section 4.2.2) (Branzi et al., 2020). Likewise, Lavoie et al., (2016) showed that, during a ToM task, activation of the left IFG and pMTG is associated with contextual adjustments of mental state inferences (and also more general physical inferences) although not the representation of mental states specifically. Left IFG activation has also been observed when newly-presented information requires one to update the initial impression formed of another person (e.g., Mende-Siedlecki et al., 2013b, 2013a; Mende-Siedlecki and Todorov, 2016).

#### 4.1.2. The wider contribution of executive processes

According to Lambon Ralph, Jefferies, and colleagues, the executive component of semantic cognition comprises both semantic control and other domain-general processes (Lambon Ralph et al., 2017; Binney & Ramsey, 2020). The latter includes top-down attentional control and working memory systems that support goal-driven behaviour irrespective of the task domain (i.e., perceptual, motor or semantic). These processes recruit nodes of the MDN (Duncan, 2010), which include the precentral gyrus, MFG, IPS, insular cortex, pre-SMA and adjacent cingulate cortex (Assem et al., 2020; Fedorenko et al., 2013). In terms of organisation, the SCN appears to be nested among domain-general executive systems (Wang et al., 2020) and could play a role in mediating interactions between the MDN and the semantic representational system (Davey et al., 2016; Lambon Ralph et al., 2017). In line with this general perspective, we expected MDN regions to be reliably engaged by all four social sub-domains explored in the present meta-analyses. While there was evidence of engagement of the MFG, the pre-SMA, ACC, insula and IPS in some of the social sub-domains, MDN regions were not part of the core social processing network identified by the overlay conjunction analysis. This could reflect the fact that the majority of contrasts included in our meta-analyses employed high-level control conditions that were well-matched to the experimental conditions in terms of general task requirements, and thus, most activation associated with general cognitive demands had been subtracted away. Consistent with this notion is the fact that studies contrasting social tasks with lower-level control conditions (e.g., passive fixation) find more extensive MDN activation in ToM (Mason et al., 2008; Mier et al., 2010), trait inference (Chen et al., 2010; Hall et al., 2012), empathy (De Greck et al., 2012; Tamm et al., 2017) and moral reasoning (Reniers et al., 2012). The role of the MDN in social cognition is otherwise becoming well-established, and it has been found to be sensitive to difficulty manipulations in social tasks, showing increased activation in response to conflicting social cues (Cassidy and Gutchess, 2015; Mitchell, 2013), social stimuli that violate expectations (Cloutier et al., 2011; Hehman et al., 2014; Ma et al., 2012; Weissman et al., 2008) and increasing social working memory load (Meyer et al., 2012).

Finally, it is important to note that, although both MDN and the SCN co-activate in social and semantic tasks, the nature of their specific contributions *and* their anatomy are at least partially dissociable. The MDN is associated with the implementation of top-down constraints to facilitate goal-driven aspects of processing that is not limited to the semantic domain (Duncan, 2013; Fedorenko et al., 2013; Gao et al., 2020; Whitney et al., 2012). In contrast, the engagement of the anterior ventrolateral IFG (pars orbitalis) and the left pMTG appear specific to the semantic domain and, in particular, controlled semantic retrieval (Badre and Wagner, 2007; Gao et al., 2020; Hodgson et al., 2021; Whitney et al., 2012). Unlike the MDN, they do not appear to respond to challenging non-semantic tasks (Gao et al., 2020; Noonan et al., 2013; Whitney et al., 2012). Further, tasks associated with low conceptual retrieval demands but a requirement for response inhibition engage the MDN but do not engage the SCN, even if conceptual knowledge is used to guide responses (Alam et al., 2018).

#### 4.1.3. Double-route vs single-route cognitive architecture of social cognition

A secondary aim of the present study was to address a pervasive distinction in the social neuroscientific literature between automatic and controlled processes (Adolphs, 2010; Happé et al., 2017; Lieberman, 2007). Some authors have argued that automatic and controlled social processes are mutually exclusive of one another and draw upon distinct cortical networks (Forbes & Grafman, 2013; Lieberman, 2007; Satpute & Lieberman, 2006). The alternative is a single-route architecture where the degree to which behaviours have particular attributes (e.g. speed, effort, intentionality) does not reflect the involvement of one system and not another, but quantitative differences in the extent to which the control system interacts with the representational system in order to produce context-/task- appropriate responses (Binney and Ramsey, 2020). Our results are consistent with the latter perspective. The brain regions reliably activated in response to explicit instructions and those associated with implicit instructions revealed more overlap than discrepancy across empathy and moral reasoning tasks. Notably, this overlap included brain areas associated with executive functions: the bilateral IFG in the case of empathy for emotions and bilateral IFG and dmPFC in the case of empathy for pain. Moreover, cluster analyses of the ALE maps associated with the four social domains suggest that studies using explicit and implicit paradigms (which are assumed to engage controlled and automatic processing respectively) contributed equally to most activation clusters, including those in brain regions associated with control processes. Contrary to the predictions of dual-process models, these findings suggest that common neural networks contribute to both explicit and implicit social processing (also see Van Overwalle & Vandekerckhove, 2013). Furthermore, exploratory analyses suggest that both the left anterior IFG and the pMTG are sensitive to executive demands of social tasks. Overall, we argue that these results support the existence of a single-route cognitive architecture wherein the contribution made by control mechanisms to implicit and explicit social processing reflects cognitive effort demanded by the task at hand. This follows similar proposals put forth specifically in the domain of ToM (Carruthers, 2017, 2016).

### 4.2. Beyond cognitive control

Our findings converged upon four further regions that have been strongly linked with key roles in social cognition: the mPFC (including the anterior cingulate), the precuneus, the ‘temporoparietal junction’ (TPJ), and the ATL. We briefly discuss the putative role of each of these regions below.

#### 4.2.1. The ‘Temporo-parietal Junction’

A region often referred to as the ‘temporo-parietal junction’ (TPJ) has been subject to an elevated status within the social neurosciences. In particular, the right TPJ has been attributed with a key role in representing the mental states of others (Saxe and Wexler, 2005). In line with previous meta-analyses (Bzdok et al., 2012; Molenberghs et al., 2016; Schurz et al., 2020, 2014, 2013), our results reveal a bilateral TPJ region that is reliably involved in ToM tasks. In the left hemisphere, an overlapping area is also implicated in trait inference, empathy for emotions and moral reasoning which is suggestive of a broader role of the left TPJ in social cognition. In contrast, the right TPJ showed more limited overlap, being reliably engaged only by ToM and trait inference tasks, which is suggestive of a more selective role of the right TPJ in social cognition.

The TPJ encompasses a large area of cortex that is poorly defined anatomically and seems to include parts of the AG, SMG, STG and MTG (Schurz et al., 2017). Moreover, this area is functionally heterogeneous and has been associated with a variety of cognitive domains including but not limited to attention, language, numerosity, episodic memory, semantic cognition and social perception (Binder et al., 2009; Decety and Lamm, 2007; Deen et al., 2015; Humphreys and Lambon Ralph, 2015a; Igelström and Graziano, 2017; Özdem et al., 2017; Quadflieg and Koldewyn, 2017). While there is some indication that the function of the TPJ may be dependent on the hemisphere (e.g., Numssen et al., 2021), many cognitive domains, including ToM, are associated with bilateral TPJ activation. Our results at least seem to suggest dissociable roles of pMTG and a more posterior TPJ region; while the left pMTG is activated within both semantic control and ToM studies, a separate and more posterior STG (TPJ) area located closer to SMG/AG was reliably engaged by all of the social tasks, but not studies of semantic control. Furthermore, the results suggest that the left pMTG is sensitive to the difficulty of ToM tasks while the bilateral pSTG (TPJ) region is not.

This finding is generally in line with previous research suggesting a functional dissociation between the left pMTG and the left ventral IPL/AG regions. From one perspective, the activation of both regions appears to be positively associated with semantic tasks (Binder et al., 2009). However, the left pMTG shows increased activation to difficult relative to easier semantic tasks (Jackson, 2021; Noonan et al., 2013), unlike the ventral IPL/AG which has been shown to deactivate to semantic tasks when they are contrasted against passive/resting conditions where there may be greater opportunity for spontaneous semantic processing or ‘mind-wandering’ (Humphreys et al., 2015; Humphreys and Lambon Ralph, 2015b). Moreover, Davey et al., (2015) found that TMS applied to pMTG disrupted processing of weak semantic associations more than for strong associations, whereas TMS applied to AG had the opposite effect. Based on these and similar observations it has been suggested that the ventral IPL/AG has a role in the automatic retrieval of semantic information.

#### 4.2.2. The Default Mode Network

The pSTG/AG and the mPFC and precuneus regions we identified as part of the core social cognition network are also considered part of the default-mode network (DMN) (Buckner et al., 2008; Spreng and Andrews-Hanna, 2015). The DMN is a resting-state network, meaning that it is a group of regions consistently co-activated without the requirement of an explicit task. It is proposed that it is ideally suited for supporting self-generated internally-oriented, as opposed to externally-oriented, cognition (i.e., it is decoupled from sensory processing; Margulies et al., 2016; Smallwood et al., 2013). Some of these regions (e.g., the AG and mPFC) have been also implicated in processes that allow the integration of information over time (Huey et al., 2006; Humphreys et al., 2020; Ramanan et al., 2018; Ramanan and Bellana, 2019). These purported functions are all presumably important for social and more general semantic processing (see Section 4.1.1.) and likely involve domain-general mechanisms (also see Van Overwalle, 2009). However, the degree to which regions implicated in the DMN and those implicated in social and/or semantic cognition do or do not overlap is contentious and much is left to be gleaned regarding the relationship between these systems (Jackson et al., 2021, 2019; Mars et al., 2012).

#### 4.2.3. The anterior temporal lobe

Our findings implicate the lateral anterior temporal lobe (ATL), and particularly the dorsolateral STG/temporal pole (BA 38) and middle anterior MTG/STS, in all the socio-cognitive domains investigated, except for empathy for pain. Exploratory cluster analyses revealed that ATL engagement is not dependent on instructional cue or task difficulty, and thus it appears to serve a role other than control.

A key contribution of the ATL to social-affective behaviour has been recognised by comparative and behavioural neurologists for well over a century, owed at first to the acclaimed work of Brown and Schafer (1888) and, later, Klüver and Bucy, (1939) who provide detailed reports of profound social and affective disturbances in non-human primates following a bilateral, full depth ATL resection. These observations are mirrored in descriptions of neurogenerative patients that associate progressive ATL damage with a wide range of socio-affective deficits (Binney et al., 2016; Chan et al., 2009; Ding et al., 2020; Perry et al., 2001), including impaired emotion recognition (Lindquist et al., 2014; Rosen et al., 2004) and empathy (Rankin et al., 2005), impaired capacity for ToM (Duval et al., 2012; Irish et al., 2014), and a loss of person-specific knowledge (Gefen et al., 2013; Snowden et al., 2012, 2004). Over the past 10 years, there been a growing acceptance of the central role played by the ATL within the social neurosciences (Olson et al., 2013) and it now features prominently in some neurobiological models of face processing (Collins & Olson, 2014), ToM (Frith & Frith, 2006), moral cognition (Moll et al., 2005), and emotion processing (Lindquist et al., 2012). It has also been pinpointed as a key source of top-down influence on social perception (Freeman & Johnson, 2016). One influential account of social ATL function proposes a domain-specific role in the representation of social knowledge, including person knowledge, and other more abstract social concepts (Olson et al., 2013; Thompson et al., 2003; Zahn et al., 2007a).

A parallel line of research focused upon general semantic cognition has given rise to an alternative, more domain-general account of ATL function; there is a large body of convergent multi-method evidence from patient and neurotypical populations in support of a role of the ATL in the formation and storage of all manner of conceptual-level knowledge (Lambon Ralph et al., 2017). Research efforts have therefore recently begun to ask whether the purported roles of the ATL in both social and semantic processes can be reconciled under a single unifying framework (Binney et al., 2016; Rice et al., 2018). Some clues already exist in the aforementioned work of Klüver and Bucy (1939), who observed a broader symptom complex comprising multimodal semantic impairments, including visual and auditory associative agnosias, that might explain rather than just co-present with social-affective disturbances. More recent work that leverages the higher spatial resolution of functional neuroimaging in humans has revealed a ventrolateral ATL region that responds equally to all types of concepts, including social, object and abstract concepts, be they referenced by verbal and/or non-verbal stimuli (Binney et al., 2016; Rice et al., 2018). Activation of the dorsal-polar ATL, on the other hand, appears to be more sensitive to socially-relevant semantic stimuli (Binney et al., 2016; Rice et al., 2018; Zahn et al., 2007b). These observations support a proposal in which the broadly-defined ATL region can be characterised as a domain-general supramodal semantic hub with graded differences in relative specialisation towards certain types of conceptual information (Binney et al., 2012; Binney et al., 2016; Lambon Ralph et al., 2017; Plaut, 2002; Rice et al., 2015). Our results reveal that the temporal poles are reliably activated across four social domains – moral reasoning, empathy for emotions, ToM and trait inference. They do not, however, provide support for the involvement of the ventrolateral ATL. We argue this is likely due to technical and methodological limitations of the fMRI studies included in the meta-analyses (see Visser et al., 2010). Most notably this includes vulnerability to susceptibility artefacts that cause BOLD signal drop-out and geometric distortions around certain brain areas, including the ventral ATLs (Jezzard and Clare, 1999; Ojemann et al., 1997). Studies that have used PET, which is not vulnerable to such artefacts, or techniques devised to overcome limitations of conventional fMRI (Devlin et al., 2000; Embleton et al., 2010), reveal activation in both the temporal poles and the ventral ATL in response to social stimuli (Balgova et al., in prep; Binney et al., 2016; Castelli et al., 2002).

### 4.3. Limitations

Because semantic control demands were not explicitly manipulated in the social contrasts we included, our results cannot directly confirm our hypothesis regarding the specific contribution made by the SCN in social cognition. Our conclusions rely on an assumption that overlap reflects a generalised neurocomputation upon which both semantic control and social processing rely. The alternative explanation is that overlapping activation reflects tightly yet separately packed cognitive functions which may only dissociate when investigated at an increased spatial resolution (Henson, 2006; Humphreys et al., 2020). Moreover, we chose to pool across heterogeneous samples of studies to investigate the cognitive domains of interest. The advantage of this approach is that it identifies activation that is generalisable across highly variable experimental conditions and washes out spurious findings associated with idiosyncratic properties of stimuli and/or paradigms. However, the preponderance of specific experimental procedures in each literature addressed still unintentionally led to systematic differences in the characteristics of the studies used to define the different cognitive domains. For example, the semantic control dataset included studies that employed verbal stimuli almost exclusively, while the majority of empathy studies employed non-verbal stimuli. Some of the differences between the associated networks (e.g, in lateralization) might therefore be attributable to verbal processing demands. As is the case with all meta-analyses, therefore, some aspects of our results should be treated with caution.

Another limitation of this study is that most of the experiments included used control conditions that were highly matched to their experimental conditions in terms of the demand for domain-general processes such as cognitive control and semantic processing, and therefore they may have subtracted away much of the activation we were aiming to explore. Despite this, we did find consistent activation of the SCN, particularly the left IFG, across all four social domains. This may be because, although a considerable subset of included experiments had high-matching control conditions, not all may have properly controlled for semantic control demands specifically. An alternative explanation is that processing socially-relevant conceptual knowledge may impose greater demands on the SCN. Consistent with this, it has been shown that processing social concepts relative to non-social concepts led to increased activation of the SCN even when controlling for potentially confounding psycholinguistic factors (Binney et al., 2016).

### 4.4. Concluding remarks and future directions

Regions of the SCN are engaged by several types of complex social tasks, including ToM, empathy, trait inference and moral reasoning. This finding sheds light on the nature and neural correlates of the cognitive control mechanisms which contribute to the regulation of social cognition and specifically implicates processes involved in the goal-directed retrieval of conceptual knowledge. Importantly, our current findings and our broader set of hypotheses can be generalised to multiple social phenomena, thereby contributing a unified account of social cognition. Future research will need to establish a causal relationship between the SCN and successful regulation of social processing. This could be done by directly probing whether SCN regions are sensitive to manipulations of semantic control demands within a social task. Similarly, the capacity for neurostimulation of SCN regions to disrupt social task performance needs to be investigated.

Elucidating the neural bases of social control and representation may help us understand the precise nature of social impairments resulting from damage to different neural systems. For example, our framework (Binney & Ramsey, 2020) predicts that damage to representational areas such as the ATL will impair social information processing irrespective of task difficulty or the need to integrate context. In contrast, we expect that damage to control areas would lead to impaired social processing specifically when it requires selecting from amongst alternative interpretations of social cues, and/or retrieving social information that is only weakly associated with a person or a situation. Damage to perisylvian frontal and/or temporo-parietal areas (comprising the SCN) leads to semantic aphasia, a disorder characterized by impaired access and use of conceptual knowledge (Corbett et al., 2009; Jefferies et al., 2008, 2007; Jefferies and Lambon Ralph, 2006; Noonan et al., 2010). This contrasts with ATL damage which leads to semantic dementia, a condition associated with a loss or degradation of semantic knowledge (including social knowledge; Hodges and Patterson, 2007; Lambon Ralph et al., 2010; Lambon Ralph and Patterson, 2008; Patterson et al., 2007; Rogers et al., 2004). As far as we are aware, the extent to which brain damage that leads to semantic aphasia also affects social abilities has not yet been formally investigated. Some insight can be found in neurodegenerative patients with prominent frontal lobe damage, where social impairments can be linked to deficits in executive function (Healey and Grossman, 2018; Kamminga et al., 2015). More generally, it will be interesting to discover whether a distinction between knowledge representation and cognitive control can inform our understanding of the precise nature of atypical or disordered social cognition in, for example, the context of dementia, acquired brain injury, autism spectrum conditions and schizophrenia.

## Supporting information

SI2. List of included experiments

SI3. Results tables

## Acknowledgements

This research was performed as part of an all-Wales ESRC Doctoral Training Programme +3 PhD studentship awarded to VD [ES/P00069X/1]. The authors would like to thank Irina Giurgea for her assistance in preparing the data for publication, and Ionela Bara and Eva Balgova for their comments on a previous version of this manuscript.

