## Supplementary material for "Establishing a Role of the Semantic Control Network in Social Cognitive Processing: A Meta-analysis of Functional Neuroimaging Studies": SI1. Additional information

for

**Contents:**

**Section S1. Literature search and data extraction** p.2

S.1.1. Deviations from the pre-registration

S.1.1. Comparison between the present meta-analyses and the prior meta-analyses

S.1.2. Data extraction

S.1.3. Missing information

S.1.4. Overview of the study selection process

**Section S2. Data analysis** p.9

**Section S3. Results** p.10

S3.1. False belief reasoning

S3.2. Global Empathy (for Pain & Emotions)

S3.3. A common network for multiple sub-domains of social cognition

S3.4. Explicit versus Implicit Social Paradigms

S3.5. The difficulty characteristics of the experiments included in the dataset for each social domain

**References** p.17

### Section S1. Literature search and data extraction.

*S.1.1. Deviations from the pre-registration*

We pre-registered our plans for the current study after accessing some of the raw data and conducting independent ALE analyses on subsets of the final semantic control, theory of mind and trait inference datasets (see the OSF pre-registration for further details: osf.io/fktb8/). There are two deviations from our pre-registered protocols which are detailed below:

1. *Inclusion/exclusion criteria*. We pre-registered the exclusion of contrasts between experimental conditions and low-level baselines (e.g., rest, fixation cross). However, we subsequently decided to collect and include data from contrasts against low-level baselines because they can reveal activity associated with domain-general cognitive processes that is subtracted out by contrasts between active conditions. This could include semantic and executive processes that are common to both social and non-social tasks and which were of interest for testing our pre-registered hypotheses. For completeness, we report the analyses whilst excluding this subset of contrasts on the project's OSF page (osf.io/fktb8/).
2. *Data collection*. In our pre-registration, empathy was considered a single sub-domain of social cognition. However, the relevant literature distinguishes between empathy for affective states and empathy for pain because there are important differences in the brain regions underpinning these two types of empathic processing (Ding et al., 2020; Kogler et al., 2020; Timmers et al., 2018). Thus, pooling across empathy for pain and emotions would preclude the identification of all brain regions engaged in empathy. Therefore, we made a distinction between experiments looking at empathy for pain and those investigating empathy for emotions, and conducted all analyses separately for these two subsets. For completeness, the results of a global empathy (pooled across pain and emotions) are reported in Section S3.

*S.1.2. Comparison between the present meta-analyses and the prior meta-analyses*

Table S1. A summary of the similarities and differences in the methodological approach taken in the present meta-analyses and the prior meta-analyses upon which updates were based.

| | Semantic Control Noonan et al. (2013) | Theory of Mind Molenberghs et al. (2016) | Empathy Timmers et al., (2018) | Moral Reasoning Eres et al., (2018) |
|---|---|---|---|---|
| Task definition | | c | | |
| Contrast definition | | d | | e |
| Included results from both fMRI and PET studies | | | | |
| Included whole-brain analyses only | | | | |
| Included multiple contrasts from a single sample | | | | |
| Adjusted for multiple contrasts from a single sample | a | b | a | a |
| Employed an updated GingerALE algorithm thereby avoiding overly liberal statistical thresholds (g) | | a | | |
| Employed recommended FWE cluster-extent correction (g) | f | f | f | f |
| Employed additional conservative height-based thresholding | | | | |

Orange coloured cells indicate differences whereas green cells indicate an identical or equivalent approach. FWE = Family Wise Error
[a]Not reported;
[b]Not applicable;
[c] Unlike the present study, Molenberghs et al. (2016) included studies of irony comprehension, trait inference & those employing 'interactive games' (see Section 2.1.2);
[d] Unlike the present study, Molenberghs et al. (2016) did not include contrasts between an experimental social condition and low-level (e.g., fixation) baseline conditions (see Section 2.1.2);

[e] Unlike the present study, Eres et al., (2018) included contrasts between different components of moral reasoning (e.g., utilitarian vs. deontological moral judgements) and did not include contrasts between experimental conditions and low-level baseline conditions(see Section 2.1.2);
[f] Used False Discovery Rate correction.
[g](Eickhoff et al., 2016)

*S.1.3. Data extraction*

We extracted the following information from each study and included it in our database: authors, year of publication, sample size, imaging method (fMRI/PET), task description, contrast, type of contrast (low-level baseline vs high-level baseline), coordinate space (MNI/Talairach), type of instructional cue (explicit/implicit) and type of difficulty category based on participant reaction times (experimental condition harder/easier than or equally difficult to the control condition). In addition, we collected information about other potential moderating factors not investigated in this study as follows:

- Semantic control: type of cognitive manipulation (semantic production/ semantic decision/ homonyms/ metaphors), input modality (verbal/non-verbal, visual/auditory).
- ToM: type of ToM inference (cognitive/affective), stimulus type (animations/ cartoon/ photos/ RMET/ video/ stories), task type (e.g., false belief reasoning, predicting behaviours based on mental states), input modality, difficulty based on accuracy scores (experimental condition harder/easier than or equally difficult to the control condition).
- Empathy: target of empathy (e.g., pain, happiness, sadness etc.), valence of target of empathy (positive/negative), input modality, difficulty based on accuracy scores.
- Trait inference: type of inference (prior person knowledge/appearance/ behaviour), trait (e.g., trustworthiness, approachability etc.), input modality, difficulty based on accuracy scores.
- Moral reasoning: input modality, difficulty based on accuracy scores.

*S.1.4. Missing information*

If activation coordinates from whole-brain analyses for the contrast of interest were not reported in the published article or in supplementary materials, the authors of the study were contacted to obtain the missing information. If this was unsuccessful, the study was excluded from the meta-analysis. If the data was obtained directly from personal communication with the authors, this is clearly indicated in section 2 of the supplementary materials.

#### *S.1.5. Overview of the study selection process*

In accordance with the PRISMA guidelines (Page et al., 2021), a detailed overview of the study selection process is provided below separately for each dataset. Figures S1- S5 depict the number of articles identified and screened at each stage, the number of articles excluded at each stage, as well as the final number of articles and experiments included in the analyses.

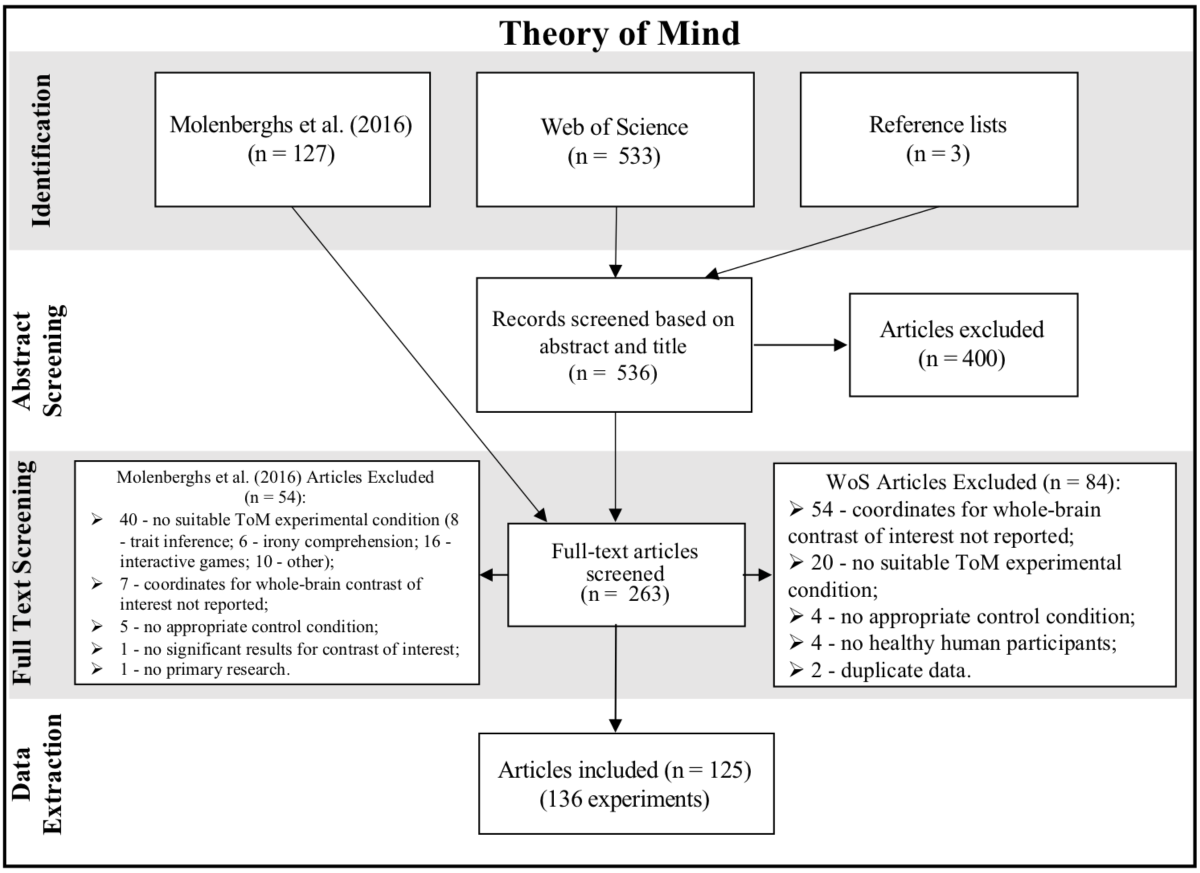


Figure S1. Overview of the selection process for the **theory of mind** meta-analysis. We included 136 experiments with a total number of 2158 coordinates and 3452 participants.

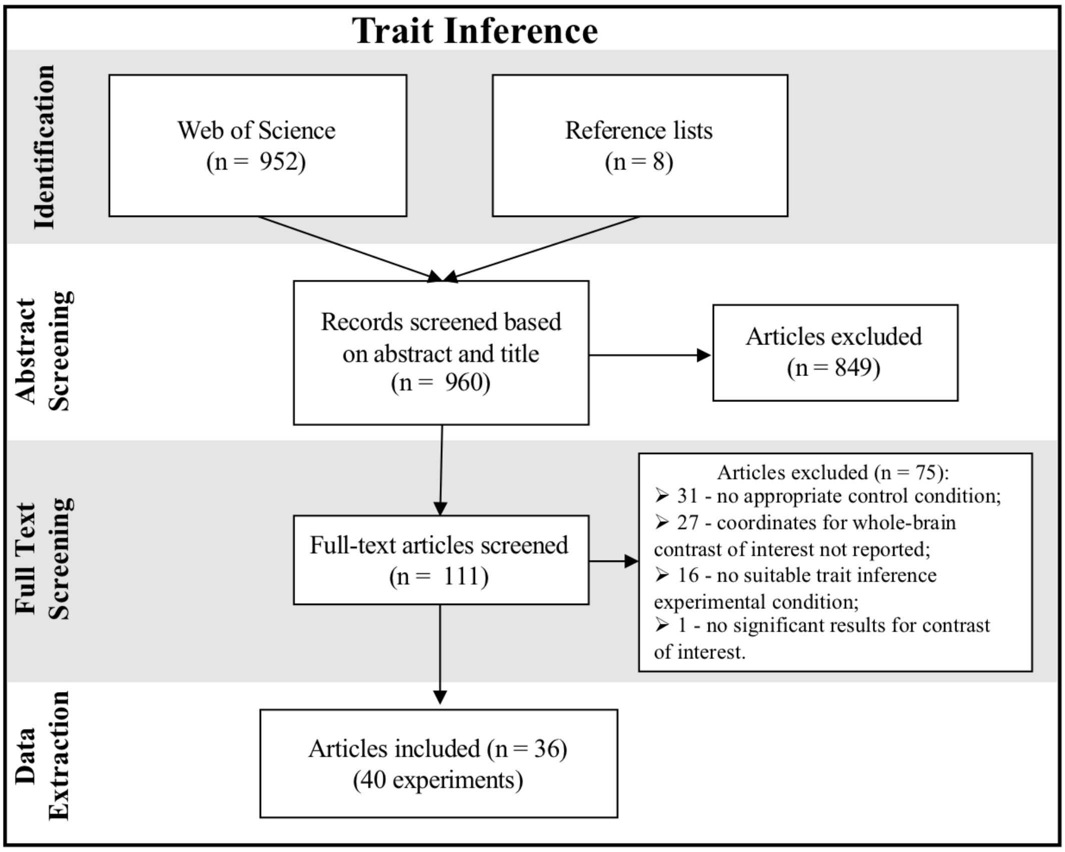


Figure S2. Overview of the selection process for the **trait inference** meta-analysis. We included 40 experiments with a total number of 523 coordinates and 732 participants.

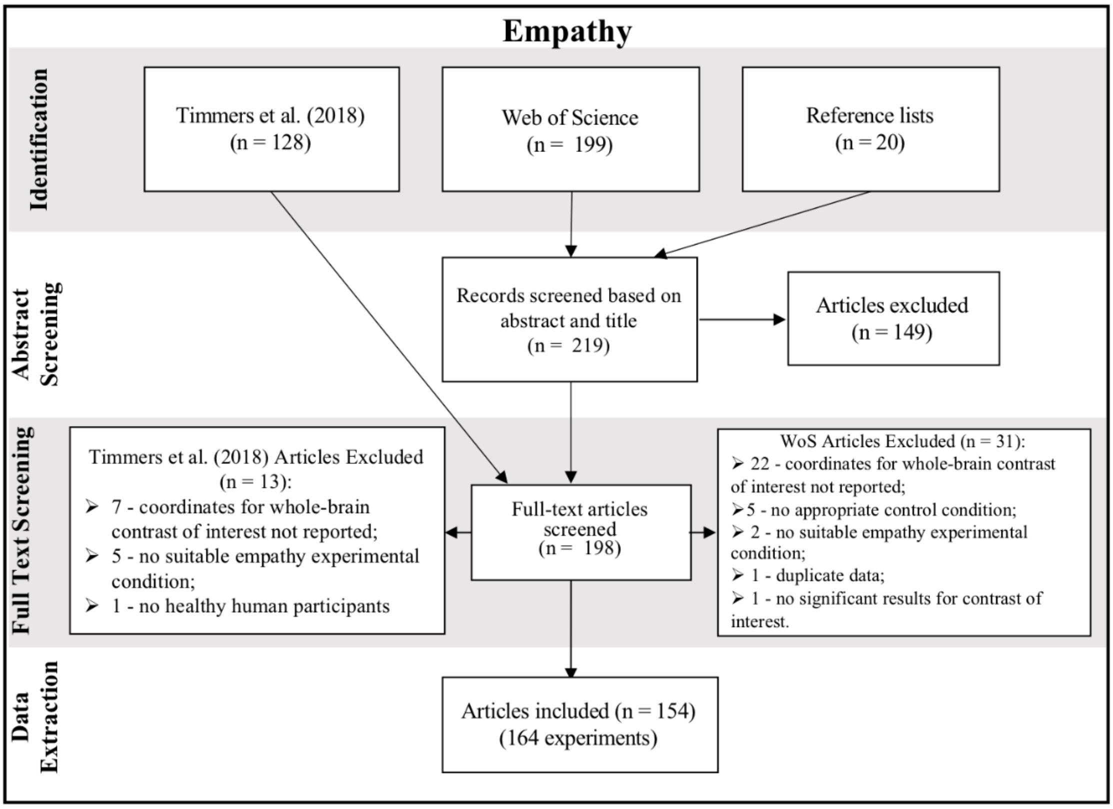


Figure S3. Overview of the selection process for the **empathy** meta-analysis. We included 164 experiments with a total number of 2704 coordinates and 4423 participants.

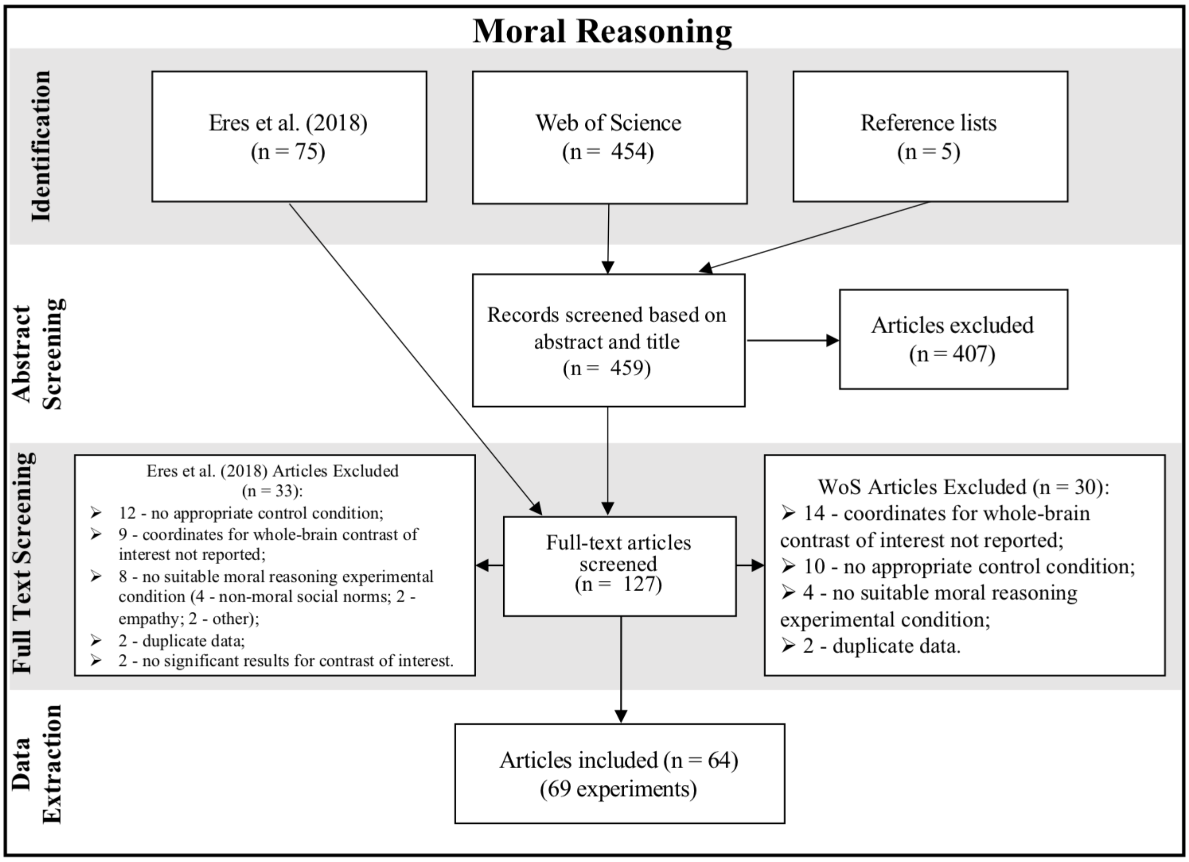


Figure S4. Overview of the selection process for the **moral reasoning** meta-analysis. We included 69 experiments with a total number of 909 coordinates and 1609 participants.

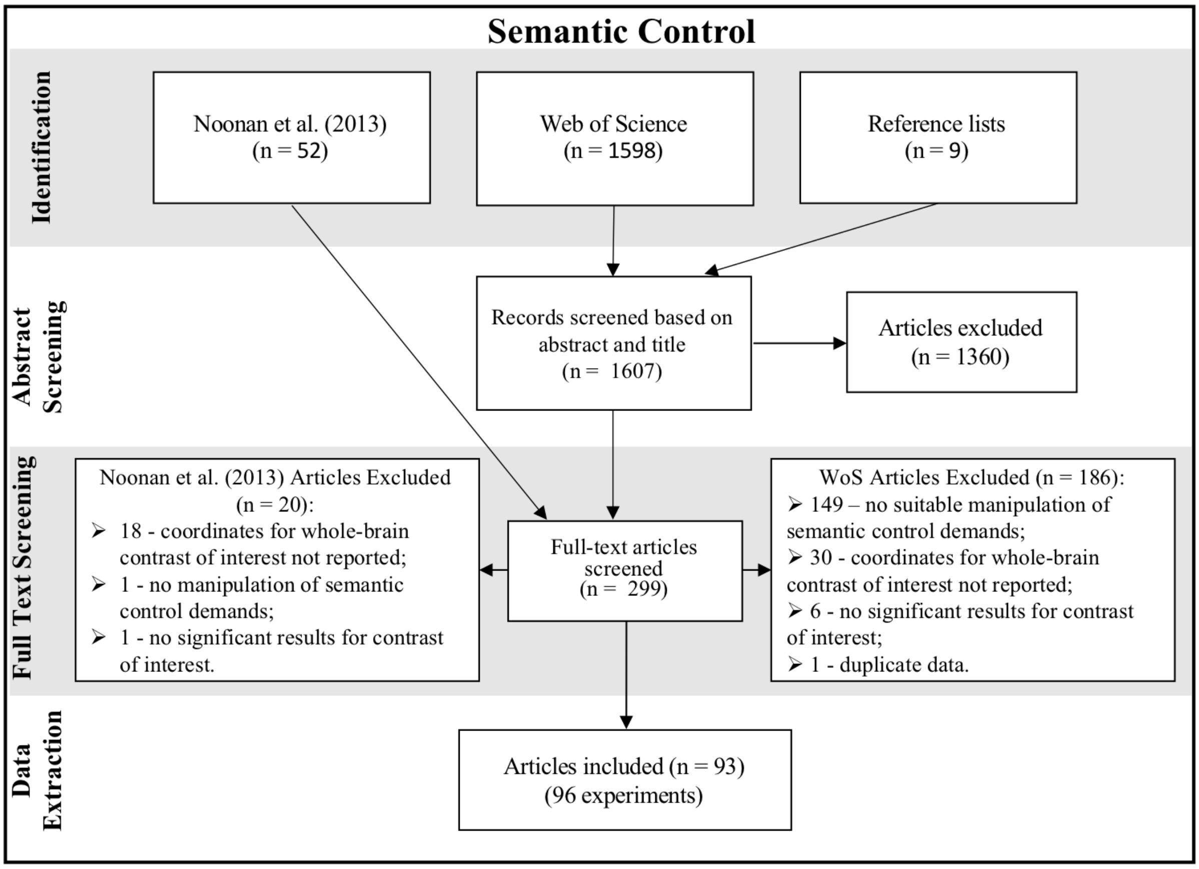


Figure S5. Overview of the selection process for the **semantic control** meta-analysis. We included 96 experiments with a total number of 981 coordinates and 2052 participants.

### Section S2. Data analysis

A list of all analyses conducted is provided in the table below. For each analysis, the table specifies the type of analysis (C - confirmatory/ E - exploratory), the results section where the findings are described, the results table number and the results figure number.

Table S2. List of all analyses conducted.

| Analysis | | | | Type | Results Section | Results Table | Figure |
|---|---|---|---|---|---|---|---|
| Semantic control | | | | C | 3.2 | S2 | 3 |
| Social cognition | Independent sub-domains | Low baselines excluded | | C | OSF page | | - |
| | | Theory of mind | Global Theory of Mind (all tasks) * | C | 3.1.1 | S1.1.1 | 1a |
| | | | False belief reasoning* | E | S3.1 | S1.1.2 | S6a |
| | | Trait Inference* | | C | 3.1.2 | S1.2 | 1b |
| | | Empathy | Global Empathy* | C | 3.1.3 | S1.3.1 | S7a |
| | | | Empathy for Pain* | E | 3.1.3 | S1.3.2 | 1d |
| | | | Empathy for Emotions* | E | 3.1.3 | S1.3.3 | 1c |
| | | | Empathy for Pain vs. Emotions^ | E | 3.1.3 | S1.3.4 | S7b |
| | | Moral reasoning* | | C | 3.1.4 | S1.4 | 1e |
| | Overlay conjunction | | | E | S3.3 | 2.1.5 | 2; S7 |
| | Explicit vs. Implicit | Empathy for Emotions | Explicit* | C | S3.4 | S4.1.1 | 5a |
| | | | Implicit* | C | S3.4 | S4.1.2 | 5a |
| | | | Explicit vs. Implicit^ | C | S3.4 | S4.2.1 | 5d |
| | | Empathy for Pain | Explicit* | C | S3.4 | S4.1.3 | 5b |
| | | | Implicit* | C | S3.4 | S4.1.4 | 5b |
| | | | Explicit vs. Implicit^ | C | S3.4 | S4.2.2 | 5e |
| | | Moral reasoning | Explicit* | C | S3.4 | S4.1.5 | 5c |
| | | | Implicit* | C | S3.4 | S4.1.6 | 5c |
| | | | Explicit vs. Implicit^ | C | S3.4 | S4.2.3 | 5f |
| | | Cluster analyses | | E | 3.4.2 | - | S8 |
| | Task difficulty | E>C ToM* | | E | 3.5 | S5.1 | 6a |
| | | E=C ToM* | | E | 3.5 | S5.2 | 6a |
| | | C>E ToM* | | E | 3.5 | S5.4 | 6a |
| | | E>C vs. E=C^ | | E | 3.5 | S5.3 | - |
| | | ToM known difficulty category* | | E | - | S5.5 | 6c |
| | | Cluster analysis | | E | 3.5 | - | 6c |

| | | | | | | |
|---|---|---|---|---|---|---|
| Social Cognition vs. Semantic Control | Theory of Mind vs. SC | Global ToM (all tasks) vs. SC^ | C | 3.3.1 | S3.1 | 4a |
| | | False belief reasoning vs. SC^ | E | S3.1 | S3.1.2 | S6b |
| | Trait inference vs. SC^ | | C | 3.3.2 | S3.2 | 4b |
| | Empathy vs. SC | Empathy for Emotions vs. SC^ | E | 3.3.3 | S3.3 | 4c |
| | | Empathy for Pain vs. SC^ | E | 3.3.4 | S3.4 | 4d |
| | Moral reasoning vs. SC^ | | C | 3.3.5 | S3.5 | 4e |

* Indicates independent ALE analyses. Each independent ALE analysis were repeated using two different statistical thresholding approaches (see the Results section).

^ Indicates contrast and conjunction analyses.

SC = Semantic control; E = exploratory analysis; C = confirmatory analysis

### Section S3. Results

*S3.1. False belief reasoning*

The validity of some of the tasks used to investigate ToM processing has been debated (Heyes, 2014; Oakley et al., 2016; Obhi, 2012; Quesque and Rossetti, 2020). For example, Quesque & Rossetti (2020) argue that some classic ToM tasks (e.g., Heider and Simmel) do not necessarily require the representation of other’s mental states and successful completion can be explain by lower-level cognitive processes (e.g., basic associative learning mechanisms). A further criticism has been that they do not necessitate distinguishing one’s own mental states from others’ mental states (Heyes, 2014). False belief tasks, on the other hand, are frequently used in both healthy and clinical populations (e.g., Yirmiya et al., 1998), as well as in developmental contexts (e.g., Wellman et al., 2001), and are more commonly accepted as a suitable task for probing ToM and for identifying the underlying brain network (Dodell-Feder et al., 2011; Saxe and Kanwisher, 2003). False belief tasks require participants to make inferences about the (false) beliefs of protagonists which may contrast their own knowledge about the state of reality, meeting both aforementioned criteria for valid ToM tasks (Quesque and Rossetti, 2020).

Our global ToM meta-analysis pooled across a variety of ToM task paradigms, not all of which may be considered valid measures of ToM processing. Therefore, we performed a separate analysis focused solely on experiments which employed false belief tasks to explore any differences in the underlying brain network when using a more conservative task definition. Across 43 false belief reasoning experiments, the ALE analysis revealed convergent activation in 9 clusters including in the precuneus, bilateral pMTG & STG, bilateral mid MTG (extending anteriorly towards the temporal pole), right MFG, mPFC, medial OFC and cerebellum (Figure S6a; Table S1.1.2). All these clusters survived both extent-based and height-based thresholding. Unlike in the inclusive ToM analysis, we did not find convergent activation in the bilateral IFG. Overlap between the neural network underpinning semantic control (i.e., SCN & regions of the MDN) and false belief reasoning was found in a single left pMTG cluster (Figure S6b; Table S3.1.2).

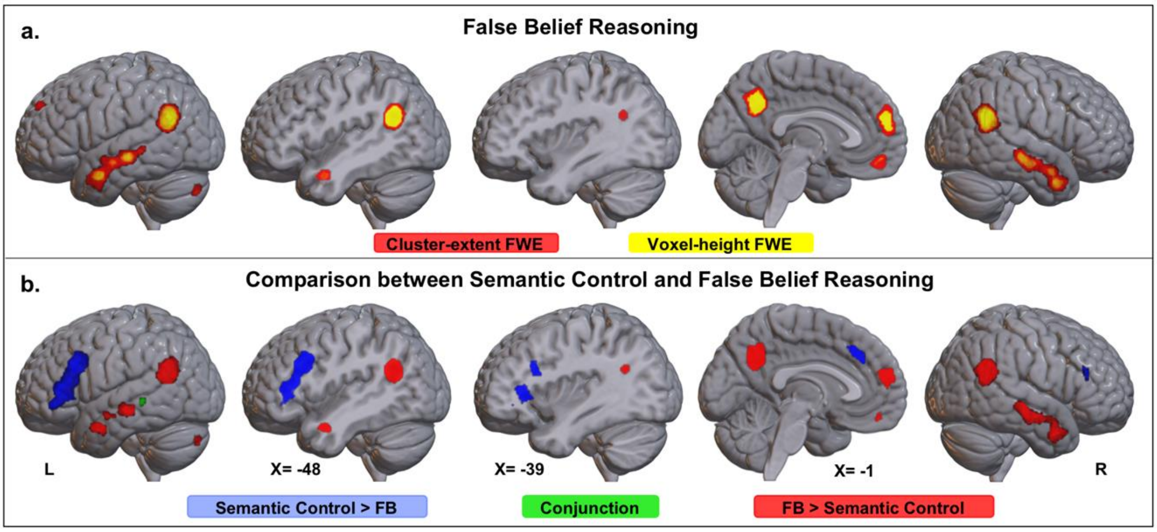


Figure S6. a) Binary whole-brain ALE maps showing statistically significant convergent activation resulting from independent meta-analyses of false belief reasoning experiments (N=43). The ALE maps were thresholded using an FWE corrected cluster-extent at $p < .05$ with a cluster-forming threshold of $p < .001$ (red) and, an FWE corrected voxel-height threshold of $p < .05$ (yellow). b) Results of the contrast (blue: semantic control > false belief reasoning; red: false belief reasoning > semantic control) and conjunction (green) analyses between the ALE maps associated with false belief reasoning and semantic control. The contrast maps were thresholded with a cluster-forming threshold of $p < .001$ and a minimum cluster size of 200 $mm^3$. The lateral views, which show projections on the cortical surface, are accompanied by brain slices at the sagittal midline and also coplanar with the peak of the left IFG [X = -39] and pSTG [X = -48] clusters that overlapped across all social domains (Table S1.5). FB = false belief reasoning.

*S3.2. Global Empathy (for Pain & Emotions)*

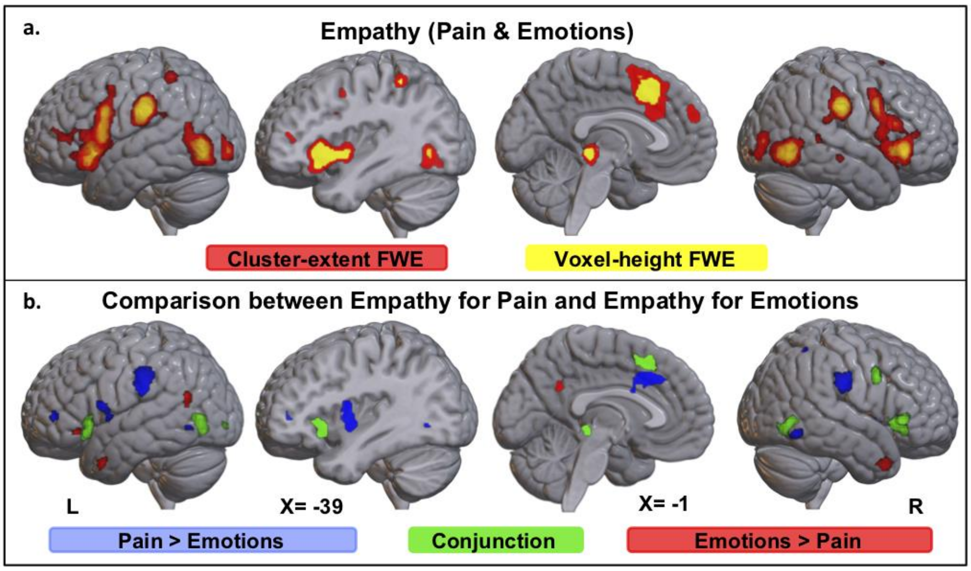


Figure S7. a) Binary whole-brain ALE maps showing statistically significant convergent activation resulting from independent meta-analyses of all empathy studies (N=164), including both empathy for pain and empathy for emotions. The ALE maps were thresholded using an FWE corrected cluster-extent at $p$ < .05 with a cluster-forming threshold of $p$ < .001 (red) and, an FWE corrected voxel-height threshold of $p$ < .05 (yellow). b) Results of the contrast (blue: empathy for pain > empathy for emotions; red: empathy for emotions > empathy for pain) and conjunction (green) analyses between the ALE maps associated with empathy for pain and empathy for emotions. The contrast maps were thresholded with a cluster-forming threshold of $p$ < .001 and a minimum cluster size of 200 $mm^3$. The lateral views, which show projections on the cortical surface, are accompanied by brain slices at the sagittal midline and also coplanar with the peak of the left IFG cluster that overlapped across all social domains [Table S1.5; X = -39]).

*S3.3. A common network for multiple sub-domains of social cognition*

To identify brain areas consistently activated across multiple sub-domains of social cognition, we performed an overlay conjunction analysis of the ALE maps associated with ToM, trait inference, empathy (for pain and/or emotions) and moral reasoning. The ALE maps were thresholded using p<.001 uncorrected. Convergent activation across all four socio-cognitive sub-domains was found in the bilateral IFG (pars orbitalis), mPFC, precuneus, left pSTG, and bilateral ATL (Figure S7).

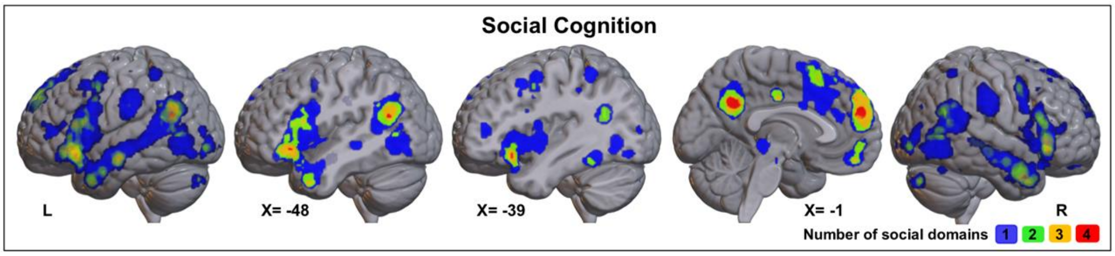


Figure S7. An overlay conjunction of the ALE maps resulting from independent meta-analyses on ToM, trait inference, empathy for pain/emotions, and moral reasoning. The map displays the number of social domains showing convergent activation in each voxel. The ALE maps were thresholded using an uncorrected cluster-forming threshold of $p$ < .oo1. The lateral views, which show projections on the cortical surface, are accompanied by brain slices at the sagittal midline and also coplanar with the peak of the left pSTG (X = -48) and left IFG (X = -39) clusters that overlapped across all four social domains.

#### *S3.4. Explicit versus Implicit Social Paradigms*

##### *S3.4.1. Conjunctions and Contrasts between Explicit and Implicit Experiments*

Empathy for affective states

An ALE analysis across all explicit paradigms showed convergent activation in the bilateral IFG, mPFC, SMA, precuneus, middle cingulate gyrus, right temporal pole, left anterior ITG, left caudate and brainstem. An ALE analysis of all implicit experiments revealed convergent activation in the bilateral IFG, bilateral pMTG and right occipital cortex. The conjunction analysis between implicit and explicit empathy for affective states revealed overlapping activation in two clusters located in pars triangularis of the right IFG and pars orbitalis of the left IFG. Formal contrasts revealed that, compared to implicit paradigms, explicit tasks also engage the dmPFC. No significant clusters were found to be consistently activated for implicit but not explicit paradigms.

Empathy for pain

The ALE analysis of empathy for pain studies employing explicit instructional cues revealed convergent activation in the bilateral insula (extending to the IFG), pars triangularis of left IFG (extending to MFG), the middle cingulate gyri (extending to SMA), bilateral SMG, left inferior occipital gyrus (extending to pMTG) left thalamus and right pallidum. An

ALE analysis of studies using an implicit paradigm revealed convergent activation in the bilateral IFG, bilateral precentral gyrus, bilateral insula, SMA, middle cingulate gyrus, bilateral SMG, left IPL, bilateral posterior ITG, calcarine fissure, left thalamus, right amygdala and brainstem. The conjunction between implicit and explicit empathy for pain revealed overlapping activation in the left IFG (pars opercularis), bilateral insula, dmPFC, middle cingulate cortex, SMA, bilateral SMG and left inferior occipital gyrus. Formal contrasts found that explicit paradigms differentially engage dmPFC while implicit paradigms show additional convergent activation in left inferior occipital cortex and right hippocampus.

Moral reasoning

Convergent activation across explicit paradigms was found in the left insula cortex, medial OFC, dmPFC, precuneus and bilateral pMTG. Convergent activation across implicit paradigms was found in the left IFG (pars opercularis), dmPFC and right anterior MTG. The conjunction between implicit and explicit moral reasoning paradigms revealed overlapping activation in mPFC, while the formal contrasts did not reveal any above-threshold differential activation.

*3.4.2. Cluster analyses investigating the contribution of explicit/implicit experiments to ALE clusters*

In addition, we conducted exploratory cluster analyses to investigate whether the explicit and implicit experiments contributed similarly to each of the significant ALE clusters found for each social domain. Given that the sample sizes of implicit experiments were too small to conduct independent ALE analyses for the ToM and Trait Inference datasets, this cluster analysis allowed us to understand whether a specific type of instructional cue drove the convergent activation in any of the identified clusters. For each social domain, we calculated the proportion of implicit and explicit experiments that contributed at least one peak to each identified cluster. The results (Figure S7) revealed that explicit and implicit paradigms contributed equally to most activation clusters (all $p > .05$), with the exception of the precuneus cluster in response to ToM ($p = .023$, explicit > implicit) and a cluster located in the right primary visual cortex in response to empathy for pain ($p = .016$, implicit > visual cortex in response to empathy for pain ($p = .016$, implicit > explicit). This suggests that implicit and explicit experiments contributed equally to most activation clusters regardless of social domain.

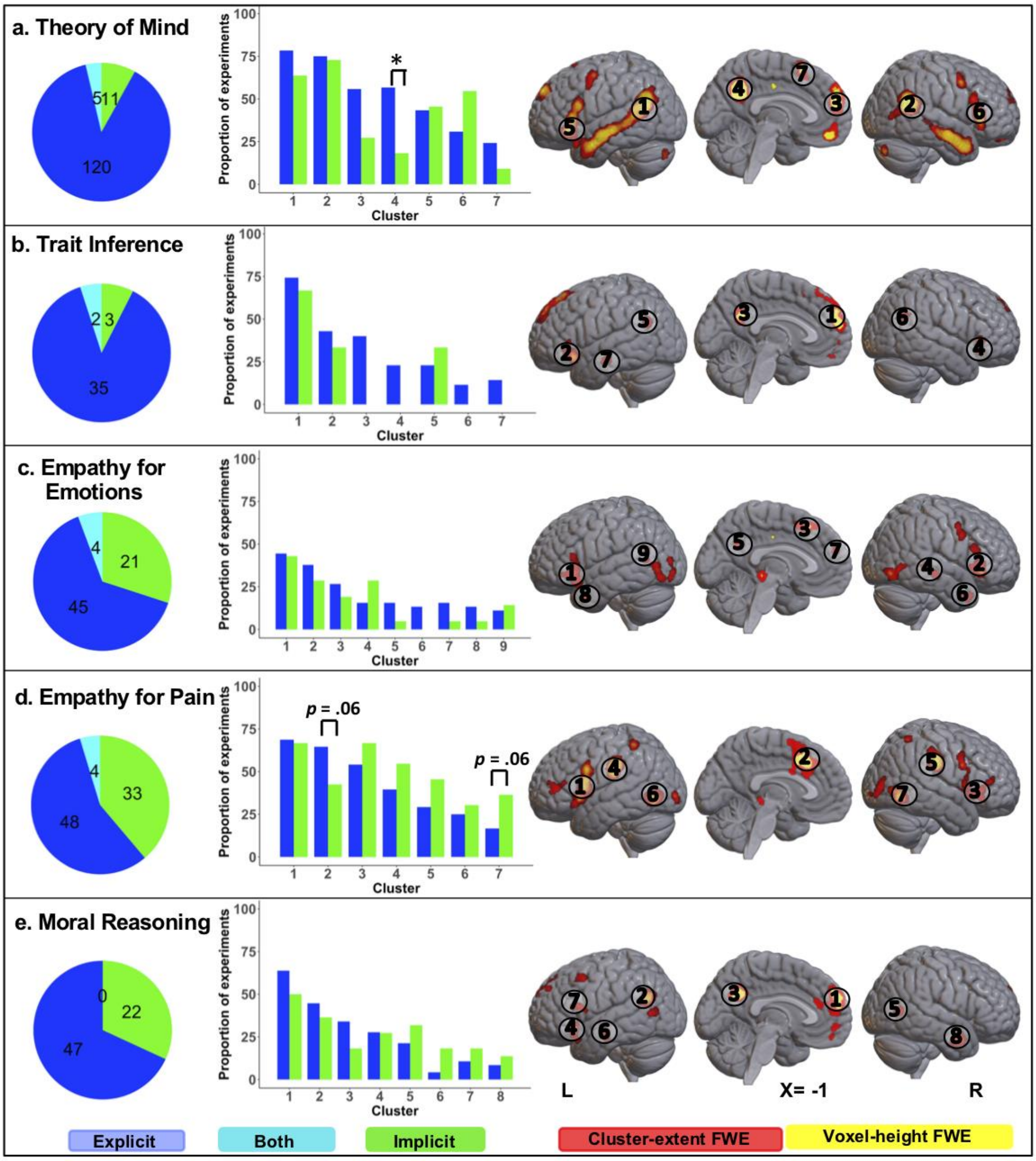


Figure S8. Pie charts illustrate the proportion of explicit, implicit and both (i.e., contrasts reporting coordinates from both implicit and implicit instructions) experiments that were included in the meta-analysis for each social domain. The labels indicate the experiment count for each category. The bar charts display the proportion of experiments in each category (explicit/implicit) that contributed at least one peak to the clusters of interest. The ALE map of convergent activation for each social domain is displayed on a standard MNI brain and each cluster of interest is indicated and numbered. * $p < .05$

*S3.5. The difficulty characteristics of the experiments included in the dataset for each social domain*

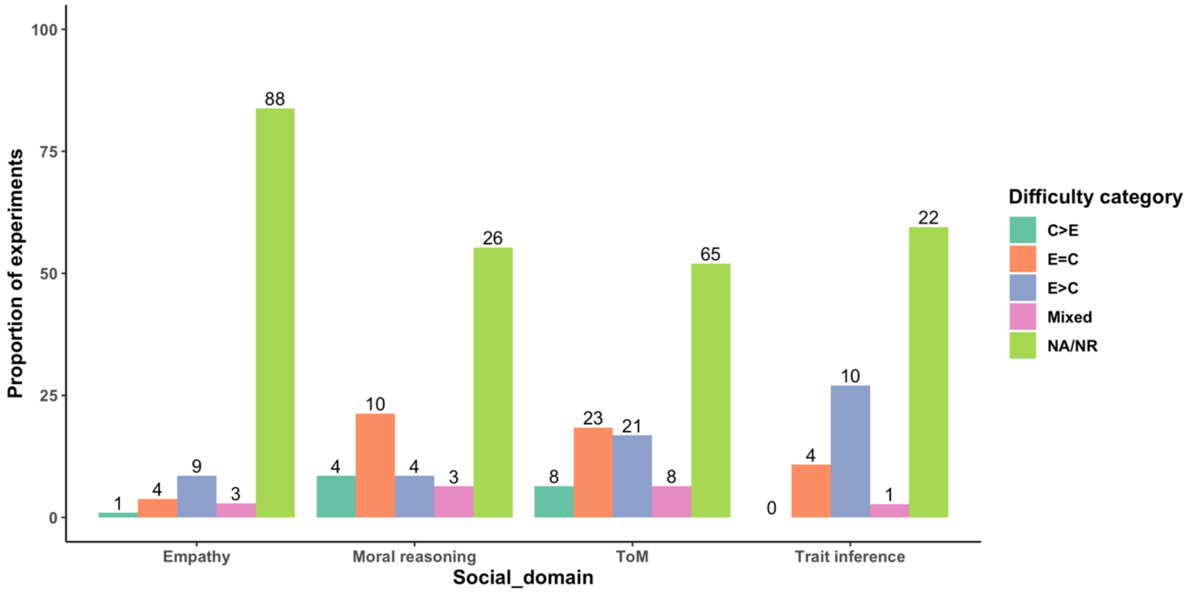


Figure S9. For each social domain, the proportion of studies featuring specific difficulty characteristics is illustrated. The bars represent the proportion of studies featuring specific difficulty characteristics. The exact number of experiments featuring specific difficulty characteristics is also displayed for each bar. C>E – reaction times were significantly faster in the experimental condition compared to the control condition suggesting that the control condition was more difficult. E=C – the was no statistically significant different between the reaction times in the experimental and control conditions suggesting equally difficult conditions. E>C – reaction times were significantly faster in control condition compared to the experimental condition suggesting that the experimental condition was more difficult. Mixed – the experiment pooled across contrasts belonging to different difficulty categories. NA – not applicable, NR – not reported.
