## Supplementary material for "Establishing a Role of the Semantic Control Network in Social Cognitive Processing: A Meta-analysis of Functional Neuroimaging Studies": SI2. List of included experiments

for

### A Meta-analysis of Functional Neuroimaging Studies

Veronica Diveica, Kami Koldewyn & Richard J. Binney

This document contains lists with all the experiments included in the meta-analysis and their characteristics and associated references. The complete raw datasets can be accessed via OSF (osf.io/fktb8/).

<u>Contents:</u>

Table S1. **Theory of Mind** p.2

Table S2. **Trait Inference** p.20

Table S3. **Empathy** p.27

Table S4. **Moral Reasoning** p.52

Table S5. **Semantic Control** p. 62

**Table S1.** List of studies included in the **theory of mind** (N = 136) meta-analysis.

| Authors | Imaging method | No. of peaks | N | Task | Contrast | Difficulty RT | Difficulty Accuracy | Instructional cue |
|---|---|---|---|---|---|---|---|---|
| Abraham et al., 2008 | fMRI | 22 | 17 | Story comprehension questions | stories about mental states > control stories | E>C | E=C | Explicit |
| Abraham et al., 2010 | fMRI | 14 | 22 | Infer the beliefs/desires of the protagonist | Mental state inference > syllogistic reasoning (belief and desire conjunction) | C>E | NR | Explicit |
| Adams et al., 2010 | fMRI | 21 | 28 | Choose the word that best describes the mental state or gender of the person depicted | mental state > gender judgement | NR | NR | Explicit |
| Aichhorn et al., 2009 | fMRI | 13 | 21 | False belief localizer task | false beliefs > false photographs | E>C | E>C | Explicit |
| | | | | False belief localizer task | false beliefs > false photographs (timepoint 2 question) | | | Explicit |
| Alderson-Day et al., 2016 | fMRI | 10 | 19 | Story ending judgement | ToM > physical causality | NR | NR | Explicit |
| Alkire et al., 2018 | fMRI | 7 | 28 | Predict other's behaviour based on their mental state/ Semantic judgements | predict behaviour > semantic judgements | E=C | E=C | Explicit |
| Bahnemann et al., 2010 | fMRI | 9 | 25 | Affective mental state / Physical appearance judgement | mental state inference > gender judgements | E>C | E=C | Explicit |
| Baron-Cohen et al., 1999 | fMRI | 53 | 12 | Choose the word that best describes the mental state or gender of the person depicted | mental state > gender judgement | NR | NR | Explicit |
| Bartholomeusz et al., 2018 | fMRI | 5 | 22 | Story ending judgement | ToM > physical causality | NR | NR | Explicit |
| Bliksted et al., 2019 | fMRI | 6 | 17 | Animated shapes task - physical judgements | intentional > random movement | | | Implicit |
| Bodden et al., 2013 | fMRI | 15 | 30 | Yoni task | affective ToM > physical | NR | E=C | Explicit |
| | | | | Yoni task | cognitive ToM > physical | NR | E=C | Explicit |
| Briend et al., 2019 | fMRI | 13 | 28 | Attend to the protagonists' mental states | known (ToM) > unknown (non-ToM) language | NA | NA | Explicit |
| Brune et al., 2008 | fMRI | 14 | 13 | Inferring the intentions/expectations of the | correct (ToM) > jumbled (non-ToM) cartoon sequence | NR | NR | Explicit |

| | | | | | | | | |
|---|---|---|---|---|---|---|---|---|
| | | | | protagonists (ToM) and physical judgements (non-ToM) | | | | |
| Brunet et al., 2000 | PET | 17 | 8 | Story ending judgement | ToM > physical causality | E=C | E=C | Explicit |
| Canessa et al., 2012 | fMRI | 34 | 27 | Indicate when a landscape stimulus appeared | cooperative social interactions > landscapes | | | Implicit |
| | | | | Indicate when a landscape stimulus appeared | affective social interactions > landscapes | | | Implicit |
| Cassidy et al., 2020 | fMRI | 19 | 40 | False belief localizer task | false belief > false photograph | NR | NR | Explicit |
| Cassidy et al., 2020 | fMRI | 15 | 35 | False belief localizer task | false belief > false photograph | NR | NR | Explicit |
| Castelli et al., 2000 | PET | 10 | 6 | Passive observation (cued and un-cued conditions) | intentional > random movement | NR | NR | Explicit/Implicit |
| Castelli et al., 2010 | fMRI | 18 | 12 | Choose the word that best describes the mental state or gender of the person depicted | mental state > gender judgement | NR | NR | Explicit |
| Castelli et al., 2010 | fMRI | 19 | 12 | Choose the word that best describes the mental state or gender of the person depicted | mental state > gender judgement | NR | NR | Explicit |
| Chakroff e al., 2016 | fMRI | 4 | 23 | False belief localizer task | false belief > false photograph | NR | NR | Explicit |
| Cheung et al., 2012 | fMRI | 25 | 20 | Sally-Ann Task Adaptation - Mental state inference/ Perceptual judgements | false belief > physical judgements | C>E | C>E | Explicit |
| | | | | Sally-Ann Task Adaptation - Mental state inference/ Perceptual judgements | true belief > physical judgements | C>E | C>E | Explicit |
| | | | | Sally-Ann Task Adaptation - Mental state inference/ Perceptual judgements | false belief > physical judgements | E>C | E>C | Explicit |
| | | | | Sally-Ann Task Adaptation - Mental state inference/ Perceptual judgements | true belief > physical judgements | E=C | E>C | Explicit |
| Cole et al., 2019 | fMRI | 9 | 20 | Infer the intention behind an action (only for unsuccessful trials) / Judge whether the action was successful | ToM > non-ToM judgements | NR | NR | Explicit |

| | | | | | | | | |
|---|---|---|---|---|---|---|---|---|
| Contreras et al., 2013 | fMRI | 14 | 25 | Mental state inference/ Physical inference | ToM (group) > physical judgements | E>C | NA | Explicit |
| | | | | Mental state inference/ Physical inference | ToM(group member) > physical judgements | E=C | NA | Explicit |
| Contreras et al., 2013 | fMRI | 18 | 13 | Mental state inference/ Physical inference | ToM > physical inference | E=C | NA | Explicit |
| | | | | False belief localizer task | false belief > false photograph | C>E | C>E | Explicit |
| Corradi-Dell'Acqua et al., 2014 | | 18 | 46 | Mental state / physical judgements | mental state > physical judgements | C>E | C>E | Explicit |
| | | | | Mental state / physical judgements | mental state > physical judgements | E=C | E=C | Explicit |
| Das et al., 2012 | fMRI | 27 | 19 | Animated shapes task - Passive observation | ToM > random movement | | | Implicit |
| de Achaval et al., 2012 | fMRI | 19 | 14 | Choose the emotion that best describes the facial expression | mental state > gender judgement | E>C | E=C | Explicit |
| | | | | Choose the word that best describes the mental state or gender of the person depicted | mental state > gender judgement | E>C | E=C | Explicit |
| Deuse et al., 2016 | fMRI | 30 | 38 | Valence judgement regarding social scenes (from the percpective of the interaction partners depicted) / regarding location in non-social scenes | social > non-social scenes | C>E | NR | Explicit |
| Dodell-Feder et al., 2011 | fMRI | 16 | 62 | False belief localizer task | false beliefs > false photographs | NR | NR | Explicit |
| Dodell-Feder et al., 2014 | fMRI | 52 | 18 | Jusge whether the statement is conisstent with the story | stories about thoughts > appearance | NR | NR | Explicit |
| | | | | Jusge whether the statement is conisstent with the story | stories about thoughts > appearance | NR | NR | Explicit |
| Dohnel et al., 2012 | fMRI | 8 | 18 | Sally-Ann Task Adaptation - Mental state inference/ Physical judgements | mental state inference > physical judgements | E=C | E=C | Explicit |

| | | | | | | | | |
|---|---|---|---|---|---|---|---|---|
| Dominguez et al., 2019 | | 5 | 20 | Choose the word that best describes the mental state or gender/age of the person depicted | ToM > age judgements | NR | NR | Explicit |
| Dufour et al., 2013 | fMRI | 20 | 27 | False belief localizer task | false beliefs > false photographs | NR | NR | Explicit |
| Ferstl & von Cramon, 2002 | fMRI | 13 | 9 | Take the perspective of the protagonist and focus on their mental states/ Same/different pseudo-language judgements | ToM stories > pseudo-sentences | NR | NR | Explicit |
| Fletcher et al, 1995 | PET | 9 | 6 | Mental state inference /physical judgements | ToM > physical stories | C>E | E=C | Explicit |
| | | | | Mental state inference /physical judgements | ToM stories > unlinked sentences | C>E | E=C | Explicit |
| Focquaert et al., 2010 | fMRI | 8 | 12 | Choose the word that best describes the mental state or gender of the person depicted | mental state > gender judgement | NR | NR | Explicit |
| Focquaert et al., 2010 | fMRI | 5 | 12 | Choose the word that best describes the mental state or gender of the person depicted | mental state > gender judgement | NR | NR | Explicit |
| Gallagher et al., 2000 | fMRI | 10 | 6 | Consider the meaning of the cartoons | ToM (attribution of false belief/ignorance) > non- ToM (no mental state attribution) cartoons | | | Implicit |
| | | | | Mental state attribution/physical inferences | ToM > non-ToM stories | | | Implicit |
| Gao et al., 2019 | fMRI | 17 | 35 | Story ending judgement | affective ToM > physical causality | E=C | E=C | Explicit |
| | | | | Story ending judgement | cognitive ToM > physical causality | E=C | E=C | Explicit |
| Geiger et al., 2019 | fMRI | 6 | 32 | Mental state/ physical inference | mental state inference > movement identification | NR | E>C | Explicit |
| Gobbini et al., 2007 | fMRI | 42 | 12 | Animated shapes task - Choose an appropriate description for the movie clip | Intentional social > random movement | NR | NR | Explicit |
| | | | | False Belief Task Adaptation | false beliefs > physical stories | NR | NR | Explicit |
| Gweon et al., 2012 | fMRI | 6 | 20 | Indicate whether the probe sentence matches the story. | ToM > physical stories | E=C | E=C | Explicit |
| Gweon et al., 2012 | fMRI | 6 | 8 | Indicate whether the probe sentence matches the ToM/ phisycal story | ToM > physical judgements | E=C | E=C | Explicit |

| | | | | | | | | |
|---|---|---|---|---|---|---|---|---|
| Hartwright et al., 2015 | | 13 | 21 | False belief localizer task | false belief > false photograph | NR | E=C | Explicit |
| Hennion et al., 2016 | fMRI | 9 | 25 | Animated shapes task - categorize the type of interaction (ToM, goal-directed, random) | ToM > goal-directed movement & ToM > random movement conjunction | NR | E=C | Explicit |
| Herve et al., 2013 | fMRI | 20 | 42 | Categorize the type of mental state (deception, belief, emotion)/ Semantic plausibility judgement | ToM > semantic judgements | E>C | E>C | Explicit |
| Hooker et al., 2008 | fMRI | 23 | 20 | Infer the emotion of the protagonist if they knew what was about to happen/ recognize their emotion | infer emotion > recognize emotion (false belief trials) | E>C | E>C | Explicit |
| Hooker et al., 2010 | fMRI | 20 | 15 | Identify the type of change (physical, social, no change) | change in the protagonist's mental state > no change | E>C | E>C | Explicit |
| Jack & Pelphrey, 2015 | fMRI | 19 | 34 | Animated shapes task - Passive observation | ToM > random movement | | | Implicit |
| Jacoby et al., 2016 | fMRI | 21 | 17 | False belief localizer task<br><br>Passive movie watching | false belief > false photograph<br><br>ToM > pain events | NR | NR | Explicit<br><br>Implicit |
| Jenkins & Mitchell, 2010 | fMRI | 9 | 15 | Mental state/physical inference | ToM > physical inference | NR | NR | Explicit |
| Jenkins et al., 2014 | | 15 | 19 | Story comprehension (no overt response)<br>False belief localizer task | ToM > non-ToM control stories<br><br>false belief > false photograph | <br><br>NR | <br><br>NR | Implicit<br><br>Explicit |
| Jimura et al., 2010 | fMRI | 4 | 34 | False belief reasoning/factual judgements | ToM > factual judgements | E>C | E=C | Explicit |
| Kana et al., 2009 | fMRI | 12 | 12 | Animated shapes task - Choose an appropriate description | Intentional social > random movement | E>C | E>C | Explicit |
| Kana et al., 2015 | fMRI | 9 | 13 | Animated shapes task - Choose an appropriate description for the movie clip | ToM > random movement | NR | NR | Explicit |
| Kandylaki et al., 2015 | fMRI | 17 | 20 | False belief attribution/physical judgements<br>False belief attribution/physical judgements | false belief > physical causality story passage<br>ToM > physical judgements | E=C<br><br>E=C | E=C<br><br>E=C | Explicit<br><br>Explicit |

| | | | | | | | | |
|---|---|---|---|---|---|---|---|---|
| Kanske et al., 2015 | fMRI | 23 | 178 | EmpaToM | ToM > factual reasoning | E=C | C>E | Explicit |
| Kanske et al., 2015 | fMRI | 49 | 25 | EmpaToM | ToM > factual reasoning | C>E | C>E | Explicit |
| | | | | False belief localizer task | false belief > false photograph | NR | NR | Explicit |
| Kim et al., 2016 | fMRI | 13 | 14 | Yoni task | cognitive ToM > physical | NR | NR | Explicit |
| | | | | Yoni task | affective ToM > physical | NR | NR | Explicit |
| Kirkovski et al., 2016 | fMRI | 13 | 23 | Animated shapes task - categorize the type of interaction (ToM, goal-directed, random) | ToM > random movement | NR | NR | Explicit |
| | | | | Animated shapes task - categorize the type of interaction (ToM, goal-directed, random) | ToM > goal-directed movement | NR | NR | Explicit |
| | | | | Animated shapes task - categorize the type of interaction (ToM, goal-directed, random) | ToM > rest | | | Explicit |
| Kliemann et al., 2008 | fMRI | 6 | 26 | False belief localizer task | false beliefs > false photographs | NR | NR | Explicit |
| Kobayashi et al., 2006 | fMRI | 5 | 16 | Infer the protagonist's belief/ Physical inference | false belief > physical causality | E=C | E=C | Explicit |
| Kobayashi et al., 2006 | fMRI | 4 | 16 | Infer the protagonist's belief/ Physical inference | false belief > physical causality | E=C | E=C | Explicit |
| Kobayashi et al., 2007 | fMRI | 7 | 12 | Infer the protagonist's belief/ Physical inference | false belief > physical causality | E=C | E=C | Explicit |
| | | | | Infer the protagonist's belief/ Physical inference | false belief > physical causality | E=C | E=C | Explicit |
| Kobayashi et al., 2007 | fMRI | 9 | 12 | Infer the protagonist's belief/ Physical inference | false belief > physical causality | E=C | E=C | Explicit |
| | | | | Infer the protagonist's belief/ Physical inference | false belief > physical causality | E=C | E=C | Explicit |
| Kobayashi et al., 2007 | fMRI | 5 | 28 | Infer the protagonist's belief/ Physical inference | false belief > physical causality | E=C | E=C | Explicit |
| Koelkebeck et al., 2011 | fMRI | 7 | 15 | Animated shapes task - Judge the interaction type | Intentional social > random movement | NR | NR | Explicit |

| | | | | | | | | |
|---|---|---|---|---|---|---|---|---|
| Lavoie et al., 2016 | fMRI | 19 | 19 | Intention/physical inference | ToM > physical inference (action description phase) | E=C | E=C | Explicit |
| | | | | Intention/physical inference | ToM > physical inference (contextual information phase) | | | Explicit |
| | | | | Intention/physical inference | ToM > physical inference (response phase) | | | Explicit |
| Lee & McCarthy, 2016 | fMRI | 14 | 19 | False belief localizer task | false belief > false photograph | E=C | NR | Explicit |
| Lee et al., 2011 | fMRI | 30 | 13 | False belief localizer task | false beliefs > false photographs | NR | C>E | Explicit |
| | | | | False belief reasoning / factual judgements | false belief > factual judgements | NR | E>C | Explicit |
| Lewis et al., 2017 | fMRI | 16 | 17 | False belief attribution/ factual judgements | ToM > factual judgements | E=C | E=C | Explicit |
| Libero et al., 2014 | fMRI | 3 | 22 | Ordinary/Unusual judgement about mental states or means | intention > means judgement | E=C | E=C | Explicit |
| Lin et al., 2018 | fMRI | 11 | 39 | False belief localizer task | false belief > false photograph | NR | NR | Explicit |
| Malhi et al., 2008 | fMRI | 16 | 20 | Animated shapes task - Passive observation | Intentional social > random movement | | | Implicit |
| Marjoram et al., 2006 | fMRI | 3 | 13 | Understand the joke | ToM> non-ToM cartoons | NR | NA | Explicit |
| Martin & Weisberg, 2003 | fMRI | 11 | 12 | Animated shapes task variant - Choose an appropriate description for the movie clip | social interaction > mechanical movement | NR | NR | Explicit |
| Mason et al., 2008 | fMRI | 27 | 18 | Animated shapes task variant - Choose an appropriate description for the movie clip | ToM inference > rest | NA | NA | Explicit |
| | | | | Animated shapes task variant - Choose an appropriate description for the movie clip | ToM inference > rest | | | Explicit |
| McAdams & Krawczyk, 2013 | fMRI | 12 | 17 | Animated shapes task - Social/perceptual judgements | Intentional social > random movement | NR | NR | Explicit |
| Mier et al., 2010 | fMRI | 36 | 16 | Predict the behaviour of the person depicted/ Emotion recognition | Mental state inference > emotion categorization | NR | NR | Explicit |
| | | | | Predict the behaviour of the person depicted | Mental state inference > rest | | | Explicit |
| Mitchell, 2008 | fMRI | 4 | 20 | False belief localizer task | false beliefs > false photographs | C>E | NR | Explicit |

| | | | | | | | | |
|---|---|---|---|---|---|---|---|---|
| Modinos et al., 2010 | fMRI | 8 | 36 | Yoni task | ToM > physical | E>C | E=C | Explicit |
| Moessnang et al., 2016 | fMRI | 43* | 46 | Animated shapes task - categorize the type of interaction (ToM, goal-directed, random) | ToM > goal-directed movement | NR | NR | Explicit |
| | | | | Animated shapes task - categorize the type of interaction (ToM, goal-directed, random) | ToM > random movement | NR | NR | Explicit |
| Mohnke et al., 2016 | fMRI | 18 | 297 | Detect changes in the protagonist's affective state / no. of human beings depicted | ToM > physical judgements | E>C | E=C | Explicit |
| Moor et al., 2012 | fMRI | 8 | 55 | RMET Adaptation - Choose the word that best describes the mental state or gender/age of the person depicted | mental state > gender/age judgement | E>C | E>C | Explicit |
| Moran et al., 2012 | fMRI | 34 | 31 | Animated shapes task - Passive observation | Intentional social > random movement | | | Implicit |
| | | | | False belief localizer task | false belief > false photograph | NR | C>E | Explicit |
| Moran et al., 2012 | fMRI | 12 | 17 | Animated shapes task - Passive observation | Intentional social > random movement | | | Implicit |
| | | | | False belief localizer task | false belief > false photograph | NR | C>E | Explicit |
| Moriguchi et al., 2007 | fMRI | 17 | 16 | Animated shapes task - Passive observation | Intentional social > random movement | | | Implicit |
| Mukerji et al., 2019 | fMRI | 9 | 32 | False belief localizer task | false belief > false photograph | E>C | E>C | Explicit |
| Naughtin et al., 2017 | fMRI | 11 | 22 | False belief localizer task | false belief > false photograph | NR | NR | Explicit |
| Nieminen-von Wendt et al., 2003 | PET | 9 | 8 | Mental state inference /physical judgements | ToM > physical stories | NR | NR | Explicit |
| Ohnishi et al., 2004 | fMRI | 13 | 11 | Animated shapes task - Passive observation | Intentional social > random movement | | | Implicit |
| Oliver et al. 2018 | fMRI | 9 | 35 | False belief localizer task | false belief > false photograph | NR | NR | Explicit |
| Otsuka et al., 2009 | fMRI | 18 | 22 | Judge story coherence based on the protagonist's mental state/tense | mental state > tense judgement | | | Explicit |

| | | | | | | | | |
|---|---|---|---|---|---|---|---|---|
| | | | | Judge story coherence based on the protagonist's mental state | mental state judgement > rest | | | Explicit |
| Otti et al. 2015 | fMRI | 10 | 20 | Animated shapes task - Passive observation | ToM > random movement | | | Implicit |
| | | | | Animated shapes task - Passive observation | ToM > rest | | | Implicit |
| Overgaauw et al. 2015 | fMRI | 9 | 32 | Choose the word that best describes the mental state or gender/age of the person depicted | mental state >gender/age judgements | E>C | E>C | Explicit |
| Perner et al., 2007 | fMRI | 13 | 19 | False belief localizer task variant | false belief > false photograph | C>E | NR | Explicit |
| | | | | False belief localizer task variant | false belief > temporal change | E>C | NR | Explicit |
| Platek et al., 2004 | fMRI | 6 | 5 | RMET Adaptation - Think about the mental state of the people depicted | face stimuli > rest | NA | NA | Explicit |
| Powell et al., 2017 | fMRI | 12 | 12 | Story ending judgement | ToM > physical causality | NR | NR | Explicit |
| Roser et al., 2012 | fMRI | 40 | 14 | Inferring the intentions/expectations of the protagonists (ToM) and physical judgements (non-ToM) | correct (ToM) > jumbled (non-ToM) cartoon sequence | NR | NR | Explicit |
| Ross & Olson, 2010 | fMRI | 10 | 15 | Animated shapes task - Social/perceptual judgements | Intentional social > random movement | NR | C>E | Explicit |
| Russel et al., 2000 | fMRI | 4 | 7 | Choose the word that best describes the mental state or gender of the person depicted | mental state > gender judgement | NR | NR | Explicit |
| Saft et al., 2013 | fMRI | 12 | 26 | Inferring the intentions/expectations of the protagonists (ToM) and physical judgements (non-ToM) | correct (ToM) > jumbled (non-ToM) cartoon sequence | NR | NR | Explicit |
| Samson et al., 2008 | fMRI | 18 | 17 | Understand the joke | ToM > non-ToM visual puns cartoons | | | Implicit |
| | | | | Understand the joke | ToM > non-ToM semantic cartoons | | | Implicit |
| Saxe et al., 2006 | fMRI | 8 | 12 | False belief localizer task | false belief > false photograph | C>E | NA | Explicit |
| Saxe & Kaniwisher, 2003 | fMRI | 5 | 25 | Story comprehension (no judgements) | false belief > mechanical inference stories | | | Implicit |

| | | | | | | | | |
|---|---|---|---|---|---|---|---|---|
| Saxe & Kaniwisher, 2003 | fMRI | 5 | 28 | False belief localizer task | false beliefs > false photographs | C>E | NA | Explicit |
| Saxe & Powell, 2006 | fMRI | 9 | 12 | False belief localizer task | false beliefs > false photographs | C>E | NA | Explicit |
| Schiffer et al., 2013 | fMRI | 6 | 22 | Choose the word that best describes the mental state or gender of the person depicted | mental state > gender judgement | E>C | E>C | Explicit |
| Schlaffke et al., 2014 | | 23 | 39 | Inferring the mental states of the protagonists (ToM) and physical judgements (non-ToM) | correct (ToM) > jumbled (non-ToM) cartoon sequence | NR | NR | Explicit |
| | | | | Inferring the mental states of the protagonists (ToM) and physical judgements (non-ToM) | correct (ToM) > jumbled (non-ToM) cartoon sequence | NR | NR | Explicit |
| Schmitgen et al., 2016 | fMRI | 32 | 21 | Detect changes in the protagonist's affective state / no. of human beings depicted | ToM > physical judgements | NR | NR | Explicit |
| Schneider et al., 2014 | fMRI | 7 | 16 | False belief localizer task | false belief > false photograph | E=C | E=C | Explicit |
| Sebastian et al., 2012 | fMRI | 13 | 30 | Story ending judgement | attribution of emotion > physical causality judgement | E=C | E=C | Explicit |
| | | | | Story ending judgement | attribution of intention > physical causality judgement | E=C | E=C | Explicit |
| Shimada et al., 2018 | fMRI | 26* | 30 | Choose the word that best describes the mental state or gender of the person depicted | ToM > gender judgement | NR | NR | Explicit |
| Sommer et al., 2010 | fMRI | 8 | 14 | Infer the protaginist's feelings/ Physical judgement | ToM inference > physical judgement | NR | NR | Explicit |
| Specht & Wigglesworth, 2018 | fMRI | 15 | 18 | Story ending judgement | ToM > physical causality | NR | NR | Explicit |
| Spunt & Adolphs, 2014 | fMRI | 15 | 29 | Why/How judgement task | why > how inferences | E>C | E>C | Explicit |
| Spunt & Adolphs, 2014 | fMRI | 14 | 21 | Why/How judgement task | why > how inferences | E>C | E>C | Explicit |
| Spunt & Lieberman, 2012a | fMRI | 10 | 21 | Judge why/how the action is performed (no overt response) | why > how judgements (conjunction video & text) | E=C | NR | Explicit |

| | | | | | | | | |
|---|---|---|---|---|---|---|---|---|
| Spunt & Lieberman, 2012b | fMRI | 34 | 22 | Judge why/how the facial expression is displayed (no overt response) | why > how judgements | E>C | NA | Explicit |
| Spunt et al., 2011 | fMRI | 23 | 15 | Judge why/how the action is performed (no overt response) | why > how judgements | C>E | NA | Explicit |
| | | | | Judge why the action is performed/ what the action is (no overt response) | why > what judgements | E>C | NA | Explicit |
| Tholen et al., 2020 | fMRI | 19 | 130 | EmpaToM | ToM > factual reasoning | E=C | C>E | Explicit |
| Thye et al., 2018 | fMRI | 25 | 18 | Choose the word that best describes the mental state or gender of the person depicted | mental state > gender judgement | NR | E>C | Explicit |
| | | | | Choose the word that best describes the mental state or gender of the person heard | mental state > gender judgement | NR | E>C | Explicit |
| | | | | Story ending judgement | ToM > physical causality | NR | E>C | Explicit |
| Van der Meer et al., 2011 | fMRI | 40 | 19 | Sally-Ann Task Adaptation | mental state > physical inference | E>C | E=C | Explicit |
| | | | | Sally-Ann Task Adaptation | mental state inference > rest | | | Explicit |
| Van Hoeck et al., 2014 | fMRI | 4 | 19 | False belief attribution/physical inference | false belief > physical inference | E>C | E=C | Explicit |
| Vanderwal et al., 2008 | fMRI | 15 | 17 | Animated shapes task - Social/perceptual judgements | Intentional social > random movement | NR | C>E | Explicit |
| Vogeley et al., 2001 | fMRI | 20 | 8 | Mental state attribution/physical inferences | ToM > non-ToM stories | NR | E=C | Explicit |
| | | | | Mental state attribution/semantic judgements | ToM (self not in the story) > unlinked sentences | | | Explicit |
| | | | | Mental state attribution/semantic judgements | ToM (self as one of the agents in the story) > unlinked sentences | | | Explicit |
| Vollm et al., 2006 | fMRI | 13 | 13 | Story ending judgement | attribution of intention > physical causality | NR | E=C | Explicit |
| Walter et al., 2004 | fMRI | 39 | 13 | Story ending judgement | communicative intention > physical causality | E=C | E=C | Explicit |
| | | | | Story ending judgement | private intention (one agent) > physical causality | E=C | E=C | Explicit |

| | | | | | | | | |
|---|---|---|---|---|---|---|---|---|
| | | | | Story ending judgement | private intention (two agents) > physical causality | E=C | E=C | Explicit |
| Walter et al., 2004 | fMRI | 40 | 12 | Story ending judgement | private intention > physical causality | E=C | E=C | Explicit |
| | | | | Story ending judgement | prospective intention > physical causality | E=C | E=C | Explicit |
| | | | | Story ending judgement | communicative intention > physical causality | E=C | E=C | Explicit |
| Walter et al., 2009 | fMRI | 39 | 12 | Story ending judgement | private intention > physical causality | NR | NR | Explicit |
| | | | | Story ending judgement | prospective social intention > physical causality | NR | NR | Explicit |
| | | | | Story ending judgement | communicative intention > physical causality | NR | NR | Explicit |
| Wang et al. 2015 | fMRI | 7 | 56 | Story ending judgement | ToM > physical causality | NR | NR | Explicit |
| Willert et al., 2015 | fMRI | 22 | 81 | Detect changes in the protagonist's affective state / no. of human being depicted | ToM > physical judgements | E>C | E>C | Explicit |
| Wolf et al., 2010 | fMRI | 20 | 18 | Passive observation | ToM > non-ToM scenes (movie watching phase) | E>C | E=C | Explicit |
| | | | | Infer the mental states of the social agents/physical judgements (covert response) | ToM > physical inferences (response phase 1) | | | Explicit |
| | | | | Infer the mental states of the social agents/physical judgements (covert response) | ToM > physical inferences (response phase 2) | | | Explicit |
| Young et al., 2011 | fMRI | 6 | 17 | False belief localizer task | false beliefs > false photographs | NR | NR | Explicit |
| Zaitchik et al., 2010 | fMRI | 13 | 15 | Story comprehension | stories about beliefs > non-ToM stories | C>E | C>E | Explicit |
| | | | | Story comprehension | stories about emotions > non-ToM stories | C>E | C>E | Explicit |

* the coordinates were obtained via personal communication with the authors.
N = sample size; E>C - the experimental condition was more difficult than the control condition; C>E - the control condition was more difficult than the experimental condition; E=C - the experimental and control conditions were equally difficult; NR - not reported; NA - not applicable because no behavioural data was collected.

**Table S2.** List of studies included in the **trait inference** (N = 40) meta-analysis.

| Authors | Imaging method | No. of peaks | N | Task | Contrast | Difficulty RT | Difficulty Accuracy | Instructional cue |
|---|---|---|---|---|---|---|---|---|
| Ames & Fiske, 2013 | fMRI | 3 | 19 | Form an opinion about the person depicted based on the statement presented | impression formation > remember statement order | NA | NA | Explicit |
| Baas et al., 2008 | fMRI | 4 | 21 | Judge whether the person depicted is trustworthy | trait inference > age judgement | NR | NR | Explicit |
| Baetens et al., 2014 | fMRI | 26 | 18 | Infer the personality trait that could drive the behaviour/Describe the behaviour | trait inference > description of behaviour | NR | E>C | Explicit |
| Benoit et al., 2010 | fMRI | 18 | 13 | Trait descriptiveness - determine the degree to which the trait presented is an appropriate description of the target person | trait inference (friend) > syllable counting | E=C | NA | Explicit |
| Bos et al., 2012 | fMRI | 2 | 16 | Judge whether the person depicted is trustworthy | trait inference > age judgement | NR | NA | Explicit |
| Brent et al., 2014 | fMRI | 11 | 22 | Trait descriptiveness - determine whether the trait presented is an appropriate description of the target person | trait inference (mother) > valence judgement | E>C | NA | Explicit |
| Chen et al., 2010 | fMRI | 14 | 21 | Decide whether they liked the face presented<br>Decide whether they liked the face presented | likeability > gender judgement<br>likeability > rest | E>C | NA | Explicit<br>Explicit |
| D'Argembeau et al., 2008 | fMRI | 30* | 20 | Trait descriptiveness - determine whether the trait presented is an appropriate description of the target person (currently & 5 years prior) | trait inference (friend/sibling) > valence judgement | E>C | NA | Explicit |
| Debbane et al., 2017 | fMRI | 23 | 44 | Trait descriptiveness - determine the degree to which the trait presented is an appropriate description of the target person<br>Trait descriptiveness - determine the degree to which the trait presented is an appropriate description of the target person | positive trait inference (friend) > syllable counting<br>negative trait inference (friend) > syllable counting | NR<br>NR | NA<br>NA | Explicit<br>Explicit |

| | | | | | | | | |
|---|---|---|---|---|---|---|---|---|
| Gilron & Gutchess, 2012 | fMRI | 60 | 20 | Form an impression about the person depicted based on the statement presented | impression formation > semantic judgements | NR | NA | Explicit |
| | | | | Form an impression about the person depicted based on the statement presented/ Semantic judgement | diagnostic > neutral statements | NR | NA | Explicit/Implicit |
| Hall et al., 2010 | fMRI | 12 | 24 | Judge whether the person depicted is approachable | Approachability > gender judgements | E>C | E>C | Explicit |
| | | | | Judge whether the person depicted is intelligent | intelligence > gender judgements | E>C | E>C | Explicit |
| Hall et al., 2012 | fMRI | 10 | 25 | Judge whether the person depicted is approachable | Approachability > gender judgements | NR | NR | Explicit |
| | | | | Judge whether the person depicted is approachable | Approachability judgement > rest | | | Explicit |
| | | | | Judge whether the person depicted is intelligent | Intelligence > gender jugements | | | Explicit |
| | | | | Judge whether the person depicted is intelligent | Intelligence jugements > rest | | | Explicit |
| Hall et a., 2012 | fMRI | 4 | 22 | Judge whether the person depicted is intelligent | Intelligence > gender judgements | NR | NR | Explicit |
| | | | | Judge whether the person depicted is intelligent | Intelligence judgement > rest | | | Explicit |
| | | | | Judge whether the person depicted is approachable | Approachability judgement > rest | | | Explicit |
| Harris et al., 2005 | fMRI | 12 | 12 | Causal attribution - Attribute the event described to the person, the stimulus or the situation | person > stimulus/situation attributions | NR | NR | Explicit |
| Heatherton et al., 2006 | fMRI | 3 | 30 | Trait descriptiveness - determine whether the trait presented is an appropriate description of the target person | trait inference (friend) > perceptual judgement (uppercase letters) | E>C | NA | Explicit |
| Kana et al., 2017 | fMRI | 7 | 15 | Trait descriptiveness - determine whether the trait presented is an appropriate description of the target person | trait inference (teacher) > lexical judgement | NR | NA | Explicit |

| | | | | | | | | |
|---|---|---|---|---|---|---|---|---|
| Kestemont et al., 2013 | fMRI | 10 | 17 | Causal attribution - Judge the degree to which the person is the cause of the event described | person attributions > semantic judgement | NR | NR | Explicit |
| Kestemont et al., 2013 | fMRI | 9 | 17 | Gender judgement and factual judgements | person attributions > factual statements | NA | NA | Implicit |
| Kestemont et al., 2015 | fMRI | 16 | 20 | Causal attribution - Attribute the event described to the person, the self or the situation | person attributions > semantic judgements | NR | NR | Explicit |
| Ma et al., 2011 | fMRI | 17 | 15 | Infer the personality traits of the agent based on the statement presented | trait inference > gender judgement | NR | NR | Explicit |
| Ma et al., 2011 | fMRI | 2 | 15 | Read the statements/ Gender judgement | trait diagnostic > non-diagnostic statements | NA | NA | Implicit |
| Ma et al., 2012 | fMRI | 6 | 14 | Infer the personality traits of the person depicted based on the photograph and the statement presented | trait inference > physical judgement | E>C | E=C | Explicit |
| | | | | Infer the personality traits of the person depicted based on the photograph presented | trait inference > physical judgement | E>C | E>C | Explicit |
| Mitchell et al., 2004 | fMRI | 13 | 17 | Infer the personality traits of the person depicted based on the statement presented | trait inference > remember statement order | NA | NA | Explicit |
| Mitchell et al., 2005 | fMRI | 9 | 14 | Form an opinion about the person/object depicted based on the statement presented | impression formation (person) > remember statement order/impression formation (objects) | NA | NA | Explicit |
| | | | | Form an opinion about the person/object depicted based on the statement presented | impression formation (person) > remember statement order | NA | NA | Explicit |
| Mitchell et al., 2006 | fMRI | 15 | 15 | Infer the personality traits of the person depicted based on the statement presented | trait inference > remember statement order (collapsed across statement diagnosticity) | NA | NA | Explicit |
| | | | | Infer the personality traits of the person depicted based on the statement presented/ Remember statement order | trait diagnostic > non-diagnostic statements (collapsed across tasks) | NA | NA | Explicit/Implicit |

| | | | | | | | | |
|---|---|---|---|---|---|---|---|---|
| Modinos et al., 2009 | fMRI | 7* | 16 | Trait descriptiveness - determine whether the trait presented is an appropriate description of the target person | trait inference (acquaintance) > semantic judgements | E=C | NA | Explicit |
| Moran et al., 2014 | fMRI | 27 | 16 | Causal attribution -Attribute the event described to the person or the situation | person > situation attributions | NR | NA | Explicit |
| | | | | Causal attribution -Attribute the event described to the person or the situation | person > situation attributions (ambiguous scenarios) | NR | NA | Explicit |
| Mukherjee et al., 2014 | fMRI | 5 | 24 | Judge whether the person depicted is approachable | Approachability > gender judgements | NR | NR | Explicit |
| Murphy et al., 2010 | fMRI | 6 | 10 | Trait descriptiveness - determine whether the trait presented is an appropriate description of the target person | trait inference (friend) > valence judgement | NR | NA | Explicit |
| Ochsner et al., 2005 | fMRI | 3 | 17 | Trait descriptiveness - determine whether the trait presented is an appropriate description of the target person | trait inference (close other) > syllable counting | E=C | NA | Explicit |
| Pauly et al., 2014 | fMRI | 3 | 13 | Trait descriptiveness - determine whether the trait presented is an appropriate description of the target person | trait inference (intimate other) > lexical judgement | E>C | NA | Explicit |
| Pinkham et al., 2008 | fMRI | 18* | 12 | Judge whether the person depicted is trustworthy | trait inference > age judgement | E>C | NA | Explicit |
| Schmitz et al., 2004 | fMRI | 3 | 18 | Trait descriptiveness - determine whether the trait presented is an appropriate description of the target person | trait inference (close other) > valence judgement | E=C | NA | Explicit |
| Singer et al., 2004 | fMRI | 5 | 11 | Gender judgement | trustworthy/untrustworthy > neutral faces | NA | NA | Implicit |
| Smith et al., 2016 | fMRI | 12 | 23 | Compare the faces depicted based on dominance | trait inference > perceptual judgement | E>C | E>C | Explicit |
| Stanfield et al., 2017 | fMRI | 5 | 33 | Judge whether the person depicted is approachable | Approachability > gender judgements | NR | NR | Explicit |

| | | | | | | | | |
|---|---|---|---|---|---|---|---|---|
| Sugiura et al., 2004 | PET | 46 | 9 | Trait descriptiveness - determine whether the trait presented is an appropriate description of the target person | trait inference (comic character) > colour judgement | E>C | E>C | Explicit |
| | | | | Trait descriptiveness - determine whether the trait presented is an appropriate description of the target person | trait inference (famous person) > colour judgement | E>C | E>C | Explicit |
| | | | | Trait descriptiveness - determine whether the trait presented is an appropriate description of the target person | trait inference (new friend) > colour judgement | E>C | E>C | Explicit |
| | | | | Trait descriptiveness - determine whether the trait presented is an appropriate description of the target person | trait inference (old friend) > colour judgement | E>C | E>C | Explicit |
| | | | | Trait descriptiveness - determine whether the trait presented is an appropriate description of the target person | trait inference (sibling) > colour judgement | E>C | E>C | Explicit |
| | | | | Trait descriptiveness - determine whether the trait presented is an appropriate description of the target person | trait inference (father) > colour judgement | E>C | E>C | Explicit |
| Turk et al., 2004 | fMRI | 7 | 18 | Preference judgement | date preference judgement > inconsequential choice | E>C | NA | Explicit |
| Zhu et al., 2007 | fMRI | 4 | 13 | Trait descriptiveness - determine whether the trait presented is an appropriate description of the target person | trait inference (mother) > perceptual judgement (uppercase letters) | NR | NA | Explicit |
| | | | | Trait descriptiveness - determine whether the trait presented is an appropriate description of the target person | trait inference (other) > perceptual judgement (uppercase letters) | NR | NA | |
| Zhu et al., 2007 | fMRI | 11 | 13 | Trait descriptiveness - determine whether the trait presented is an appropriate description of the target person | trait inference (mother) > perceptual judgement (uppercase letters) | NR | NA | Explicit |

| | | | | | |
|---|---|---|---|---|---|
| | Trait descriptiveness - determine whether the trait presented is an appropriate description of the target person | trait inference (other) > perceptual judgement (uppercase letters) | NR | NA | Explicit |

* the coordinates were obtained via personal communication with the authors.
N = sample size; E>C - the experimental condition was more difficult than the control condition; C>E - the control condition was more difficult than the experimental condition; E=C - the experimental and control conditions were equally difficult; NR - not reported; NA - not applicable because no behavioural data was collected.

**Table S3.** List of studies included in the **empathy** (N = 164) meta-analysis.

| Authors | Imaging method | No. of peaks | N | Task | Contrast | Difficulty RT | Difficulty Accuracy | Instructional cue | Target of empathy |
|---|---|---|---|---|---|---|---|---|---|
| Akitsuki & Decety, 2009 | fMRI | 37 | 26 | Passive observation | painful > non-painful situation | | | Implicit | Pain |
| Azevedo et al., 2013 | fMRI | 15 | 27 | Passive observation | pain infliction > non-painful touch | | | Implicit | Pain |
| Azevedo et al., 2014 | fMRI | 12 | 12 | Passive observation | pain infliction > non-painful touch | | | Implicit | Pain |
| Bagozzi et al., 2013 | fMRI | 9 | 24 | Passive observation | negative emotional facial expression > moving geometric shapes | | | Implicit | Emotion |
| | | | | Passive observation | positive emotional facial expression > moving geometric shapes | | | Implicit | Emotion |
| Ballotta et al., 2018 | fMRI | 2 | 30 | Dual task: gender judgement and temporal production | painful > neutral facial expression | | | Implicit | Pain |
| Benuzzi et al., 2008 | fMRI | 14 | 15 | Rate the intensity of the perceived unpleasantness of the stimulus | pain infliction > non-painful touch | NR | NA | Explicit | Pain |
| | | | | Rate the intensity of the perceived unpleasantness of the stimulus | disgusting > neutral stimulation | | | Explicit | Emotion |
| Berlingeri et al., 2016 | fMRI | 20 | 25 | Rate the intensity of pain experienced by the depicted person | pain infliction > non-painful touch (stimulus presentation) | E>C | NA | Explicit | Pain |
| | | | | Rate the intensity of pain experienced by the depicted person | pain infliction > non-painful touch (response phase) | E>C | NA | Explicit | Pain |
| Bos et al., 2015 | fMRI | 12 | 24 | Attentional task - indicate when the fixation cross changes colour | pain infliction > non-painful touch | | | Implicit | Pain |
| Botvinick et al., 2005 | fMRI | 17 | 12 | Passive observation | painful > neutral facial expression | | | Implicit | Pain |

| | | | | | | | | | |
|---|---|---|---|---|---|---|---|---|---|
| Braboszcz et al., 2017 | fMRI | 9 | 17 | Go/No-go task | painful > non-painful situations | | | Implicit | Pain |
| Bruneau et al., 2012 | fMRI | 10 | 44 | Rate the intensity of compassion felt for the protagonist | physical pain > neutral | NR | NA | Explicit | Pain |
| | | | | Rate the intensity of compassion felt for the protagonist | emotional pain > neutral | NR | NA | Explicit | Emotion |
| Bruneau et al., 2012 | fMRI | 10 | 41 | Rate the intensity of compassion felt for the protagonist | physical pain > neutral | E=C | NA | Explicit | Pain |
| | | | | Rate the intensity of compassion felt for the protagonist | emotional pain > neutral | E=C | NA | Explicit | Emotion |
| Brunnlieb et al., 2013 | fMRI | 18 | 18 | Imagine how you would feel in the depicted situation | emotional/pain > neutral situations | NA | NA | Explicit | Pain/Emotion |
| Budell et al., 2010 | fMRI | 41 | 18 | Rate the intensity of pain experienced by the depicted person | painful > neutral facial expression | NR | NA | Explicit | Pain |
| | | | | Perceptual judgement | painful > neutral facial expression | | | Implicit | Pain |
| Budell et al., 2015 | fMRI | 67 | 23 | Observe then imitate the intensity of pain depicted | painful > neutral facial expressions (during observation) | NR | NR | Explicit | Pain |
| | | | | Observe then imitate the movement depicted | painful > neutral facial expressions (during observation) | | | Implicit | Pain |
| Cao et al., 2015 | fMRI | 5 | 30 | Rate the intensity of pain experienced by the depicted person | pain infliction > non-painful touch | NR | NA | Explicit | Pain |
| Cao et al., 2019 | fMRI | 11 | 29 | Rate the intensity of pain experienced by the depicted person/ Rate your own experience of discomfort | pain infliction > non-painful touch | NR | NA | Explicit | Pain |
| Cheng et al., 2007 | fMRI | 17 | 14 | Attentional task - report the number of video interruptions | pain infliction > non-painful touch | | | Implicit | Pain |
| | | | | | pain infliction > rest | | | Implicit | Pain |

| | | | | | | | | | |
|---|---|---|---|---|---|---|---|---|---|
| | | | | Attentional task - report the number of video interruptions | | | | | |
| Cheng et al., 2007 | fMRI | 9 | 14 | Attentional task - report the number of video interruptions | pain infliction > non-painful touch | | | Implicit | Pain |
| | | | | Attentional task - report the number of video interruptions | pain infliction > rest | | | Implicit | Pain |
| Cheng et al., 2010 | fMRI | 14 | 36 | Imagine the scenario from the perspective of the self/a loved one/a stranger | painful > non-painful situations | NR | NA | Explicit | Pain |
| Cheon et al., 2013 | fMRI | 19 | 27 | Attentional task - indicate stimulus onset | emotionally painful > neutral situation | | | Implicit | Emotion |
| Chiao et al., 2009 | fMRI | 15 | 14 | Rate the intensity of empathic concern experienced | emotionally painful > neutral situation | E>C | NA | Explicit | Emotion |
| Christov-Moore & Iacoboni, 2019 | fMRI | 29 | 70 | Passive observation | pain infliction > non-painful touch | | | Implicit | Pain |
| Christov-Moore et al., 2017 | fMRI | 35 | 19 | Passive observation | pain infliction > non-painful touch | | | Implicit | Pain |
| Cogoni et al., 2018 | fMRI | 15 | 36 | Cyberball task adaptation - Rate the valence of other's emotion | social exclusion > inclusion | NR | NA | Explicit | Emotion |
| Coll et al., 2017 | fMRI | 9 | 15 | Indicate how much help they would offer the patient presented in the photo | painful > neutral facial expressions | NR | NA | Explicit | Pain |
| Contreras-Huerta et al., 2013 | fMRI | 10 | 20 | Rate the intensity of pain experienced by the depicted person | painful > non-painful situations | NR | NA | Explicit | Pain |
| Corradi-Dell'Acqua et al., 2019 | fMRI | 19 | 33 | Handedness judgement | painful > non-painful situations | | | Implicit | Pain |

| | | | | | | | | | |
|---|---|---|---|---|---|---|---|---|---|
| Corradi-Dell'Acqua et al., 2011 | fMRI | 29 | 28 | Handedness judgement | painful > neutral non-painful situations | | | Implicit | Pain |
| | | | | Handedness judgement | painful > negative non-painful situations | | | Implicit | Pain |
| Danziger et al., 2009 | fMRI | 23 | 13 | Imagine how the depicted person feels | painful > non-painful situations | NA | NA | Explicit | Pain |
| | | | | Imagine how the depicted person feels | painful > neutral facial expressions | NA | NA | Explicit | Pain |
| Dapretto et al., 2005 | fMRI | 8 | 10 | Passive observation | emotional facial expressions > rest | | | Implicit | Emotion |
| de Gelder et al., 2004 | fMRI | 45 | 7 | Passive observation | fearful > neutral whole-body expressions | | | Implicit | Emotion |
| | | | | Passive observation | happy > neutral whole-body expressions | | | Implicit | Emotion |
| de Greck et al., 2012 | fMRI | 17 | 20 | Rate the ability to share the emotional state of the person depicted | emotional facial expressions > scrambled stimuli | NA | NA | Explicit | Emotion |
| de Greck et al., 2012 | fMRI | 28 | 20 | Rate your ability to empathize with the depicted person/ Skin colour evaluation | empathy > perceptual judgements | C>E | NA | Explicit | Emotion |
| | | | | Rate your ability to empathize with the depicted person | empathy > rest | NA | NA | Explicit | Emotion |
| Decety & Michalska, 2010 | fMRI | 10 | 57 | Passive observation | painful > non-painful situations | | | Implicit | Pain |
| Decety et al., 2008 | fMRI | 11 | 17 | Passive observation | painful > non-painful situations | | | Implicit | Pain |
| Decety et al., 2009 | fMRI | 9 | 8 | Passive observation | painful > non-painful situations | | | Implicit | Pain |
| Deeley et al., 2006 | fMRI | 14 | 9 | Gender judgement | happy > neutral facial expressions | | | Implicit | Emotion |
| | | | | Gender judgement | fearful > neutral facial expressions | | | Implicit | Emotion |

| | | | | | | | | | |
|---|---|---|---|---|---|---|---|---|---|
| Dong et al., 2017 | fMRI | 13 | 36 | Judge whether the depicted person was experiencing pain | pain infliction > non-painful touch & facial expression | NR | NR | Explicit | Pain |
| Enzi et al., 2016 | fMRI | 10 | 20 | Empathize with the depicted person | painful > non-painful situation | NR | E=C | Explicit | Pain |
| Ernst et al., 2013 | fMRI | 5 | 18 | Indicate whether they were able to share the emotional state of the person depicted | emotional facial expressions > blurred photos | E=C | NA | Explicit | Emotion |
| Fan et al., 2014 | fMRI | 16 | 21 | Passive observation | painful > non-painful situations | | | Implicit | Pain |
| Feng et al., 2016 | fMRI | 34 | 22 | Empathize with and judge the intensity of pain experienced by the depicted person | pain infliction > non-painful touch & facial expression | NR | NA | Explicit | Pain |
| Flasbeck et al., 2019 | fMRI | 30* | 19 | Empathize with the person depicted | pain infliction > non-painful touch | NR | NA | Explicit | Pain |
| | | | | Empathize with the person depicted | pain infliction > non-painful touch | | | Explicit | Pain |
| Fourie et al., 2017 | fMRI | 37 | 38 | Empathize with person depicted | painful > neutral facial expressions | NR | NA | Explicit | Pain |
| | | | | Empathize with person depicted | emotionally distressing > neutral situations | | | Explicit | Emotion |
| Fourie et al., 2019 | fMRI | 42 | 36 | Empathize with the protagonist | forgiving victim > neutral | NR | NA | Explicit | Emotion |
| | | | | Empathize with the protagonist | unforgiving victim > neutral | | | Explicit | Emotion |
| | | | | Empathize with the protagonist | apologetic perpetrator > neutral | | | Explicit | Emotion |
| | | | | Empathize with the protagonist | unapologetic perpetrator > neutral | | | Explicit | Emotion |
| Fujino et al., 2014 | fMRI | 12 | 11 | Rate the intensity of pain experienced by the depicted person | painful > non-painful situations | NR | NA | Explicit | Pain |
| Gao et al., 2017 | fMRI | 8 | 35 | Judge whether the depicted person was experiencing pain | painful > non-painful situations | NR | NR | Explicit | Pain |
| Geday et al., 2003 | PET | 1 | 8 | Passive observation | emotional > neutral facial expressions/situations | | | Implicit | Emotion |

| | | | | | | | | | |
|---|---|---|---|---|---|---|---|---|---|
| Geday et al., 2007 | PET | 1 | 12 | Passive observation | emotional > neutral facial expressions/situations | | | Implicit | Emotion |
| Gottlich et al., 2017 | fMRI | 18 | 26 | Passive observation | emotional > neutral situations | | | Implicit | Pain/Emotion |
| Greimel et al., 2010 | fMRI | 39 | 47 | Emotion categorization | Emotion categorization > perceptual judgement (face width) | NR | NR | Explicit | Emotion |
| | | | | Rate your own emotional response to the depicted face | empathic > perceptual judgement (face width) | NR | NR | Explicit | Emotion |
| Grice-Jackson et a., 2017 | fMRI | 4 | 21 | Judge whether you experienced a painful sensation while observing the depicted person | painful > non-painful situations | NR | NA | Explicit | Pain |
| Grice-Jackson et a., 2017 | fMRI | 14 | 13 | Judge whether you experienced a painful sensation while observing the depicted person | painful > non-painful situations | NR | NA | Explicit | Pain |
| Grice-Jackson et a., 2017 | fMRI | 11 | 10 | Judge whether you experienced a painful sensation while observing the depicted person | painful > non-painful situations | NR | NA | Explicit | Pain |
| Grosbras & Paus, 2006 | fMRI | 75 | 20 | Passive observation | angry facial expression > non-biological motion control | | | Implicit | Emotion |
| | | | | Passive observation | angry hand movement > non-biological motion control | | | Implicit | Emotion |
| | | | | Passive observation | angry > neutral hand movement | | | Implicit | Emotion |
| Gu & Han, 2007 | fMRI | 13 | 12 | Rate the intensity of pain experienced by the person depicted (painful situations)/ Count the number of hands depicted (non-painful situations) | painful > non-painful situations | NR | NR | Explicit | Pain |
| | | | | Rate the intensity of pain experienced by the person depicted (painful situations)/ Count the number of hands | painful > non-painful situations | | | Explicit | Pain |

| | | | | | | | | | |
|---|---|---|---|---|---|---|---|---|---|
| | | | | depicted (non-painful situations) | | | | | |
| Gu et al., 2010 | fMRI | 32 | 18 | Judge whether the depicted person was experiencing pain | painful > non-painful situations | E=C | E>C | Explicit | Pain |
| | | | | Handedness judgement | painful > non-painful situations | | | Implicit | Pain |
| Gu et al., 2013 | fMRI | 89 | 18 | Judge whether the depicted person was experiencing pain | painful > non-painful situations | E=C | E=C | Explicit | Pain |
| | | | | Judge whether the depicted person was experiencing pain | pain judgement > rest | NA | NA | Explicit | Pain |
| | | | | Body part identification/ Laterality judgement | painful > non-painful situations | | | Implicit | Pain |
| Guo et al., 2012 | fMRI | 16 | 16 | Empathize with the depicted person | painful > non-painful situations | NR | NA | Explicit | Pain |
| Guo et al., 2013 | fMRI | 10 | 40 | Empathize with the depicted person | painful > non-painful situations | NR | NA | Explicit | Pain |
| Hadjikhani et al., 2014 | fMRI | 41 | 31 | Attentional task - indicate when the fixation cross changed colour | painful > neutral facial expressions | | | Implicit | Pain |
| Han et al., 2009 | fMRI | 3 | 24 | Judge whether the depicted person was experiencing pain | pain infliction > non-painful touch | NR | NA | Explicit | Pain |
| Han et al., 2009 | fMRI | 6 | 22 | Judge whether the depicted person was experiencing pain | pain infliction > non-painful touch | NR | NA | Explicit | Pain |
| | | | | Judge whether the depicted person was experiencing pain | painful > neutral facial expressions | NR | NA | Explicit | Pain |
| Han et al., 2017 | fMRI | 21 | 33 | Passive observation | pain infliction > non-painful stimulation & facial expressions | | | Implicit | Pain |
| | | | | Press a button throughout stimulus presentation | pain infliction > non-painful stimulation & facial expressions | | | Implicit | Pain |
| Hennenlotter et al., 2005 | fMRI | 20 | 12 | Passive observation | emotional > neutral facial expressions | | | Implicit | Emotion |
| Horan et al., 2014 | fMRI | 4 | 22 | Passive observation | emotional facial expressions > rest | | | Implicit | Emotion |

| | | | | | | | | | |
|---|---|---|---|---|---|---|---|---|---|
| Jackson et al., 2005 | fMRI | 15 | 15 | Rate the intensity of pain experienced by the person depicted | painful > non-painful situations | NR | NA | Explicit | Pain |
| Jackson et al., 2017 | fMRI | 8 | 24 | Rate the intensity of pain experienced by depicted infant | painful > non-painful situation | NR | NA | Explicit | Pain |
| | | | | Rate the intensity of pain experienced by the depicted adult | painful > non-painful situation | NR | NA | Explicit | Pain |
| Jackson et al., 2017 | fMRI | 2 | 27 | Rate the intensity of pain experienced by the depicted adult | painful > non-painful situation | NR | NA | Explicit | Pain |
| Jankowiak-Siuda et al., 2015 | fMRI | 20 | 27 | Rate the intensity of compassion felt for the depicted person | painful > non-painful situation | NR | NA | Explicit | Pain |
| Jensen et al., 2014 | fMRI | 2 | 18 | Rate your own satisfaction | pain > no pain | NR | NA | Explicit | Pain |
| Kana et al., 2016 | fMRI | 18 | 15 | Emotion categorization | emotional > neutral scene | E>C | NR | Explicit | Emotion |
| | | | | Identify the blurred object | emotional > neutral scene | | | Implicit | Emotion |
| Kanske et al., 2015 | fMRI | 44 | 25 | EmpaToM task - Rate the valence of your own emotional state and the intensity of compassion felt | emotional > neutral facial expressions | NR | NA | Explicit | Emotion |
| | | | | Rate the valence of your own emotional state and the intensity of compassion felt | emotional > neutral situations | NR | NA | Explicit | Emotion |
| Kanske et al., 2015 | fMRI | 23 | 178 | EmpaToM task - Rate the valence of your own emotional state and the intensity of compassion felt | emotional > neutral facial expressions | NR | NA | Explicit | Emotion |
| Kim et al., 2009 | fMRI | 21 | 21 | Observe the depicted person with a compassionate attitude | emotional > neutral facial expression | NA | NA | Explicit | Emotion |
| | | | | Observe the depicted person with a compassionate attitude/Passive observation | compassion > passive observation | NA | NA | Explicit | Emotion |

| | | | | | | | | | |
|---|---|---|---|---|---|---|---|---|---|
| Kim et al., 2010 | fMRI | 7 | 19 | Infer what will improve the emotional state of the protagonist | emotional inference > physical causality | NR | C >E | Explicit | Emotion |
| Klimecki et al., 2012 | fMRI | 23 | 94 | Rate the intensity of compassion felt for the depicted person and the valence of your own emotional reaction | emotional > neutral situations | NR | NA | Explicit | Emotion |
| Krach et al., 2015 | fMRI | 28 | 16 | Rate the intensity of physical pain experienced by the depicted person | painful > non-painful situation | NR | NR | Explicit | Pain |
| | | | | Rate the intensity of vicarious embarrassment experienced | embarrassing > neutral scenarios | NR | NR | Explicit | Emotion |
| Kramer et al., 2010 | fMRI | 6 | 16 | Passive observation | emotional/painful > neutral situations | | | Implicit | Pain/Emotion |
| Labek et al., 2017 | fMRI | 14 | 17 | Passive observation | emotional > neutral | | | Implicit | Emotion |
| Lamm & Decety, 2008 | fMRI | 33 | 18 | Rate the intensity of pain experienced by the depicted person | pain infliction > no pain | NR | NA | Explicit | Pain |
| Lang et al., 2011 | fMRI | 11 | 22 | Passive listening | affective > neutral human sounds | | | Implicit | Pain/Emotion |
| Lassalle et al., 2019 | fMRI | 88 | 20 | Categorize the type of stimulation (painful/disgusting/neutral) | pain infliction > non-painful touch | E=C | E=C | Explicit | pain |
| | | | | Categorize the type of stimulation (painful/disgusting/neutral) | disgusting > neutral stimulation | E=C | E=C | Explicit | Emotion |
| Lee et al., 2006 | fMRI | 18 | 14 | Empathic judgements about emotional scenario / Social judgements about non-emotional scenarios | emotional > neutral scenarios (time point 1) | E>C | NA | Explicit | Emotion |
| | | | | Empathic judgements about emotional scenarios/ Social judgements about non-emotional scenarios | emotional > neutral scenarios (time point 2) | E>C | NA | Explicit | Emotion |
| Lee et al., 2010 | fMRI | 15 | 18 | Story ending judgement | empathic inference > physical causality judgement | NR | NR | Explicit | Pain/Emotion |

| | | | | | | | | | |
|---|---|---|---|---|---|---|---|---|---|
| Lee et al., 2013 | fMRI | 2 | 12 | Passive observation | emotional > neutral facial expressions | | | Implicit | Emotion |
| Leiberg et al., 2012 | fMRI | 22 | 24 | Passive observation (pre-regulation) | emotional > neutral situations | | | Implicit | Emotion |
| | | | | Empathize with the depicted person/ Passive observation | empathize > passive observation | NA | NA | Explicit | Emotion |
| Li et al., 2015 | fMRI | 2 | 40 | Judge whether the depicted person was experiencing pain | painful > neutral facial expressions | C>E | NR | Explicit | Pain |
| Luo et al., 2014 | fMRI | 11 | 36 | Judge whether the depicted person was experiencing pain | pain infliction > non-painful touch | NR | NR | Explicit | Pain |
| Luo et al., 2015 | fMRI | 18 | 30 | Judge whether the depicted person (Asian) was experiencing pain | pain infliction > non-painful touch | NR | NR | Explicit | Pain |
| | | | | Judge whether the depicted person (Caucasian) was experiencing pain | pain infliction > non-painful touch | NR | NR | Explicit | Pain |
| Luo et al., 2015 | fMRI | 18 | 30 | Judge whether the depicted person (Asian) was experiencing pain | pain infliction > non-painful touch | NR | NR | Explicit | Pain |
| | | | | Judge whether the depicted person (Caucasian) was experiencing pain | pain infliction > non-painful touch | NR | NR | Explicit | Pain |
| Ma et al., 2011 | fMRI | 5 | 33 | Judge whether the depicted person was experiencing pain | pain infliction > non-painful stimulation & facial expressions | NR | NR | Explicit | Pain |
| Mackes et al., 2018 | fMRI | 17 | 34 | Categorize and rate the intensity of the emotion experienced by the depicted person and rate your own emotional reaction | emotional > neutral | NR | NR | Explicit | Emotion |
| Mascaro et al., 2014 | fMRI | 5 | 36 | Passive listening | affective human sounds > neutral tones | | | Implicit | Emotion |
| Mathur et al., 2010 | fMRI | 13 | 28 | Rate the intensity of empathic feelings elicited by the stimuli | emotional > neutral situations | E>C | NA | Explicit | Emotion |

| | | | | | | | | | |
|---|---|---|---|---|---|---|---|---|---|
| Mathur et al., 2016 | fMRI | 21 | 15 | Rate the intensity of empathic feelings elicited by the stimuli | emotional > neutral situations | E>C | NA | Explicit | Emotion |
| Mazza et al., 2013 | fMRI | 8 | 10 | Passive observation | emotional/painful situations > scrambled photos | | | Implicit | Pain/Emotion |
| Mazzola et al., 2010 | fMRI | 7 | 30 | Judge the degree of familiarity with the person depicted | painful > neutral facial expressions | | | Implicit | Pain |
| Melchers et al., 2015 | fMRI | 6 | 60 | Passive observation | embarrassing > neutral situations | | | Implicit | Emotion |
| Mercadillo et al., 2011 | fMRI | 23 | 24 | Judge whether the stimuli elicited compassionate feelings | emotional > neutral social scenes | NR | NA | Explicit | Emotion |
| | | | | Judge whether the stimuli elicited compassionate feelings | emotional > neutral non-social scenes | NR | NA | Explicit | Emotion |
| Mercadillo et al., 2015 | fMRI | 14 | 24 | Judge whether the stimuli elicited compassionate feelings | emotional > neutral social scenes | NR | NA | Explicit | Emotion |
| | | | | Judge whether the stimuli elicited compassionate feelings | emotional > neutral non-social scenes | NR | NA | Explicit | Emotion |
| Michalska et al., 2013 | fMRI | 11 | 65 | Passive observation | painful > non-painful situation | | | Implicit | Pain |
| Molenberghs et al., 2016 | fMRI | 17 | 48 | Passive observation | painful > non-painful situation | | | Implicit | Pain |
| Moll et al., 2007 | fMRI | 11 | 12 | Imagine you are the protagonist | compassion-eliciting > neutral scenarios | NR | NA | Explicit | Emotion |
| Montoya et al., 2012 | fMRI | 22 | 17 | One-back memory task | affective human sounds (low distress) > noise | | | Implicit | Emotion |
| | | | | One-back memory task | affective human sounds (high distress) > noise | | | Implicit | Emotion |
| | | | | One-back memory task | emotional > neutral facial expressions | | | Implicit | Emotion |
| | | | | One-back memory task | emotional > neutral facial expressions | | | Implicit | Emotion |

| | | | | | | | | | |
|---|---|---|---|---|---|---|---|---|---|
| Moore et al., 2015 | fMRI | 5 | 27 | Match facial emotional expressions | emotional faces > geometric shapes | NR | NR | Explicit | Emotion |
| Morelli et al., 2014 | fMRI | 32 | 32 | Imagine how intense the pain experienced by the depicted person is | painful > non-painful situations | NA | NA | Explicit | Pain |
| | | | | Imagine how you would feel in the depicted situation | emotional > neutral situation | NA | NA | Explicit | Emotion |
| | | | | Imagine how you would feel in the depicted situation | emotional > neutral situation | NA | NA | Explicit | Emotion |
| Moriguchi et al., 2007 | fMRI | 25 | 14 | Rate the intensity of pain experienced by the depicted person | painful > non-painful situations | NR | NA | Explicit | Pain |
| Morrison & Downing, 2007 | fMRI | 5 | 11 | Attentional task - indicate when the target body part was stimulated | pain infliction > non-painful touch | | | Implicit | Pain |
| Morrison et al., 2004 | fMRI | 4 | 14 | Passive observation | pain infliction > rest | | | Implicit | Pain |
| Morrison et al., 2013 | fMRI | 12 | 14 | Evaluate whether the depicted action was appropriate given noxious/innocuous stimulus | painful > non-painful action | | | Implicit | Pain |
| Noll-Hussong et al., 2013 | fMRI | 13 | 19 | Passive observation | painful > non-painful situations | | | Implicit | Pain |
| | | | | Passive observation | painful situations > rest | | | Implicit | Pain |
| Nomi et al., 2008 | fMRI | 12 | 14 | Emotion categorization/ Empathize with the depicted person/ Perceptual judgement | emotional facial expressions > scrambled stimuli | NA | NA | Explicit/Implicit | Emotion |
| Nummenmaa et al., 2008 | fMRI | 6 | 10 | Empathize with the target person | emotional > neutral scenes | NA | NA | Explicit | Emotion |
| Oliver et al., 2018 | fMRI | 14 | 36 | Rate your own emotional reaction to the people depicted | empathic > age judgement | NR | NR | Explicit | Emotion |
| Osborn & Derbyshire, 2010 | fMRI | 21* | 10 | Rate the intensity of vicarious painful sensations experienced and the unpleasantness of the stimuli | painful > non-painful situations | NR | NA | Explicit | Pain |

| | | | | | | | | | |
|---|---|---|---|---|---|---|---|---|---|
| Osborn & Derbyshire, 2010 | fMRI | 2* | 10 | Rate the intensity of vicarious painful sensations experienced and the unpleasantness of the stimuli | painful > non-painful situations | NA | NA | Explicit | Pain |
| Paulus et al., 2015 | fMRI | 16 | 32 | Rate the intensity of vicarious embarrassment experienced | embarrassing > neutral scenarios | E>C | NA | Explicit | Emotion |
| Paulus et al., 2018 | fMRI | 13 | 17 | Rate the intensity of vicarious embarrassment experienced | embarrassing > neutral scenarios | NR | NA | Explicit | Emotion |
| Pehrs et al., 2015 | fMRI | 2 | 26 | Rate the intensity of compassion felt for the protagonist | emotional > neutral context | NR | NA | Explicit | Emotion |
| Powell et al., 2017 | fMRI | 12 | 12 | Infer what will improve the emotional state of the protagonist | emotional inference > physical causality | NR | NR | Explicit | Emotion |
| Preis et al., 2013 | fMRI | 9 | 64 | Rate the intensity of pain experienced by the depicted person | painful > non-painful situations | NR | NA | Explicit | Pain |
| Preston et al., 2007 | PET | 18 | 16 | Imagine the scenario and focus on the emotional experience | emotional > neutral scenarios | NA | NA | Explicit | Emotion |
| Prochnow et al., 2013 | fMRI | 20 | 15 | Emotion categorization and Press alternating buttons | emotional expressions > scrambled stimuli | NA | NA | Explicit/Implicit | Emotion |
| | | | | Emotion categorization | emotion categorization > motor control (observation phase) | NA | NA | Explicit | Emotion |
| | | | | Emotion categorization and Press alternating buttons | emotional expressions > scrambled stimuli (observation phase) | NA | NA | Explicit/Implicit | Emotion |
| | | | | Emotion categorization | emotion categorization > motor control (observation phase) | NA | NA | Explicit | Emotion |
| Prochnow et al., 2014 | fMRI | 20 | 26 | Match descriptive sentences to facial expressions | emotional expressions > scrambled stimuli | NA | NA | Explicit | Emotion |
| | | | | Match descriptive sentences to facial expressions | emotional expressions > scrambled stimuli (response phase) | NA | NA | Explicit | Emotion |

| | | | | | | | | | |
|---|---|---|---|---|---|---|---|---|---|
| Qiao-Tasserit et al., 2018 | fMRI | 13 | 24 | Handedness judgement following positive context | pain > no-pain | | | Implicit | Pain |
| | | | | Handedness judgement following neutral context | pain > no-pain | | | Implicit | Pain |
| | | | | Handedness judgement following negative context | pain > no-pain | | | Implicit | Pain |
| Regenbogen et al., 2012 | fMRI | 24 | 27 | Rate the valence of the emotions experienced by the depicted person and the self | emotional > neutral communication | NR | NA | Explicit | Emotion |
| Reniers et al., 2014 | fMRI | 13 | 15 | Imagine what the person depicted is feeling | emotional > neutral situations | NA | NA | Explicit | Emotion |
| Richins et al., 2019 | fMRI | 19 | 69 | Rate the intensity of pain experienced by the depicted person and rate your own emotional reaction to the stimuli | painful > neutral stimulation & facial expression | NR | NA | Explicit | Pain |
| Riekki et al., 2018 | fMRI | 24 | 38 | Passive observation | emotional > neutral situations | | | Implicit | Pain/Emotion |
| | | | | Rate the stimuli on how emotionally touching they are/ how easily recognizable the facial expressions are | emotional > neutral situations | NR | NA | Explicit | Pain/Emotion |
| Riem et al., 2011 | fMRI | 2 | 21 | Passive listening | affective human sounds > neutral sounds | | | Implicit | Emotion |
| Ruckmann et al., 2015 | fMRI | 11 | 30 | Rate the intensity of pain experienced by the depicted person | painful > non-painful situations | NR | NA | Explicit | Pain |
| Rutgen et al., 2019 | fMRI | 5 | 35 | Rate the unpleasantness of the stimulation and own emotional reaction to the stimuli | painful stimulation & facial expression > rest | NA | NA | Explicit | Pain |
| Rymarczyk et al., 2019 | fMRI | 66 | 46 | Passive observation | emotional > neutral facial expressions | | | Implicit | Emotion |
| | | | | Passive observation | emotional > neutral facial expressions | | | Implicit | Emotion |

| | | | | | | | | | |
|---|---|---|---|---|---|---|---|---|---|
| Schulte-Ruther et al., 2008 | fMRI | 24 | 26 | Evaluate your own emotional reaction to the emotional facial expressions/ Gender/age judgements of neutral facial expressions | emotional > neutral facial expressions | E>C | NA | Explicit | Emotion |
| | | | | Emotion categorization of emotional facial expressions/ Gender/age judgements of neutral facial expressions | emotional > neutral facial expressions | E>C | NR | Explicit | Emotion |
| Seara-Cardoso et al., 2015 | fMRI | 15 | 46 | Body part identification | painful > non-painful situations | | | Implicit | Pain |
| Seara-Cardoso et al., 2016 | fMRI | 2 | 30 | Rate your emotional reaction to the stimuli | emotional > neutral facial expressions | NR | NA | Explicit | Emotion |
| | | | | Rate your emotional reaction to the stimuli | emotional > neutral facial expressions | | | Explicit | Emotion |
| Sheng et al., 2014 | fMRI | 13 | 21 | Judge whether the depicted person was experiencing pain | painful > neutral facial expressions | E>C | E=C | Explicit | Pain |
| | | | | Judge whether the depicted person was experiencing pain/ Race judgement | pain > race judgement | E=C | E=C | Explicit | Pain |
| Simon-Thomas et al., 2012 | fMRI | 1 | 17 | Rate the intensity of your emotional reaction to the stimuli | emotional > neutral scenes | NR | NA | Explicit | Pain/Emotion |
| Singer et al., 2004 | fMRI | 13 | 16 | Passive observation | pain > no pain | | | Implicit | Pain |
| Singh et al., 2015 | fMRI | 14 | 14 | Focus on your emotional reaction to the stimuli | emotional scenes > scrambled stimuli | NA | NA | Explicit | Emotion |
| Spencer et al., 2011 | fMRI | 12 | 40 | Gender judgement | emotional > neutral facial expressions | | | Implicit | Emotion |
| | | | | Gender judgement | emotional > neutral facial expressions | | | Implicit | Emotion |
| Stroth et al., 2019 | fMRI | 5 | 9 | Rate the intensity of pain experienced by the depicted person | painful > non-painful situations | NR | NA | Explicit | Pain |

| | | | | | | | | | |
|---|---|---|---|---|---|---|---|---|---|
| | | | | Rate the intensity of embarrassment experienced by the protagonist | embarrassment > no embarrassment | NR | NA | Explicit | Emotion |
| Szycik et al., 2017 | fMRI | 8 | 30 | Imagine how you would feel in the depicted situation | emotional > neutral scenarios | NA | NA | Explicit | Pain/Emotion |
| Tamm et al., 2017 | fMRI | 40 | 71 | Rate the intensity of your negative emotional reaction<br>Rate the intensity of your negative emotional reaction | pain infliction > non-painful stimulation & facial expressions<br>pain infliction > rest | NR<br>NA | NA<br>NA | Explicit<br>Explicit | Pain<br>Pain |
| Tholen et al., 2020 | fMRI | 20 | 130 | EmpaToM task - Rate the valence of your own emotional state and the intensity of compassion felt | emotional > neutral facial expressions | NR | NA | Explicit | Emotion |
| Toller et al., 2015 | fMRI | 14 | 30 | Passive observation | emotional facial expressions > neutral non-social scenes | | | Implicit | Emotion |
| Tousignant et al., 2018 | fMRI | 9 | 40 | Cyberball task adaptation - passive observation | social exclusion > inclusion | | | Implicit | Emotion |
| Tusche et al., 2016 | fMRI | 3 | 31 | EmpaToM task - Rate the valence of your own emotional state and the intensity of compassion felt | emotional > neutral facial expressions | NR | NA | Explicit | Emotion |
| Ushida et al., 2008 | fMRI | 14 | 15 | Passive observation | pain infliction > non-painful situation | | | Implicit | Pain |
| Ushida et al., 2008 | fMRI | 6 | 15 | Passive observation | pain infliction > non-painful situation | | | Implicit | Pain |
| Vachon-Presseau et al., 2012 | fMRI | 2 | 20 | Judge whether the depicted person was experiencing pain | painful > non-painful situations/facial expressions | NR | NR | Explicit | Pain |
| van der Heiden et al., 2013 | fMRI | 1 | 18 | Passive observation | painful > non-painful situations | | | Implicit | Pain |
| Vaughn et al., 2018 | fMRI | 6 | 105 | Passive observation | pain infliction > non-painful touch | | | Implicit | Pain |
| Vistoli et al., 2016 | fMRI | 11 | 21 | Rate the intensity of pain yourself/another person would experience in the situation depicted | painful > non-painful situations | E>C | NA | Explicit | Pain |

| | | | | | | | | | |
|---|---|---|---|---|---|---|---|---|---|
| Vollm et al., 2006 | fMRI | 9 | 13 | Infer what will improve the emotional state of the protagonist | empathic inference > physical causality | NR | C>E | Explicit | Emotion |
| Wang et al., 2015 | fMRI | 5 | 56 | Infer what will improve the emotional state of the protagonist | empathic inference > physical causality | NR | NR | Explicit | Emotion |
| Wickere et al., 2003 | fMRI | 23 | 14 | Passive observation<br>Passive observation | emotional > neutral facial expressions<br>emotional > neutral facial expressions | | | Implicit<br>Implicit | Emotion<br>Emotion |
| Xu et al., 2009 | fMRI | 4 | 16 | Judge whether the depicted person was experiencing pain | pain infliction > non-painful touch | NR | NR | Explicit | Pain |
| Xu et al., 2009 | fMRI | 3 | 17 | Judge whether the depicted person was experiencing pain | pain infliction > non-painful touch | NR | NR | Explicit | Pain |
| Zhao et al., 2020 | fMRI | 7 | 24 | Rate the intensity of pain experienced by the depicted person and your own experience of negative emotions | pain infliction > no pain | NR | NA | Explicit | Pain |
| Zheng et al., 2015 | fMRI | 6 | 20 | Empathize with the depicted person | painful > non-painful situations | NA | NA | Explicit | Pain |
| Zheng et al., 2016 | fMRI | 8 | 20 | Passive observation | painful > non-painful situations | | | Implicit | Pain |

*the coordinates were obtained via personal communication with the authors.
N = sample size; E>C - the experimental condition was more difficult than the control condition; C>E - the control condition was more difficult than the experimental condition; E=C - the experimental and control conditions were equally difficult; NR - not reported; NA - not applicable because no behavioural data was collected.

**Table S4.** List of studies included in the **moral reasoning** (N = 69) meta-analysis.

| Authors | Imaging method | No. of peaks | N | Task | Contrast | Difficulty RT | Difficulty Accuracy | Instructional cue |
|---|---|---|---|---|---|---|---|---|
| Akitsuki & Decety, 2009 | fMRI | 1 | 26 | Passive observation | intentional harm > accidental/no harm | | | Implicit |
| Avram et al., 2014 | fMRI | 10 | 16 | Right/wrong moral/factual judgements | moral (first person perspective) > non-moral | NR | NR | Explicit |
| | | | | Right/wrong moral/factual judgements | moral (third person perspective) > non-moral | | | Explicit |
| Bahnemann et al., 2010 | fMRI | 11 | 25 | Judge whether the protagonist is violating a moral/non-moral norm | moral > non-moral norm | E=C | E=C | Explicit |
| Bellucci et al., 2017 | fMRI | 6 | 26 | Determine the level of punishment deserved by the protagonist/ Estimate the number of syllables | moral > syllable judgement | NR | NA | Explicit |
| Chakroff et al., 2016 | fMRI | 20 | 20 | Rate how morally wrong the described act is | moral violations > non-moral (conjunction harmful & impure moral violations) | NR | NA | Explicit |
| Chen et al., 2020 | fMRI | 20 | 57 | Imagine being the protagonist/ Passive observation | harm > neutral scene | | | Implicit |
| | | | | Imagine being the protagonist/ Passive observation | help > neutral scene | | | Implicit |
| Chiong et al., 2013 | fMRI | 3 | 16 | Judge whether you would perform a hypothetical action in response to moral/practical dilemmas | moral > non-moral dilemmas | NR | NA | Explicit |
| de Achaval et al., 2013 | fMRI | 6 | 13 | Judge whether you would perform a hypothetical action in response to moral/practical dilemmas | moral > non-moral dilemmas | E=C | NA | Explicit |
| Dominguez et al., 2018 | fMRI | 2 | 48 | Imagine how you would feel about the decision to shoot an armed/unarmed target | unjustified (guilt-inducing) > justified killing | NR | NA | Explicit |
| Eslinger et al., 2009 | fMRI | 9 | 9 | Right/wrong moral/factual judgements | moral > non-moral | NR | NA | Explicit |
| FeldmanHall et al., 2012 | fMRI | 8 | 14 | Decide how much pain to administer to another person/ Decide which finger the other should move | moral > non-moral judgement (real & hypothetical scenario conjunction) | NR | NA | Explicit |

| | | | | | | | | |
|---|---|---|---|---|---|---|---|---|
| FeldmanHall et al., 2014 | fMRI | 10 | 38 | Judge whether you would perform a hypothetical action in response to moral/practical dilemmas | difficult moral > difficult non-moral dilemmas | E>C | NA | Explicit |
| | | | | Judge whether you would perform a hypothetical action in response to moral/practical dilemmas | easy moral > easy non-moral dilemmas | | | Explicit |
| Finger et al., 2006 | fMRI | 5 | 16 | Imagine you are the protagonist | moral > social/no violations | | | Implicit |
| Fourie et al., 2014 | fMRI | 8 | 22 | Implicit Associations Task with feedback | feedback after prejudice > neutral conditions | | | Implicit |
| Han et al., 2014 | fMRI | 28 | 8 | Evaluate whether the solution presented for the moral/practical dilemma is appropriate | moral (personal) > non-moral dilemmas | NR | NA | Explicit |
| | | | | Evaluate whether the solution presented for the moral/practical dilemma is appropriate | moral (impersonal) > non-moral dilemmas | NR | NA | Explicit |
| Han et al., 2014 | fMRI | 27 | 8 | Evaluate whether the solution presented for the moral/practical dilemma is appropriate | moral (personal) > non-moral dilemmas | NR | NA | Explicit |
| | | | | Evaluate whether the solution presented for the moral/practical dilemma is appropriate | moral (impersonal) > non-moral dilemmas | NR | NA | Explicit |
| Han et al., 2016 | fMRI | 25 | 16 | Judge whether an action is appropriate in response to moral/practical dilemmas | moral (personal) > non-moral judgements | NR | NA | Explicit |
| | | | | Judge whether an action is appropriate in response to moral/practical dilemmas | moral (impersonal) > non-moral judgements | NR | NA | Explicit |
| Harada et al., 2009 | fMRI | 9 | 18 | Judge whether the protagonist's behaviour was morally good or bad / Indicate the protagonist's gender | moral judgement > gender judgement | E=C | E=C | Explicit |
| Harenski & Hamann, 2006 | fMRI | 33 | 10 | Passive observation | moral violations > non-moral situations | | | Implicit |
| | | | | Passive observation | moral violations > odd/even number judgement | | | Implicit |
| | | | | Decrease your own emotional response to the depicted situations | moral violations > non-moral situations | | | Implicit |

| | | | | | | | | |
|---|---|---|---|---|---|---|---|---|
| | | | | Decrease your own emotional response to the depicted situation | moral violations > odd/even number judgement | | | Implicit |
| Harenski et al., 2008 | | 10 | 28 | Judge whether the pictures depict a moral violation & Rate the severity of the moral violation | moral violation > non-moral situation | NR | NA | Explicit |
| Harenski et al., 2010 | fMRI | 5 | 14 | Perceptual judgement - indoors/outdoors scene judgement | moral > non-moral situation | | | Implicit |
| Harenski et al., 2012 | fMRI | 12 | 36 | Rate the severity of the moral violation | moral violation > unpleasant non-moral situation | NR | NA | Explicit |
| | | | | Rate the severity of the moral violation | moral violation > neutral non-moral situation | NR | NA | Explicit |
| Harenski et al., 2012 | fMRI | 7 | 15 | Rate the severity of the moral violation | moral violation > unpleasant non-moral situation | NR | NA | Explicit |
| | | | | Rate the severity of the moral violation | moral violation > neutral non-moral situation | NR | NA | Explicit |
| Harrison et al., 2012 | fMRI | 11 | 73 | Judge whether you would perform a hypothetical action in response to moral/practical dilemmas | moral > non-moral dilemmas | NR | NA | Explicit |
| Heekeren et al., 2003 | fMRI | 9 | 8 | Right/wrong moral judgements/ Semantic plausibility judgement | moral > semantic judgements | C>E | NA | Explicit |
| Heekeren et al., 2005 | fMRI | 8 | 12 | Right/wrong moral/semantic judgements | moral > semantic judgements | C>E | NA | Explicit |
| Li et al., 2016 | fMRI | 5 | 24 | Rate the degree of blame deserved by the protagonist | immoral > neutral intentions | E>C | NA | Explicit |
| | | | | Rate the degree of blame deserved by the protagonist | immoral > neutral behaviours | E=C | NA | Explicit |
| Lim et al., 2017 | fMRI | 30 | 19 | Rate how acceptable specific actions are in response to moral/practical dilemmas | moral > non-moral dilemmas | C>E | NA | Explicit |
| Michl et al., 2014 | fMRI | 29 | 14 | Imagine the scenario described | shame > neutral scenarios | | | Implicit |
| | | | | Imagine the scenario described | guilt > neutral scenarios | | | Implicit |
| Molenberghs et al., 2014 | fMRI | 7 | 48 | Deliver shocks/monetary rewards to other | reward > neutral | E=C | E=C | Explicit |
| | | | | Deliver shocks/monetary rewards to other | shock > neutral | E=C | E=C | Explicit |
| Molenberghs et al., 2015 | fMRI | 12 | 48 | Imagine being the protagonist that shoots others and Indicate who was shot | shooting civilians > no shooting | E=C | E=C | Explicit |

| | | | | | | | | |
|---|---|---|---|---|---|---|---|---|
| | | | | Imagine being the protagonist that shoots others and Indicate who was shot | shooting soldiers > no shooting | E=C | E=C | Explicit |
| Molenberghs et al., 2016 | fMRI | 17 | 48 | Passive observation | intentional harm > neutral interactions | | | Implicit |
| Moll et al., 2001 | fMRI | 10 | 10 | Right/wrong moral/factual judgements | moral > factual judgements | NR | NA | Explicit |
| Moll et al., 2002a | fMRI | 3 | 7 | Right/wrong moral judgements | moral > non-moral scenarios | E=C | NA | Explicit |
| Moll et al., 2002b | fMRI | 17 | 7 | Passive observation | moral > unpleasant non-moral situations | | | Implicit |
| | | | | Passive observation | moral > neutral non-moral situations | | | Implicit |
| Moll et al., 2005 | fMRI | 22 | 13 | Indicate when you finish to read the statements | moral > neutral non-moral scenarios | | | Implicit |
| | | | | Indicate when you finish to read the statements | moral > disgusting non-moral scenarios | | | Implicit |
| Moll et al., 2007 | fMRI | 6 | 12 | Imagine you are the protagonist | guilt > neutral scenarios | | | Implicit |
| Oaten et al., 2018 | fMRI | 34 | 22 | Indicate when a scrambled stimulus appeared on the screen | moral (disgust-inducing) > non-moral scenarios | | | Implicit |
| | | | | Indicate when scrambles stimulus appeared | moral (anger-inducing) > non-moral scenarios | | | Implicit |
| Parkinson et al., 2011 | fMRI | 28 | 38 | Right/wrong moral judgements | harmful moral violation > non-moral | NR | NA | Explicit |
| | | | | Right/wrong moral judgements | dishonest moral violations > non-moral | NR | NA | Explicit |
| | | | | Right/wrong moral judgements | disgusting moral violations > non-moral | NR | NA | Explicit |
| Pujol et al., 2012 | fMRI | 10 | 22 | Judge whether you would perform a hypothetical action in response to moral dilemmas/ Factual judgements for non-moral scenarios | moral > factual judgements | NR | NR | Explicit |
| Reniers et al., 2012 | fMRI | 32 | 24 | Right/wrong moral/practical judgements | moral > practical scenarios | E=C | NA | Explicit |
| | | | | Right/wrong moral/practical judgements | moral judgements > rest | NA | NA | Explicit |
| Robertson et al., 2007 | fMRI | 11 | 16 | Read the scenarios and identify important aspects | moral > neutral non-moral scenarios | | | Implicit |
| | | | | Read the scenarios and identify important aspects | moral > strategical/tactical non-moral scenarios | | | Implicit |
| Sakai et al., 2017 | fMRI | 26 | 24 | Decide whether to donate money to charity or keep the money | donation decision > mathematical calculation | NR | NA | Explicit |

| | | | | | | | | |
|---|---|---|---|---|---|---|---|---|
| Schaich Borg et al., 2006 | fMRI | 7 | 24 | Right/wrong moral/practical judgements and Judge whether you would perform a hypothetical action in response to the scenarios | moral > non-moral scenarios | E=C & C>E | NA | Explicit |
| Schaich Borg et al., 2008 | fMRI | 17 | 50 | Memory recognition task | moral violations > disgusting non-moral scenarios | | | Implicit |
| Schleim et al., 2011 | fMRI | 6 | 40 | Right/wrong moral/practical judgements | moral > non-moral scenarios | E>C | NA | Explicit |
| Schneider et al., 2013 | fMRI | 4 | 28 | Judge whether you would perform a hypothetical action in response to moral/ practical dilemmas | moral > non-moral dilemmas | E>C | NA | Explicit |
| Seara-Cardoso et al., 2016 | fMRI | 10 | 28 | Imagine yourself in the scenario and rate the intensity of guilt you would feel | moral violations > non-moral scenarios | | | Implicit |
| Seara-Cardoso et al., 2016 | fMRI | 5 | 28 | Rate how wrong the described behaviour is | moral violations > non-moral | NR | NA | Explicit |
| Sevinc et al., 2017 | fMRI | 24 | 20 | Sentence completion | moral > non-moral scenarios | | | Implicit |
| Shin et al., 2000 | PET | 8 | 8 | Imagine the scenario | guilt > neutral scenario | | | Implicit |
| Smith et al., 2015 | fMRI | 7 | 30 | Judge the fairness of resource allocation/ the correctedness of the diagram representation | moral > non-moral judgement | NR | NA | Explicit |
| Sommer et al., 2010 | fMRI | 6 | 12 | Choose a hypothetical action in response to moral/non-moral dilemmas | moral > non-moral dilemmas | E=C | NA | Explicit |
| Sommer et al., 2014 | fMRI | 6 | 16 | Choose a hypothetical action in response to moral/social non-moral dilemmas | moral > non-moral dilemmas | E=C | NA | Explicit |
| Sommer et al., 2014 | fMRI | 10 | 16 | Choose a hypothetical action in response to moral/social non-moral dilemmas | moral > non-moral dilemmas | E=C | NA | Explicit |
| Takahashi et al., 2004 | fMRI | 5 | 19 | Rate the intensity of guilt/embarrassment of the situations | moral guilt > neutral scenarios | | | Implicit |
| Takahashi et al., 2008 | fMRI | 4 | 15 | Imagine the scenario described and rate how moral/immoral and praiseworthy/blameworthy they are | moral beauty > non-moral scenarios | NR | NA | Explicit |

| | | | | | | | | |
|---|---|---|---|---|---|---|---|---|
| Teed et al., 2020 | fMRI | 2 | 20 | Rate your willingness to participate in the action described | self-transcendence > self-enhancement | NR | NA | Explicit |
| Theriault et al., 2017 | fMRI | 55 | 25 | Rate the degree of agreement with the fact/preference/moral statement | moral > non-moral factual statements | NR | NA | Explicit |
| | | | | Rate the degree of agreement with the fact/preference/moral statement | moral > non-moral preference statements | NR | NA | Explicit |
| Thomas et al., 2019 | fMRI | 5 | 41 | Judge whether you would perform a hypothetical action in response to moral/practical dilemmas | moral > non-moral dilemmas | NR | NA | Explicit |
| Tsoi et al., 2018 | fMRI | 25 | 25 | Judge how morally wrong the behaviour described is | moral violations (psychological harm) > non-moral scenarios | E>C | NA | Explicit |
| | | | | Judge how morally wrong the behaviour described is | moral violations (physical harm) > non-moral scenarios | E>C | NA | Explicit |
| Ty et al., 2017 | fMRI | 18 | 18 | Decide whether to donate money to charity | harm induced by participant decision > computer decision | NA | NA | Explicit |
| Verdejo-Garcia et al., 2014 | fMRI | 8 | 14 | Judge whether you would perform a hypothetical action in response to moral dilemmas/ Indicate the outcome of a non-moral scenario | moral > non-moral scenarios | NR | NA | Explicit |
| Wagner et al., 2011 | fMRI | 24 | 15 | Remember and mentally relive past experiences & Rate emotional experience | guilt > neutral memories | | | Implicit |
| Wang et al., 2015 | fMRI | 10 | 28 | Judge whether the cartoon depicted a moral or neutral behaviour/ Gender judgement | moral > gender judgement | E=C | E=C | Explicit |
| Wang et al., 2015 | fMRI | 11 | 22 | Gender judgement | moral beauty > non-moral scene | | | Implicit |
| White et al., 2017 | fMRI | 16 | 23 | Rate how wrong the depicted action is | moral > social norm violations | C>E | NA | Explicit |
| Yang et al., 2018 | fMRI | 13* | 32 | Recall the scenario associated with the videos and rate the intensity of emotions elicited | moral virtue > skill scenarios | | | Implicit |
| Yu et al., 2014 | fMRI | 1 | 24 | Perceptual (size) judgement in an interactive game; feedback in the form of painful stimulation to both interaction partners | participant incorrect (guilt-inducing) > both incorrect | | | Implicit |

*the coordinates were obtained via personal communication with the authors.

N = sample size; E>C - the experimental condition was more difficult than the control condition; C>E - the control condition was more difficult than the experimental condition; E=C - the experimental and control conditions were equally difficult; NR - not reported; NA - not applicable because no behavioural data was collected.

**Table S5.** List of studies included in the **semantic control** (N = 96) meta-analysis.

| Authors | Imaging method | No. of peaks | N | Task | Contrast |
|---|---|---|---|---|---|
| Ahrens et al., 2007 | fMRI | 32 | 8 | Sentence comprehension<br>Sentence comprehension | figurative (conventional metaphors) > literal sentences<br>figurative (anomalous metaphors) > literal sentences |
| Allen et al., 2008 | fMRI | 5 | 15 | Hayling sentence completion task | Non-salient > salient completion |
| Amunts et al., 2004 | fMRI | 7 | 11 | Verbal fluency | category > rote (days of the week, months, alphabet letters) |
| Assaf et al., 2006 | fMRI | 9 | 18 | Object recall from feature pairs | target object present > absent |
| Bambini et al., 2011 | fMRI | 10 | 9 | Semantic relatedness judgement | figurative > literal sentences |
| Bedny et al., 2008 | fMRI | 11 | 20 | Semantic relatedness judgement<br>Semantic relatedness judgement | inconsistent > consistent > control<br>ambiguous > unambiguous words |
| Benedek et al., 2014 | fMRI | 8 | 28 | Generate metaphoric/literal synonyms for the words depicted | figurative > literal meaning |
| Benedek et al., 2018 | fMRI | 12 | 42 | Alternate object uses generation<br>Alternate object uses generation | Non-salient (create) > salient attributes<br>Non-salient (recall) > salient attributes |
| Bitan et al., 2017 | fMRI | 4 | 23 | Semantic relatedness judgement (homonyms) | subordinate > dominant meaning |
| Bottini et al., 1994 | PET | 6 | 6 | Sentence plausibility judgement | figurative > literal meaning |
| Boulenger, et al., 2009 | fMRI | 6 | 18 | Sentence comprehension<br>Sentence comprehension | figurative > literal meaning (timepoint 1)<br>figurative > literal meaning (timepoint 2) |
| Chan et al., 2004 | fMRI | 17 | 6 | Generate a word that is associated with the target word | ambiguous > unambiguous words |
| Chen et al., 2008 | fMRI | 5 | 14 | Sentence plausibility judgement | figurative > literal meaning |
| Chiou et al., 2018 | fMRI | 6* | 18 | Semantic relatedness judgement | feature matching > global association |
| Citron & Goldberg, 2014 | fMRI | 19 | 26 | Sentence comprehension | figurative > literal sentences |
| Citron et al., 2016 | fMRI | 69 | 24 | Sentence comprehension | figurative > literal sentences |
| Citron et al., 2019 | fMRI | 15 | 23 | Sentence comprehension | figurative > literal sentences |
| Collette et al., 2001 | PET | 4 | 12 | Hayling sentence completion task | non-salient > salient completion |
| Davey et al., 2015 | fMRI | 30 | 17 | Semantic relatedness judgement | feature matching > global association |
| Davey et al., 2016 | fMRI | 29 | 20 | Semantic relatedness judgement | feature matching > global associations |
| de Zubicaray et al., 2000 | fMRI | 13 | 6 | Hayling task variant - provide the superordinate category | initiate > suppress the correct response |
| de Zubicaray et al., 2001 | fMRI | 4 | 8 | Picture-word interference task | semantically-related distractors > lexical control |
| de Zubicaray et al., 2006 | fMRI | 3 | 13 | Competitor priming in a picture naming task | competitor primed > not primed |
| de Zubicaray et al., 2017 | fMRI | 2 | 21 | Verb generation | switching > same category |
| Desai et al., 2011 | fMRI | 8 | 22 | Sentence plausibility judgement | figurative > literal sentences |
| Desai et al., 2013 | fMRI | 7 | 27 | Semantic plausibility judgement | figurative (metaphors) > literal abstract sentences |

| | | | | | |
|---|---|---|---|---|---|
| | | | | Semantic plausibility judgement | figurative (idioms) > literal abstract sentences |
| Diaz & Hogstrom, 2011 | fMRI | 6 | 16 | Semantic relatedness judgement | figurative > literal sentences |
| Fink et al., 2010 | fMRI | 1 | 31 | Alternate object uses generation | Non-salient > salient attributes |
| Forgacs et al., 2012 | fMRI | 8 | 40 | Familiarity judgement | figurative > literal expressions |
| Gurd et al., 2002 | fMRI | 10 | 11 | Verbal fluency | category > rote (days of the week, months, alphabet letters) |
| | | | | Verbal fluency | switching categories > free generation |
| Hallam et al., 2016 | fMRI | 8 | 18 | Semantic relatedness judgement | weak > strong target-probe association |
| Hargreaves et al., 2011 | fMRI | 2 | 20 | Semantic classification | ambiguous > unambiguous words |
| Hillert & Buracas, 2009 | fMRI | 5 | 10 | Sentence plausibility judgement | figurative > literal sentences |
| Hirshorn & Thompson-Schill, 2006 | fMRI | 15 | 10 | Verbal fluency | switching categories > free generation |
| Hirshorn & Thompson-Schill, 2006 | fMRI | 21 | 9 | Verbal fluency | Switching > clustering |
| Hoenig & Scheef, 2009 | fMRI | 15 | 22 | Semantic relatedness judgement | ambiguous > unambiguous sentences |
| Humphreys & Lambon Ralph, 2017 | fMRI | 10* | 20 | Semantic relatedness judgement | ambiguous > unambiguous words |
| Joue et al., 2018 | fMRI | 6 | 32 | Judge whether the sentences & gestures conveyed a figurative/literal meaning | figurative (external metonymic) > literal sentences |
| | | | | Judge whether the sentences & gestures conveyed a figurative/literal meaning | figurative (internal metonymic) > literal sentences |
| Kennedy et al., 2015 | fMRI | 17* | 271 | Semantic classification | hard > easy decisions |
| Kircher et al., 2007 | fMRI | 1 | 12 | Valence judgement | figurative > literal sentences |
| Klepousniotou et al., 2014 | fMRI | 3 | 15 | Semantic relatedness judgement (homonyms) | subordinate > dominant meaning |
| | | | | Semantic relatedness judgement (metaphors/idioms) | figurative > literal words |
| Krieger-Redwood et al., 2015 | fMRI | 53 | 22 | Semantic relatedness judgement | weak > strong probe-target association |
| | | | | Semantic relatedness judgement | weak > strong probe-target association |
| Lacey et al., 2012 | fMRI | 4 | 7 | Sentence comprehension | figurative > literal sentences |
| Lauro et al., 2007 | fMRI | 10 | 22 | Judge whether the cartoon matched the meaning of the sentence | figurative > literal sentences |
| Lauro et al., 2013 | fMRI | 17 | 24 | Semantic congruency judgement | figurative > literal sentences |
| Lee & Dapretto, 2006 | fMRI | 3 | 12 | Semantic relatedness judgement | figurative > literal meaning |
| Lopes et al., 2016 | fMRI | 6 | 24 | Attribute judgement | difficult > easy decision |
| Mashal et al., 2007 | fMRI | 8 | 15 | Semantic decision | figurative > literal expressions |
| | | | | Judge the type of semantic relatedness (metaphorical/literal/unrelated) | figurative > literal expressions |

| | | | | | |
|---|---|---|---|---|---|
| Mashal et al., 2009 | fMRI | 5 | 15 | Valence judgement<br>Valence judgement | figurative > nonsensical sentences<br>figurative > literal meaning |
| Mashal et al., 2013 | fMRI | 2 | 14 | Semantic plausibility judgement | figurative > literal expressions |
| Mason & Just, 2007 | fMRI | 9 | 12 | Sentence comprehension<br>Sentence comprehension | ambiguous > unambiguous sentences<br>subordinate > dominant meaning |
| Mestres-Misse et al. 2016 | fMRI | 5 | 23 | Semantic plausibility judgement | ambiguous > unambiguous sentences |
| Methqal et al., 2017 | fMRI | 8 | 20 | Semantic relatedness judgement | switching > maintaining rule |
| Methqal et al., 2017 | fMRI | 7 | 20 | Semantic relatedness judgement | switching > maintaining rule |
| Methqal et al., 2017 | fMRI | 25 | 40 | Semantic relatedness judgement | weak > strong target-probe association |
| Methqal et al., 2019 | fMRI | 18 | 13 | Verbal fluency | category > rote (months) |
| Methqal et al., 2019 | fMRI | 9 | 13 | Verbal fluency | category > rote (months) |
| Musz & Thompson-Schill, 2017 | fMRI | 2 | 13 | Sentence comprehension<br>Sentence comprehension | delayed > prior disambiguating context<br>ambiguous > unambiguous sentences |
| Nelson et al., 2009 | fMRI | 5 | 17 | Verb generation | high > low selection demands |
| Noppeny & Price, 2002 | PET | 7 | 12 | Attribute judgement | semantic decision > repetition |
| Noppeny et al., 2004 | fMRI | 2 | 15 | Semantic relatedness judgement | target word repeated in different > same triads |
| Obert et al., 2014 | fMRI | 3 | 19 | Judge whether the sentences conveyed a figurative/literal meaning | figurative > literal sentences |
| Ochsner et al., 2009 | fMRI | 16 | 16 | Eriksen flanker task variant | incongruent > congruent |
| Paschke et al., 2015 | fMRI | 12 | 115 | Eriksen flanker task variant | incongruent > congruent |
| Prat et al., 2012 | MRI | 12 | 24 | Sentence comprehension | difficult > easy comprehension |
| Ragland et al., 2008 | fMRI | 6 | 14 | Semantic fluency<br>Semantic fluency | category > rote<br>switching > same categories |
| Raposo et al., 2012 | fMRI | 1 | 17 | Factual judgement | less > more shared features |
| Rapp et al., 2004 | fMRI | 3 | 15 | Valence judgement | figurative > literal meaning |
| Rapp et al., 2011 | fMRI | 8 | 14 | Semantic plausibility judgement | figurative > literal sentences |
| Raucher-Chene et al., 2018 | fMRI | 3 | 35 | Semantic relatedness judgement | ambiguous > unambiguous words |
| Rizio et al., 2017 | fMRI | 11 | 40 | Picture-word interference task | semantically-related distractors > lexical control |
| Rodd et al., 2005 | fMRI | 7 | 30 | Semantic relatedness judgement/Passive listening | ambiguous > unambiguous sentences |
| Rodd et al., 2010 | fMRI | 1 | 14 | Sentence comprehension | ambiguous > unambiguous sentences |
| Samur et al., 2015 | fMRI | 3 | 20 | Sentence comprehension | figurative > literal sentences |

| | | | | | |
|---|---|---|---|---|---|
| Satpute et al., 2014 | fMRI | 6 | 33 | Semantic relatedness judgement | weak > strong probe-target association & high > low selection demands |
| Schmidt & Seger, 2009 | fMRI | 8 | 10 | Sentence comprehension<br>Sentence comprehension | figurative > literal sentences<br>difficult > easy comprehension |
| Seger et al., 2000 | fMRI | 14 | 7 | Verb generation | Non-salient > salient |
| Sharp et al., 2010 | PET | 4 | 12 | Semantic relatedness judgement | weak > strong probe-target association |
| Shibata et al., 2007 | fMRI | 5 | 13 | Sentence comprehension | figurative > literal meaning |
| Shibata et al., 2012 | fMRI | 6 | 24 | Sentence comprehension | figurative > literal sentences |
| Smirnov et al., 2014 | fMRI | 1 | 20 | Sentence comprehension | mismatching > matching contextual cues |
| Snyder et al., 2007 | fMRI | 13 | 14 | Semantic relatedness judgement | feature matching > global association |
| Snyder et al., 2011 | fMRI | 47 | 17 | Verb generation<br>Verb generation | strong > weak selection demands<br>weak > strong cue-response association |
| Stringaris et al., 2006 | fMRI | 8 | 12 | Semantic relatedness judgement<br>Semantic relatedness judgement | non-relevant words following metaphors > literal sentences<br>relevant words following metaphors > literal sentences |
| Stringaris et al., 2007 | fMRI | 9 | 11 | Sentence plausibility judgement | figurative > literal meaning |
| Thompson-Schill et al., 1997 | fMRI | 16 | 5 | Verb generation<br>Semantic classification<br>Semantic relatedness judgement | high > low selection demands<br>high > low selection demands<br>feature matching > global association |
| Uchiyama et al., 2012 | fMRI | 11 | 20 | Judge whether the sentences conveyed a figurative/sarcastic/literal meaning | figurative > literal sentences |
| Vitello et al., 2014 | fMRI | 3 | 17 | Semantic relatedness judgement | ambiguous > unambiguous sentences |
| Wagner et al., 2001 | fMRI | 21 | 14 | Semantic relatedness judgement<br>Semantic relatedness judgement | Four-word > two-word choices<br>weak > strong probe-target association |
| Whitney et al., 2009 | fMRI | 7 | 15 | Semantic relatedness judgement<br>Semantic relatedness judgement<br>Semantic relatedness judgement | ambiguous > unambiguous words (double-related)<br>ambiguous > unambiguous words (single-related)<br>subordinate > dominant meaning |
| Whitney et al., 2011 | fMRI | 8 | 15 | Semantic relatedness judgement | subordinate > dominant meaning |
| Yang et al., 2009 | fMRI | 7 | 18 | Valence judgement | figurative > literal sentences |
| Yang et al., 2010 | fMRI | 13* | 13 | Valence judgement | figurative > literal sentences |
| Yi et al., 2017 | fMRI | 3 | 15 | Sentence comprehension | figurative > literal sentences |
| Zempleni et al., 2007 | fMRI | 8 | 15 | Sentence comprehension | figurative > literal meaning |
| Zhang et al., 2004 | fMRI | 3 | 14 | Semantic relatedness judgement for reversible words | semantically-related > unrelated distractors |

*the coordinates were obtained via personal communication with the authors.
N = sample size.
