## Supplementary material for "Establishing a Role of the Semantic Control Network in Social Cognitive Processing: A Meta-analysis of Functional Neuroimaging Studies": SI3. Results tables

for

**A Meta-analysis of Functional Neuroimaging Studies**

Veronica Diveica, Kami Koldewyn & Richard J. Binney

This document contains the results tables for all analyses conducted. The input files and outputs of all analyses can be accessed via OSF (osf.io/fktb8/).

**Table S1.1.1.** Significant ALE clusters (cluster extent-based FWE threshold, p < .05) of convergent activation across 136 **ToM** experiments. Anatomical labels are derived from the Automatic Anatomical Labelling Atlas.

| Cluster | Volume (mm3) | ALE value | Z value | Peak MNI coordinates | | | Hemisphere | Anatomical region |
|---|---|---|---|---|---|---|---|---|
| | | | | X | Y | Z | | |
| 1 | 26232 | 0.17* | 14.11 | -50 | -58 | 24 | L | Middle temporal gyrus |
| | | 0.10* | 9.46 | -58 | -10 | -14 | L | Middle temporal gyrus |
| | | 0.09* | 8.45 | -52 | 2 | -26 | L | Middle temporal gyrus |
| | | 0.08* | 7.84 | -60 | -22 | -8 | L | Middle temporal gyrus |
| | | 0.07* | 7.15 | -56 | -38 | 0 | L | Middle temporal gyrus |
| 2 | 24304 | 0.13* | 11.37 | 54 | -52 | 22 | R | Middle temporal gyrus |
| | | 0.10* | 9.20 | 54 | 0 | -22 | R | Middle temporal gyrus |
| | | 0.10* | 8.98 | 60 | -8 | -16 | R | Middle temporal gyrus |
| | | 0.09* | 8.37 | 50 | 8 | -28 | R | Middle temporal gyrus |
| | | 0.06* | 5.89 | 50 | -30 | -4 | R | Middle temporal gyrus |
| | | 0.04 | 3.89 | 50 | -72 | 6 | R | Middle temporal gyrus |
| 3 | 12040 | 0.13* | 11.04 | -10 | 54 | 34 | L | Medial superior frontal gyrus |
| | | 0.10* | 9.05 | 4 | 56 | 22 | R | Medial superior frontal gyrus |
| | | 0.08* | 7.86 | -4 | 56 | 24 | L | Medial superior frontal gyrus |
| 4 | 10512 | 0.14* | 12.23 | -2 | -54 | 36 | L | Precuneus |
| 5 | 8656 | 0.10* | 8.90 | -48 | 28 | -10 | L | Inferior frontal gyrus (pars orbitalis) |
| | | 0.09* | 8.32 | -52 | 24 | 6 | L | Inferior frontal gyrus (pars triangularis) |
| | | 0.07* | 7.16 | -52 | 18 | 18 | L | Inferior frontal gyrus (pars opercularis) |
| | | 0.04 | 4.21 | -40 | 20 | -24 | L | Superior temporal pole |
| 6 | 6544 | 0.10* | 9.31 | 56 | 28 | 6 | R | Inferior frontal gyrus (pars triangularis) |
| | | 0.05* | 5.26 | 44 | 20 | 24 | R | Inferior frontal gyrus (pars triangularis) |
| 7 | 4096 | 0.07* | 7.13 | 0 | 48 | -18 | L/R | Gyrus rectus |
| | | 0.04 | 4.76 | -8 | 52 | 0 | L | Anterior cingulate gyrus |
| 8 | 3552 | 0.07* | 7.30 | -6 | 14 | 62 | L | Supplementary motor area |
| 9 | 2320 | 0.08* | 8.02 | -24 | -78 | -36 | L | Cerebellum (crus 2) |
| 10 | 1976 | 0.06* | 6.11 | 42 | 6 | 42 | R | Precentral gyrus |
| 11 | 1968 | 0.05* | 5.62 | -42 | 0 | 54 | L | Precentral gyrus |

| | | | | | | | | |
|---|---|---|---|---|---|---|---|---|
| 12 | 1472 | 0.06* | 6.40 | -42 | -50 | -16 | L | Fusiform gyrus |
| 13 | 1408 | 0.07* | 7.22 | 28 | -78 | -34 | R | Cerebellum (crus 1) |

MNI = Montreal Neurological Institute (MNI) standard stereotaxic space
* Indicates the peaks/sub-peaks withstanding the voxel-level FWE p < .05 threshold. This analysis revealed an additional cluster in the middle cingulate gyrus ( [0, -16, 38], ALE value 0.06, Z value 6.04, cluster size 152 $mm^{3)}$ which did not withstand the cluster extent-based threshold.

**Table S1.1.2.** Significant ALE clusters (cluster extent-based FWE threshold, p < .05) of convergent activation across 43 **false belief reasoning** experiments. Anatomical labels are derived from the Automatic Anatomical Labelling Atlas.

| **Cluster** | **Volume (mm3)** | **ALE value** | **Z value** | **Peak MNI coordinates** | | | **Hemisphere** | **Anatomical region** |
|---|---|---|---|---|---|---|---|---|
| | | | | **X** | **Y** | **Z** | | |
| 1 | 7536 | 0.04* | 6.43 | 52 | 6 | -32 | R | Inferior temporal gyrus |
| | | 0.04* | 6.29 | 56 | -18 | -14 | R | Middle temporal gyrus |
| | | 0.04* | 5.90 | 56 | 2 | -22 | R | Middle temporal gyrus |
| 2 | 7264 | 0.07* | 9.49 | 2 | -56 | 34 | R | Precuneus |
| 3 | 6744 | 0.05* | 7.08 | -62 | -24 | -8 | L | Middle temporal gyrus |
| | | 0.04* | 6.78 | -54 | 2 | -24 | L | Middle temporal gyrus |
| | | 0.03* | 5.37 | -58 | -10 | -14 | L | Middle temporal gyrus |
| | | 0.02 | 3.87 | -60 | -34 | -4 | L | Middle temporal gyrus |
| 4 | 6192 | 0.05* | 7.13 | 2 | 56 | 18 | R | Medial superior frontal gyrus |
| | | 0.05* | 7.08 | 2 | 54 | 22 | R | Medial superior frontal gyrus |
| | | 0.03* | 5.13 | -10 | 54 | 32 | L | Medial superior frontal gyrus |
| 5 | 6032 | 0.10* | 12.05 | -52 | -58 | 24 | L | Middle temporal gyrus |
| 6 | 5632 | 0.08* | 9.93 | 54 | -54 | 24 | R | Superior temporal gyrus |
| 7 | 1632 | 0.04* | 6.11 | 2 | 50 | -16 | R | Gyrus rectus |
| 8 | 1312 | 0.03* | 5.47 | -26 | 80 | -36 | R | Cerebellum |
| 9 | 1120 | 0.03* | 5.65 | 24 | 26 | 44 | R | Middle frontal gyrus |

MNI = Montreal Neurological Institute (MNI) standard stereotaxic space
* Indicates the peaks/sub-peaks withstanding the voxel-level FWE p < .05 threshold.

**Table S1.2.** Significant ALE clusters (cluster extent-based FWE threshold, p< .05) of convergent activation across 40 **trait inference** experiments. Anatomical labels are derived from the Automatic Anatomical Labelling Atlas.

| Cluster | Volume (mm3) | ALE value | Z value | Peak MNI coordinates | | | Hemisphere | Anatomical region |
|---|---|---|---|---|---|---|---|---|
| | | | | X | Y | Z | | |
| 1 | 12928 | 0.04* | 6.78 | 0 | 56 | 22 | L/R | Medial superior frontal gyrus |
| | | 0.04* | 6.71 | -6 | 54 | 34 | L | Medial superior frontal gyrus |
| | | 0.03* | 5.87 | -8 | 34 | 56 | L | Medial superior frontal gyrus |
| | | 0.03* | 5.46 | -8 | 54 | 42 | L | Medial superior frontal gyrus |
| | | 0.02 | 3.90 | -8 | 20 | 56 | L | Supplementary motor area |
| | | 0.02 | 3.78 | 4 | 48 | 46 | R | Medial superior frontal gyrus |
| | | 0.02 | 3.52 | -8 | 32 | 44 | L | Medial superior frontal gyrus |
| | | 0.02 | 3.29 | 0 | 18 | 50 | L/R | Supplementary motor area |
| 2 | 3856 | 0.03* | 5.84 | -40 | 22 | -16 | L | Inferior frontal gyrus (pars orbitalis) |
| | | 0.03* | 5.81 | -44 | 22 | -14 | L | Superior temporal pole |
| | | 0.02 | 4.45 | -30 | 18 | -24 | L | Inferior frontal gyrus (pars orbitalis) |
| | | 0.02 | 3.83 | -50 | 28 | 2 | L | Inferior frontal gyrus (pars triangularis) |
| 3 | 3088 | 0.04* | 6.74 | -2 | -52 | 28 | L | Posterior cingulate gyrus |
| 4 | 2064 | 0.03* | 4.94 | 48 | 28 | -12 | R | Inferior frontal gyrus (pars orbitalis) |
| | | 0.02 | 4.24 | 54 | 28 | 0 | R | Inferior frontal gyrus (pars triangularis) |
| 5 | 1368 | 0.03* | 4.93 | -48 | -62 | 26 | L | Angular gyrus |
| 6 | 1088 | 0.02 | 4.12 | 56 | -64 | 24 | R | Middle temporal gyrus |
| | | 0.02 | 4.12 | 48 | -62 | 30 | R | Angular gyrus |
| | | 0.02 | 3.66 | 48 | -52 | 30 | R | Angular gyrus |
| 7 | 1072 | 0.03* | 4.98 | -62 | -16 | -18 | L | Middle temporal gyrus |
| 8 | 1056 | 0.02 | 4.59 | 4 | 56 | -6 | R | Medial orbitofrontal cortex |
| | | 0.02 | 3.47 | -4 | 48 | -18 | L | Gyrus rectus |

MNI = Montreal Neurological Institute (MNI) standard stereotaxic space
* Indicates the peaks/sub-peaks withstanding the voxel-level FWE p < .05 threshold.

**Table S1.3.1.** Significant ALE clusters (cluster extent-based FWE threshold, p< .05) of convergent activation across 164 **empathy** experiments. Anatomical labels are derived from the Automatic Anatomical Labelling Atlas.

| | **Volume (mm3)** | **ALE value** | **Z value** | **Peak MNI coordinates** | | | **Hemisphere** | **Anatomical region** |
|---|---|---|---|---|---|---|---|---|
| | | | | **X** | **Y** | **Z** | | |
| 1 | 23216 | 0.11* | 9.42 | -32 | 22 | 4 | L | Insula |
| | | 0.08* | 7.30 | -40 | 14 | -4 | L | Insula |
| | | 0.07* | 6.73 | -52 | 10 | 10 | L | Inferior frontal gyrus (pars opercularis) |
| | | 0.07* | 6.68 | -52 | 10 | 16 | L | Inferior frontal gyrus (pars opercularis) |
| | | 0.07* | 6.22 | -40 | 0 | -4 | L | Insula |
| | | 0.07* | 6.19 | -54 | 8 | 30 | L | Precentral gyrus |
| | | 0.05 | 4.60 | -54 | 24 | 10 | L | Inferior frontal gyrus (pars triangularis) |
| | | 0.04 | 4.10 | -42 | 2 | 46 | L | Precentral gyrus |
| | | 0.04 | 3.82 | -42 | 42 | 8 | L | Inferior frontal gyrus (pars triangularis) |
| | | 0.04 | 3.47 | -36 | 50 | 14 | L | Middle frontal gyrus |
| | | 0.04 | 3.42 | -46 | 32 | 14 | L | Inferior frontal gyrus (pars triangularis) |
| 2 | 18136 | 0.08* | 7.43 | 50 | 30 | -2 | R | Inferior frontal gyrus (pars orbitalis) |
| | | 0.08* | 7.04 | 44 | 24 | -2 | R | Insula |
| | | 0.07* | 6.13 | 34 | 22 | 4 | R | Insula |
| | | 0.07* | 6.03 | 50 | 6 | 32 | R | Precentral gyrus |
| | | 0.06* | 5.41 | 52 | 22 | 20 | R | Inferior frontal gyrus (pars triangularis) |
| | | 0.06* | 5.24 | 42 | 4 | 0 | R | Insula |
| | | 0.05 | 4.94 | 30 | 22 | -16 | R | Insula |
| | | 0.05 | 4.91 | 60 | 12 | 14 | R | Inferior frontal gyrus (pars opercularis) |
| | | 0.05 | 4.29 | 58 | 14 | 22 | R | Inferior frontal gyrus (pars opercularis) |
| | | 0.04 | 3.72 | 46 | 38 | 10 | R | Inferior frontal gyrus (pars triangularis) |
| 3 | 15144 | 0.10* | 8.52 | -4 | 18 | 44 | L | Medial superior frontal gyrus |
| | | 0.06* | 5.60 | 6 | 10 | 62 | R | Supplementary motor area |
| | | 0.06* | 5.46 | 2 | 10 | 58 | R | Supplementary motor area |
| | | 0.05 | 4.85 | -4 | 26 | 26 | L | Anterior cingulate gyrus |
| | | 0.05 | 4.68 | 0 | 18 | 28 | L/R | Anterior cingulate gyrus |
| 4 | 6912 | 0.09* | 7.55 | 50 | -64 | -4 | R | Inferior temporal gyrus |
| | | 0.05 | 4.95 | 44 | -48 | -20 | R | Fusiform gyrus |

| | | | | | | | | |
|---|---|---|---|---|---|---|---|---|
| 5 | 6824 | 0.11* | 9.25 | -46 | -70 | -2 | L | Inferior occipital gyrus |
| | | 0.04 | 3.91 | -52 | -60 | 16 | L | Middle temporal gyrus |
| 6 | 5792 | 0.12* | 10.21 | -58 | -24 | 32 | L | Supramarginal gyrus |
| | | 0.04 | 3.34 | -60 | -40 | 26 | L | Supramarginal gyrus |
| 7 | 4736 | 0.11* | 9.57 | 62 | -22 | 34 | R | Supramarginal gyrus |
| | | 0.04 | 3.83 | 62 | -34 | 22 | R | Superior temporal gyrus |
| 8 | 4432 | 0.08* | 7.19 | -4 | -30 | -4 | L | Brainstem^ |
| | | 0.06* | 5.61 | -10 | -12 | 8 | L | Thalamus |
| | | 0.04 | 3.65 | -6 | -28 | -18 | L | Brainstem^ |
| | | 0.04 | 3.62 | -6 | -16 | -2 | L | Thalamus^ |
| 9 | 2856 | 0.06* | 5.32 | 36 | -86 | 6 | R | Middle occipital gyrus |
| | | 0.05* | 5.10 | 18 | -92 | -2 | R | Calcarine fissure |
| | | 0.05 | 4.43 | 32 | -88 | -4 | R | Inferior occipital gyrus |
| 10 | 2544 | 0.07* | 6.08 | -14 | -96 | -6 | L | Inferior occipital gyrus |
| | | 0.06* | 5.49 | -32 | -92 | -2 | L | Middle occipital gyrus |
| 11 | 1888 | 0.06* | 5.53 | -12 | 4 | 12 | L | Caudate |
| | | 0.05 | 4.34 | -12 | 12 | 2 | L | Caudate |
| | | 0.04 | 3.91 | -18 | 2 | 4 | L | Pallidum |
| | | 0.03 | 3.22 | -8 | -2 | 0 | L | Pallidum^ |
| 12 | 1880 | 0.05 | 4.57 | 14 | 6 | 8 | R | Pallidum^ |
| | | 0.05 | 4.46 | 16 | 4 | 2 | R | Pallidum |
| 13 | 1448 | 0.05 | 4.82 | 50 | -34 | 0 | R | Middle temporal gyrus |
| | | 0.04 | 4.20 | 54 | -24 | -8 | R | Middle temporal gyrus |
| | | 0.04 | 3.51 | 60 | -40 | 8 | R | Middle temporal gyrus |
| 14 | 1144 | 0.05 | 4.89 | -2 | 54 | 26 | L | Medial frontal gyrus |
| 15 | 1120 | 0.08* | 7.01 | -20 | -8 | -16 | L | Hippocampus |
| 16 | 1008 | 0.06* | 5.38 | -40 | -46 | 56 | L | Inferior parietal lobule |

MNI = Montreal Neurological Institute (MNI) standard stereotaxic space
* Indicates the peaks/sub-peaks withstanding the voxel-level FWE $p < .05$ threshold. This analysis revealed two additional clusters in the right cerebellum ( [34, -62, -26], ALE value 0.06, Z value 5.26, cluster size 24 $mm^3$) and the right hippocampus ( [22, -6, -14], ALE value 0.06, Z value 5.24, cluster size 24 $mm^3$) and which did not withstand the cluster extent-based threshold.

^ The anatomical region was labelled based on visual inspection of the ALE map because the coordinates fall outside the Automated Anatomical Labelling Atlas.

**Table S1.3.2.** Significant ALE clusters (cluster extent-based FWE threshold, p< .05) of convergent activation across 85 **empathy for pain** experiments. Anatomical labels are derived from the Automatic Anatomical Labelling Atlas.

| Cluster | Volume (mm3) | ALE value | Z value | Peak MNI coordinates | | | Hemisphere | Anatomical region |
|---|---|---|---|---|---|---|---|---|
| | | | | X | Y | Z | | |
| 1 | 16768 | 0.09* | 9.03 | -32 | 22 | 4 | L | Insula |
| | | 0.07* | 7.43 | -52 | 10 | 14 | L | Inferior frontal gyrus (pars opercularis) |
| | | 0.06* | 7.21 | -40 | 12 | -4 | L | Insula |
| | | 0.06* | 7.19 | -40 | 0 | -4 | L | Insula |
| | | 0.06* | 6.51 | -56 | 8 | 30 | L | Precentral gyrus |
| | | 0.04 | 4.64 | -34 | 22 | -10 | L | Insula |
| | | 0.03 | 4.15 | -38 | -2 | 14 | L | Insula |
| 2 | 13728 | 0.08* | 8.64 | -4 | 20 | 40 | L | Middle cingulate gyrus |
| | | 0.05* | 5.60 | 4 | 18 | 48 | R | Supplementary motor area |
| | | 0.04* | 5.15 | 6 | 12 | 60 | R | Supplementary motor area |
| | | 0.04 | 4.65 | 0 | 4 | 62 | L/R | Supplementary motor area |
| | | 0.03 | 4.50 | -2 | 20 | 26 | L | Anterior cingulate gyrus |
| | | 0.03 | 4.03 | -2 | 4 | 30 | L | Anterior cingulate gyrus |
| 3 | 10160 | 0.05* | 6.26 | 42 | -2 | -6 | R | Insula |
| | | 0.05* | 5.78 | 36 | 26 | 2 | R | Insula |
| | | 0.04* | 5.17 | 50 | 8 | 28 | R | Inferior frontal gyrus (pars opercularis) |
| | | 0.04 | 4.58 | 42 | 14 | -6 | R | Insula |
| | | 0.03 | 4.47 | 52 | 6 | 38 | R | Precentral gyrus |
| | | 0.03 | 4.39 | 44 | 40 | 10 | R | Inferior frontal gyrus (pars triangularis) |
| | | 0.03 | 4.23 | 52 | 12 | 6 | R | Inferior frontal gyrus (pars opercularis) |
| | | 0.03 | 3.96 | 40 | 30 | -8 | R | Inferior frontal gyrus (pars orbitalis) |
| | | 0.03 | 3.55 | 60 | 12 | 14 | R | Inferior frontal gyrus (pars opercularis) |
| | | 0.03 | 3.54 | 58 | 16 | 24 | R | Inferior frontal gyrus (pars opercularis) |
| 4 | 6000 | 0.11* | 10.93 | -58 | -24 | 32 | L | Supramarginal gyrus |
| 5 | 5312 | 0.10* | 10.45 | 62 | -20 | 32 | R | Supramarginal gyrus |

| | | | | | | | | |
|---|---|---|---|---|---|---|---|---|
| 6 | 4488 | 0.08* | 8.54 | -44 | -70 | -4 | L | Inferior occipital gyrus |
| 7 | 4280 | 0.06* | 6.89 | 50 | -62 | -8 | R | Inferior temporal gyrus |
| | | 0.04* | 5.37 | 48 | -70 | 0 | R | Inferior temporal gyrus |
| 8 | 2208 | 0.05* | 5.80 | -14 | -96 | -6 | L | Inferior occipital gyrus |
| | | 0.04* | 5.18 | -30 | -94 | 0 | L | Middle occipital gyrus |
| 9 | 1584 | 0.05* | 5.83 | -40 | -46 | 56 | L | Inferior parietal lobule |
| 10 | 1528 | 0.04 | 4.70 | 36 | -86 | 4 | R | Middle occipital gyrus |
| | | 0.04 | 4.61 | 34 | -88 | -2 | R | Inferior occipital gyrus |
| 11 | 1464 | 0.05* | 5.54 | -12 | -12 | 8 | L | Thalamus |
| | | 0.03 | 4.44 | -8 | -16 | -2 | L | Thalamus^ |
| 12 | 1440 | 0.04 | 4.59 | -42 | 42 | 8 | L | Inferior frontal gyrus (pars tringularis) |
| | | 0.03 | 3.74 | -44 | 32 | 14 | L | Inferior frontal gyrus (pars tringularis) |
| | | 0.03 | 3.60 | -34 | 50 | 14 | L | Middle frontal gyrus |
| 13 | 1280 | 0.04* | 5.24 | 34 | -54 | 58 | R | Superior parietal lobule |
| 14 | 1152 | 0.03 | 4.45 | -6 | -28 | -18 | L | Brainstem^ |
| | | 0.03 | 4.15 | -2 | -32 | -4 | L | Brainstem^ |
| | | 0.03 | 3.90 | 2 | -30 | -6 | R | Brainstem^ |
| 15 | 944 | 0.03 | 4.37 | 16 | 4 | 2 | R | Pallidum |
| 16 | 928 | 0.04* | 5.01 | 18 | -94 | -4 | R | Calcarine fissure |

MNI = Montreal Neurological Institute (MNI) standard stereotaxic space
* Indicates the peaks/sub-peaks withstanding the voxel-level FWE p < .05 threshold.
^ The anatomical region was labelled based on visual inspection of the ALE map because the coordinates fall outside the Automated Anatomical Labelling Atlas.

**Table S1.3.3.** Significant ALE clusters (cluster extent-based FWE threshold, p< .05) of convergent activation across 70 **empathy for emotions** experiments. Anatomical labels are derived from the Automatic Anatomical Labelling Atlas.

| Cluster | Volume (mm3) | ALE value | Z value | Peak MNI coordinates | | | Hemisphere | Anatomical region |
|---|---|---|---|---|---|---|---|---|
| | | | | X | Y | Z | | |
| 1 | 6952 | 0.05* | 6.54 | -38 | 24 | -4 | L | Inferior frontal gyrus (pars orbitalis) |
| | | 0.04 | 4.74 | -48 | 28 | -8 | L | Inferior frontal gyrus (pars orbitalis) |
| | | 0.03 | 4.55 | -46 | 18 | -16 | L | Superior temporal pole |
| | | 0.03 | 4.51 | -52 | 24 | 12 | L | Inferior frontal gyrus (pars triangularis) |
| | | 0.03 | 3.76 | -48 | 20 | 4 | L | Inferior frontal gyrus (pars triangularis) |
| 2 | 5200 | 0.04* | 5.59 | 50 | 30 | -6 | R | Inferior frontal gyrus (pars orbitalis) |
| | | 0.04* | 4.98 | 52 | 22 | 20 | R | Inferior frontal gyrus (pars triangularis) |
| | | 0.03 | 3.84 | 56 | 24 | 4 | R | Inferior frontal gyrus (pars triangularis) |
| 3 | 3280 | 0.04* | 5.15 | -48 | -72 | -2 | L | Inferior occipital gyrus |
| | | 0.04 | 4.71 | -46 | -76 | -10 | L | Inferior occipital gyrus |
| | | 0.03 | 4.57 | -42 | -78 | -12 | L | Inferior occipital gyrus |
| | | 0.03 | 4.28 | -36 | -88 | 8 | L | Middle occipital gyrus |
| | | 0.03 | 4.17 | -34 | -90 | -8 | L | Inferior occipital gyrus |
| 4 | 3080 | 0.05* | 6.03 | -4 | 22 | 50 | L | Supplementary motor area |
| | | 0.02 | 3.46 | 2 | 8 | 54 | R | Supplementary motor area |
| | | 0.02 | 3.31 | 6 | 10 | 50 | R | Supplementary motor area |
| 5 | 2360 | 0.04* | 5.50 | 46 | -74 | -8 | R | Inferior occipital gyrus |
| | | 0.03 | 4.43 | 50 | -64 | -2 | R | Inferior temporal gyrus |
| 6 | 2256 | 0.04 | 4.88 | 54 | -24 | -8 | R | Middle temporal gyrus |
| | | 0.03 | 4.28 | 52 | -34 | -2 | R | Middle temporal gyrus |
| 7 | 1488 | 0.04* | 5.62 | -4 | -28 | -6 | L | Brainstem^ |
| 8 | 1456 | 0.03 | 4.43 | 50 | 6 | 34 | R | Precentral gyrus |
| | | 0.03 | 3.94 | 46 | 4 | 46 | R | Precentral gyrus |
| 9 | 1432 | 0.04* | 5.17 | -6 | -50 | 32 | L | Posterior cingulate gyrus |
| | | 0.04 | 4.92 | 0 | -58 | 32 | L/R | Precuneus |
| | | 0.03 | 3.53 | 8 | -54 | 26 | R | Precuneus |
| 10 | 1080 | 0.03 | 4.46 | 48 | 14 | -36 | R | Middle temporal pole |

| | | | | | | | | |
|---|---|---|---|---|---|---|---|---|
| | | 0.03 | 3.96 | 52 | 8 | -34 | R | Inferior temporal gyrus |
| 11 | 952 | 0.05* | 6.36 | -12 | 6 | 12 | L | Caudate |
| 12 | 944 | 0.04* | 5.23 | -6 | 54 | 22 | L | Medial frontal gyrus |
| 13 | 904 | 0.05* | 5.78 | -18 | -8 | -16 | L | Hippocampus |
| 14 | 784 | 0.04* | 5.19 | -48 | 10 | -34 | L | Inferior temporal gyrus |
| 15 | 776 | 0.03 | 4.09 | -52 | -60 | 20 | L | Middle temporal gyrus |
| | | 0.03 | 3.85 | -52 | -60 | 8 | L | Middle temporal gyrus |

MNI = Montreal Neurological Institute (MNI) standard stereotaxic space
* Indicates the peaks/sub-peaks withstanding the voxel-level FWE p < .05 threshold. This analysis revealed two additional clusters in the middle cingulate gyrus ( [0, -18, 38], ALE value 0.04, Z value 5.65, cluster size 56 $mm^3$) and the right insula ( [30, 22, -14], ALE value 0.04, Z value 5.00, cluster size 8 $mm^3$) which did not withstand the cluster extent-based threshold.
^ The anatomical region was labelled based on visual inspection of the ALE map because the coordinates fall outside the Automated Anatomical Labelling Atlas.

**Table S1.3.4.** Significant clusters revealed by the conjunction and contrast analyses (p < .001 uncorrected, min. cluster size 200 $mm^3$) between **empathy for pain (85)** and **empathy for emotions** (70) experiments. Anatomical labels are derived from the Automatic Anatomical Labelling Atlas.

| Cluster | Volume (mm3) | ALE value | Z value | Peak MNI coordinates | | | Hemisphere | Anatomical region |
|---|---|---|---|---|---|---|---|---|
| | | | | X | Y | Z | | |
| **Empathy for Emotions & Empathy for Pain (conjunction)** | | | | | | | | |
| 1 | 2240 | 0.04 | | -30 | 26 | 2 | L | Insula |
| | | 0.04 | | -36 | 24 | 0 | L | Insula |
| | | 0.03 | | -36 | 20 | -8 | L | Insula |
| | | 0.03 | | -46 | 18 | 4 | L | Inferior frontal gyrus (pars triangularis) |
| 2 | 2208 | 0.05 | | -4 | 20 | 48 | L | Supplementary motor area |
| | | 0.02 | | 2 | 8 | 56 | R | Supplementary motor area |
| | | 0.02 | | 6 | 10 | 50 | R | Supplementary motor area |
| 3 | 1600 | 0.04 | | 42 | 24 | -2 | R | Insula |
| | | 0.03 | | 42 | 28 | -8 | R | Inferior frontal gyrus (pars orbitalis) |
| 4 | 1224 | 0.04 | | -48 | -72 | -2 | L | Inferior occipital gyrus |

| | | | | | | | | |
|---|---|---|---|---|---|---|---|---|
| 5 | 1016 | 0.03 | | 50 | -64 | -2 | R | Inferior temporal gyrus |
| | | 0.03 | | 48 | -70 | -6 | R | Inferior temporal gyrus |
| 6 | 736 | 0.03 | | 50 | 6 | 34 | R | Precentral gyrus |
| 7 | 616 | 0.03 | | -2 | -32 | -4 | L | Brainstem^ |
| | | 0.03 | | 2 | -30 | -6 | R | Brainstem^ |
| 8 | 88 | 0.03 | | -32 | -92 | -4 | L | Middle occipital gyrus |
| 9 | 16 | 0.02 | | 48 | 16 | 2 | R | Inferior frontal gyrus (pars opercularis) |
| 10 | 8 | 0.02 | | -34 | -90 | 0 | L | Middle occipital gyrus |
| **Empathy for Emotions - Empathy for Pain** | | | | | | | | |
| 1 | 672 | | 3.72 | -0.7 | -52.2 | 31.3 | L | Posterior cingulate gyrus |
| | | | 3.54 | -7 | -52.5 | 33.3 | L | Posterior cingulate gyrus |
| 2 | 640 | | 3.72 | 47.8 | 13.4 | -36.4 | R | Middle temporal pole |
| 3 | 464 | | 3.72 | -52.3 | -62.3 | 22.3 | L | Middle temporal gyrus |
| | | | 3.54 | -51.3 | -61.6 | 15.6 | L | Middle temporal gyrus |
| 4 | 408 | | 3.54 | -49.3 | 6 | -34.7 | L | Inferior temporal gyrus |
| | | | 3.35 | -49 | 6 | -30 | L | Middle temporal gyrus |
| | | | 3.16 | -50.2 | 10 | -35.2 | L | Inferior temporal gyrus |
| 5 | 256 | | 3.72 | -48 | 30 | -8.5 | L | Inferior frontal gyrus (pars orbitalis) |
| | | | 3.54 | -47 | 26 | -7 | L | Inferior frontal gyrus (pars orbitalis) |
| **Empathy for Pain - Empathy for Emotions** | | | | | | | | |
| 1 | 4032 | | 3.72 | -59.1 | -25.3 | 34.4 | L | Supramarginal gyrus |
| 2 | 2128 | | 3.72 | -43.8 | -0.1 | 4.1 | L | Insula |
| 3 | | | 3.72 | -1.2 | 19.4 | 36.5 | L | Middle cingulate gyrus |
| 4 | 608 | | 3.72 | 60.8 | -19.2 | 30.4 | R | Supramarginal gyrus |
| | | | 3.54 | 64 | -26 | 38 | R | Supramarginal gyrus |
| 5 | 4032 | | 3.72 | 41.9 | 0.6 | -3.5 | R | Insula |
| 6 | 2128 | | 3.72 | 52.3 | -57.7 | -10.9 | R | Inferior temporal gyrus |
| 7 | | | 3.72 | -45.7 | -62 | -4.2 | L | Inferior temporal gyrus |
| 8 | 608 | | 3.54 | -44 | 48 | 0 | L | Middle orbitofrontal cortex |
| | | | 3.35 | -43.2 | 47.5 | 4 | L | Inferior frontal gyrus (pars triangularis) |
| | | | 3.24 | -42 | 47 | 8 | L | Inferior frontal gyrus (pars triangularis) |

| | | | | | | | | |
|---|---|---|---|---|---|---|---|---|
| 9 | 2128 | | 3.72 | 35.8 | -49.3 | 53.3 | R | Inferior parietal lobule |
| | | | 3.54 | 38 | -50 | 58 | R | Superior parietal lobule |
| 10 | 608 | | 3.72 | -36.7 | 16.7 | 6 | L | Insula |
| | | | 3.54 | -34 | 16 | 2 | L | Insula |

MNI = Montreal Neurological Institute (MNI) standard stereotaxic space
^ The anatomical region was labelled based on visual inspection of the ALE map because the coordinates fall outside the Automated Anatomical Labelling Atlas.

**Table S1.4.** Significant ALE clusters (cluster extent-based FWE threshold, $p < .05$) of convergent activation across 69 **moral reasoning** experiments. Anatomical labels are derived from the Automatic Anatomical Labelling Atlas.

| **Cluster** | **Volume (mm3)** | **ALE value** | **Z value** | **Peak MNI coordinates** | | | **Hemisphere** | **Anatomical region** |
|---|---|---|---|---|---|---|---|---|
| | | | | **X** | **Y** | **Z** | | |
| 1 | 11848 | 0.05* | 6.46 | -2 | 58 | 28 | L | Medial superior frontal gyrus |
| | | 0.04* | 5.88 | -6 | 52 | 10 | L | Medial superior frontal gyrus |
| | | 0.04* | 5.36 | 6 | 60 | 14 | R | Medial superior frontal gyrus |
| | | 0.03 | 4.32 | -18 | 50 | 36 | L | Superior frontal gyrus |
| | | 0.03 | 3.95 | -2 | 36 | 18 | L | Anterior cingulate gyrus |
| | | 0.02 | 3.85 | -8 | 40 | 10 | L | Anterior cingulate gyrus |
| | | 0.02 | 3.50 | 8 | 50 | 6 | R | Medial superior frontal gyrus |
| 2 | 4880 | 0.06* | 7.38 | -48 | -62 | 24 | L | Middle temporal gyrus |
| | | 0.02 | 3.26 | -60 | -52 | 16 | L | Middle temporal gyrus |
| | | 0.02 | 3.23 | -60 | -52 | 20 | L | Middle temporal gyrus |
| 3 | 4352 | 0.06* | 8.06 | 0 | -56 | 32 | L/R | Precuneus |
| 4 | 2968 | 0.04* | 6.00 | -32 | 20 | -12 | L | Insula |
| | | 0.02 | 3.84 | -46 | 18 | -24 | L | Superior temporal pole |
| 5 | 2336 | 0.05* | 6.33 | 54 | 4 | -28 | R | Middle temporal gyrus |
| | | 0.02 | 3.67 | 60 | -6 | -32 | R | Inferior temporal gyrus |
| 6 | 1352 | 0.03 | 4.27 | 4 | 58 | -6 | R | Medial orbitofrontal cortex |
| | | 0.03 | 4.20 | 0 | 54 | -6 | L/R | Medial orbitofrontal cortex |
| | | 0.02 | 3.71 | -2 | 50 | -16 | L | Gyrus rectus |

| | | | | | | | | |
|---|---|---|---|---|---|---|---|---|
| 7 | 1080 | 0.03 | 4.56 | -62 | -16 | -20 | L | Middle temporal gyrus |
| 8 | 1032 | 0.03 | 4.35 | -50 | 16 | 16 | L | Inferior frontal gyrus (pars opercularis) |
| | | 0.03 | 4.02 | -52 | 24 | 14 | L | Inferior frontal gyrus (pars triangularis) |
| 9 | 960 | 0.03 | 4.65 | 50 | -64 | 6 | R | Middle temporal gyrus |
| 10 | 864 | 0.03 | 4.29 | -52 | -66 | 8 | L | Middle temporal gyrus |
| 11 | 808 | 0.03 | 4.33 | -40 | 12 | 48 | L | Middle frontal gyrus |

MNI = Montreal Neurological Institute (MNI) standard stereotaxic space
* Indicates the peaks/sub-peaks withstanding the voxel-level FWE p < .05 threshold.

**Table S1.5.** Overlay conjunction of **all social sub-domains**. All brain regions consistently activated by at least two of the four social sub-domains are listed. X indicates which social sub-domains reliably engaged each specified brain region.

| Anatomical region | Theory of Mind | Trait Inference | Empathy (Pain and/or Emotions) | Moral Reasoning |
|---|---|---|---|---|
| **All four social domains** | | | | |
| Left IFG (pars orbitalis) | X | X | X | X |
| mPFC | X | X | X | X |
| Precuneus | X | X | X | X |
| Left pSTG/AG* | X | X | X | X |
| **Three social domains** | | | | |
| Left IFG (pars opercularis) | X | | X | X |
| Right IFG | X | X | X | |
| SMA | X | X | X | |
| Medial OFC | X | X | | X |
| Left anterior MTG | X | X | | X |
| Right anterior MTG | X | | X | X |
| Right pMTG/ITG | X | | X | X |
| **Two social domains** | | | | |
| Right IFG (pars opercularis) | X | | X | |

| | | | | |
|---|---|---|---|---|
| Left Precentral gyrus | X | | | X |
| Right Precentral gyrus | X | | X | |
| Right pSTG/AG* | X | X | | |
| Right pMTG | X | X | | |
| Left TP | X | | X | |
| Left pMTG/ITG | | | X | X |

Note * corresponds to a brain region typically called the 'temporo-parietal junction' in the social neuroscience literature.

**Table S2.** Significant ALE clusters (cluster extent-based FWE threshold, p< .05) of convergent activation across 96 **semantic control** experiments which contrasted an experimental condition with increased semantic demands with a less demanding control condition. Anatomical labels are derived from the Automatic Anatomical Labelling Atlas.

| **Cluster** | **Volume (mm3)** | **ALE value** | **Z value** | **Peak MNI coordinates** | | | **Hemisphere** | **Anatomical region** |
|---|---|---|---|---|---|---|---|---|
| | | | | **X** | **Y** | **Z** | | |
| 1 | 25120 | 0.07* | 8.50 | -48 | 26 | 18 | L | Inferior frontal gyrus (pars triangularis) |
| | | 0.06* | 8.03 | -46 | 18 | 28 | L | Inferior frontal gyrus (pars triangularis) |
| | | 0.06* | 7.61 | -48 | 30 | 0 | L | Inferior frontal gyrus (pars triangularis) |
| | | 0.04* | 5.67 | -40 | 6 | 26 | L | Inferior frontal gyrus (pars opercularis) |
| | | 0.03* | 4.95 | -46 | 6 | 46 | L | Precentral gyrus |
| | | 0.03 | 4.90 | -44 | 10 | 38 | L | Middle frontal gyrus |
| | | 0.03 | 4.80 | -32 | 22 | -6 | L | Insula |
| | | 0.03 | 4.25 | -44 | 42 | -6 | L | Inferior frontal gyrus (pars orbitalis) |
| | | 0.02 | 3.74 | -36 | 28 | -16 | L | Inferior frontal gyrus (pars orbitalis) |
| 2 | 10200 | 0.05* | 6.73 | -2 | 20 | 50 | L | Supplementary motor area |
| | | 0.05* | 6.18 | -2 | 26 | 44 | L | Medial superior frontal gyrus |
| | | 0.04* | 5.61 | 8 | 28 | 36 | R | Cingulate gyrus |
| 3 | 6624 | 0.04* | 5.97 | -58 | -42 | -2 | L | Middle temporal gyrus |
| | | 0.04* | 5.21 | -48 | -58 | -18 | L | Fusiform gyrus |
| | | 0.03 | 4.42 | -54 | -56 | 2 | L | Middle temporal gyrus |
| | | 0.03 | 4.11 | -50 | -60 | -8 | L | Inferior temporal gyrus |
| 4 | 3504 | 0.05* | 6.19 | 36 | 26 | -8 | R | Inferior frontal gyrus (pars orbitalis) |
| | | 0.04* | 5.94 | 40 | 26 | -14 | R | Inferior frontal gyrus (pars orbitalis) |

| | | | | | | | | |
|---|---|---|---|---|---|---|---|---|
| 5 | 2264 | 0.03 | 4.87 | -34 | -66 | 42 | L | Inferior parietal lobule |
| | | 0.03 | 4.28 | -30 | -56 | 46 | L | Inferior parietal lobule |
| 6 | 2184 | 0.04* | 5.37 | 54 | 24 | 26 | R | Inferior frontal gyrus (pars triangularis) |
| | | 0.02 | 3.28 | 38 | 20 | 18 | R | Inferior frontal gyrus (pars opercularis) |
| 7 | 960 | 0.03 | 4.56 | -44 | -48 | 44 | L | Inferior parietal lobule |

MNI = Montreal Neurological Institute (MNI) standard stereotaxic space
* Indicates the peaks/sub-peaks withstanding the voxel-level FWE p < .05 threshold.

**Table S3.1.** Significant clusters revealed by the conjunction and contrast analyses ($p < .001$ uncorrected, min. cluster size 200 $mm^3$) between **semantic control** (96) and **ToM** (136) experiments. Anatomical labels are derived from the Automatic Anatomical Labelling Atlas.

| Cluster | Volume (mm3) | ALE value | Z value | Peak MNI coordinates | | | Hemisphere | Anatomical region |
|---|---|---|---|---|---|---|---|---|
| | | | | X | Y | Z | | |
| **Semantic Control & ToM (conjunction)** | | | | | | | | |
| 1 | 7376 | 0.06 | | -48 | 28 | -2 | L | Inferior frontal gyrus (pars orbitalis) |
| | | 0.06 | | -52 | 20 | 22 | L | Inferior frontal gyrus (pars triangularis) |
| | | 0.05 | | -50 | 28 | 4 | L | Inferior frontal gyrus (pars triangularis) |
| | | 0.05 | | -50 | 22 | 18 | L | Inferior frontal gyrus (pars triangularis) |
| 2 | 2168 | 0.04 | | -58 | -42 | -2 | L | Middle temporal gyrus |
| | | 0.03 | | -54 | -56 | 4 | L | Middle temporal gyrus |
| 3 | 2024 | 0.05 | | -2 | 18 | 52 | L | Supplementary motor area |
| 4 | 768 | 0.04 | | 54 | 24 | 24 | R | Inferior frontal gyrus (pars triangularis) |
| 5 | 728 | 0.03 | | -44 | -54 | -18 | L | Fusiform gyrus |
| 6 | 664 | 0.03 | | -46 | 6 | 46 | L | Precentral gyrus |
| 7 | 128 | 0.02 | | 46 | 26 | -10 | R | Inferior frontal gyrus (pars orbitalis) |
| 8 | 32 | 0.02 | | -50 | 14 | -18 | L | Superior temporal pole |
| **ToM - Semantic Control** | | | | | | | | |
| 1 | 9464 | | 3.72 | 54.1 | -1.4 | -21.6 | R | Middle temporal gyrus |
| 2 | 9432 | | 3.72 | 53.4 | -53.6 | 19.4 | R | Middle temporal gyrus |
| | | | 3.16 | 50 | -44 | 7 | R | Middle temporal gyrus |
| 3 | 8184 | | 3.72 | -51 | -55.9 | 19.7 | L | Middle temporal gyrus |
| 4 | 8080 | | 3.72 | 1.7 | -55 | 33.5 | R | Precuneus |

| | | | | | | | | |
|---|---|---|---|---|---|---|---|---|
| | | | 3.54 | -6.8 | -49.2 | 48.4 | L | Precuneus |
| 5 | 5584 | | 3.72 | -56.9 | -9.7 | -17.2 | L | Middle temporal gyrus |
| 6 | 5096 | | 3.72 | 3.5 | 56.7 | 21.9 | R | Medial superior frontal gyrus |
| | | | 3.54 | -7.5 | 56.5 | 14 | L | Medial superior frontal gyrus |
| | | | 3.24 | -1 | 55 | 8 | L | Medial superior frontal gyrus |
| 7 | 2056 | | 3.72 | -22.4 | -77.7 | -35.1 | L | Cerebellum |
| 8 | 1600 | | 3.72 | -10.3 | 53.6 | 38.3 | L | Superior frontal gyrus |
| | | | 3.54 | -17 | 50 | 33 | L | Superior frontal gyrus |
| 9 | 592 | | 3.72 | 49.9 | -33.1 | -3.8 | R | Middle temporal gyrus |
| 10 | 568 | | 3.72 | 54.7 | 28.2 | 3.8 | R | Inferior frontal gyrus (pars triangularis) |
| 11 | 248 | | 3.72 | -4 | 44 | -17 | L | Gyrus rectus |
| | | | 3.24 | 0 | 48 | -18 | L/R | Gyrus rectus |
| **Semantic Control - ToM** | | | | | | | | |
| 1 | 4032 | | 3.72 | -43.6 | 29.1 | 14.8 | L | Inferior frontal gyrus (pars triangularis) |
| 2 | 2128 | | 3.72 | 2.5 | 25.5 | 38.4 | R | Middle cingulate gyrus |
| | | | 3.54 | 11 | 26 | 31 | R | Middle cingulate gyrus |
| 3 | 608 | | 3.72 | -32.2 | -67.3 | 45.2 | L | Inferior parietal gyrus |
| 4 | 320 | | 3.72 | -35.3 | 16.8 | -6.4 | L | Insula |
| | | | 3.16 | -32 | 24 | -10 | L | Inferior frontal gyrus (pars orbitalis) |

MNI = Montreal Neurological Institute (MNI) standard stereotaxic space

**Table S3.1.2.** Significant clusters revealed by the conjunction and contrast analyses ($p < .001$ uncorrected, min. cluster size 200 $mm^3$) between **semantic control** (96) and **false belief reasoning** (43) experiments. Anatomical labels are derived from the Automatic Anatomical Labelling Atlas.

| Cluster | Volume (mm3) | ALE value | Z value | Peak MNI coordinates | | | Hemisphere | Anatomical region |
|---|---|---|---|---|---|---|---|---|
| | | | | X | Y | Z | | |
| **False Belief Reasoning & Semantic Control (Conjunction)** | | | | | | | | |
| 1 | 200 | 0.02 | | -60 | -34 | -4 | L | Middle temporal gyrus |
| **False Belief Reasoning – Semantic Control** | | | | | | | | |
| 1 | 5560 | | 3.72 | 3.1 | -55.6 | 33.8 | R | Precuneus |

| | | | | | | | | |
|---|---|---|---|---|---|---|---|---|
| 2 | 5256 | | 3.72 | 56.3 | -10.2 | -17.6 | R | Middle temporal gyrus |
| | | | 0 | 56.8 | 2 | -23.2 | R | Middle temporal gyrus |
| | | | 3.35 | 54 | 6 | -18 | R | Middle temporal pole |
| 3 | 5088 | | 3.72 | -51.7 | -57.5 | 23.3 | L | Middle temporal gyrus |
| 4 | 4952 | | 3.72 | 54 | -53.3 | 23.9 | R | Superior temporal gyrus |
| 5 | 2840 | | 3.72 | 0.4 | 55.7 | 19 | L/R | Medial superior frontal gyrus |
| | | | 3.54 | 2 | 58 | 30 | L/R | Medial superior frontal gyrus |
| 6 | 1576 | | 3.72 | -61.4 | -22.4 | -10.5 | L | Middle temporal gyrus |
| 7 | 1144 | | 3.72 | -24.9 | -78.9 | -34.6 | L | Cerebellum |
| | | | 3.54 | -23 | -81.5 | -40.5 | L | Cerebellum |
| 8 | 1128 | | 3.72 | -54 | 7.6 | -24.8 | L | Middle temporal gyrus |
| | | | 3.54 | -53.1 | 0 | -25.8 | L | Middle temporal gyrus |
| 9 | 864 | | 3.72 | 23.6 | 26.2 | 44.8 | R | Middle frontal gyrus |
| **Semantic Control - False Belief Reasoning** | | | | | | | | |
| 1 | 7096 | | 3.35 | -46 | 28 | 16.4 | L | Inferior frontal gyrus (pars triangularis) |
| | | | 3.54 | -52 | 12 | 40 | L | Precentral gyrus |
| 2 | 1336 | | 3.72 | -1.9 | 29.3 | 39 | L | Medial superior frontal gyrus |
| | | | 3.54 | 2.3 | 35 | 29.7 | R | Middle cingulate gyrus |
| 3 | 272 | | 3.54 | 50.2 | 33.4 | 21.6 | R | Inferior frontal gyrus (pars triangularis) |

MNI = Montreal Neurological Institute (MNI) standard stereotaxic space

**Table S3.2.** Significant clusters revealed by the conjunction and contrast analyses ($p < .001$ uncorrected, min. cluster size 200 $mm^3$) between **semantic control** (96) and **trait inference** (40) experiments. Anatomical labels are derived from the Automatic Anatomical Labelling Atlas.

| Cluster | Volume (mm3) | ALE value | Z value | Peak MNI coordinates | | | Hemisphere | Anatomical region |
|---|---|---|---|---|---|---|---|---|
| | | | | X | Y | Z | | |
| **Semantic Control & Trait Inference (conjunction)** | | | | | | | | |
| 1 | 2664 | 0.03 | | -46 | 24 | -12 | L | Inferior frontal gyrus (pars orbitalis) |
| | | 0.02 | | -50 | 28 | 2 | L | Inferior frontal gyrus (pars triangularis) |
| 2 | 760 | 0.03 | | 48 | 28 | -12 | R | Inferior frontal gyrus (pars orbitalis) |

| | | | | | | | | |
|---|---|---|---|---|---|---|---|---|
| 3 | 552 | 0.02 | | -8 | 20 | 56 | L | Supplementary motor area |
| | | 0.02 | | 0 | 18 | 50 | L/R | Supplementary motor area |
| 4 | 160 | 0.02 | | -6 | 32 | 52 | L | Medial superior frontal gyrus |
| | | 0.02 | | -6 | 32 | 44 | L | Medial superior frontal gyrus |
| 5 | 16 | 0.02 | | -8 | 30 | 50 | L | Medial superior frontal gyrus |
| 6 | 8 | 0.01 | | -6 | 28 | 54 | L | Medial superior frontal gyrus |
| **Trait Inference – Semantic Control** | | | | | | | | |
| 1 | 5192 | | 3.72 | -1.4 | 57.4 | 25.4 | L | Medial superior frontal gyrus |
| | | | 3.54 | 5.1 | 53.4 | 41.1 | R | Medial superior frontal gyrus |
| 2 | 2376 | | 3.72 | 0 | -54.4 | 27.5 | L/R | Precuneus |
| 3 | 1120 | | 3.72 | -9.7 | 34.2 | 56.9 | L | Medial superior frontal gyrus |
| | | | 3.54 | -8 | 40 | 48 | L | Medial superior frontal gyrus |
| 4 | 600 | | 3.72 | 50.2 | -57.6 | 25.6 | R | Angular gyrus |
| 5 | 416 | | 3.72 | 3 | 56 | -2 | R | Medial orbitofrontal cortex |
| | | | 3.54 | 4.5 | 60 | -3.5 | R | Medial orbitofrontal cortex |
| 6 | 400 | | 3.54 | -64 | -13 | -20.6 | L | Middle temporal gyrus |
| | | | 3.35 | -65.3 | -17.3 | -17.3 | L | Middle temporal gyrus |
| **Semantic Control - Trait Inference** | | | | | | | | |
| 1 | 4992 | | 3.72 | -45.1 | 25.6 | 17.2 | L | Inferior frontal gyrus (pars triangularis) |
| | | | 3.09 | -50.7 | 22 | 25.3 | L | Inferior frontal gyrus (pars triangularis) |

MNI = Montreal Neurological Institute (MNI) standard stereotaxic space

**Table S3.3.** Significant clusters revealed by the conjunction and contrast analyses ($p < .001$ uncorrected, min. cluster size 200 $mm^3$) between **semantic control** (96) and **empathy for emotions** (70) experiments. Anatomical labels are derived from the Automatic Anatomical Labelling Atlas.

| Cluster | Volume (mm3) | ALE value | Z value | Peak MNI coordinates X | Y | Z | Hemisphere | Anatomical region |
|---|---|---|---|---|---|---|---|---|
| **Semantic Control & Empathy Emotions (conjunction)** | | | | | | | | |
| 1 | 4480 | 0.04 | | -48 | 28 | -8 | L | Inferior frontal gyrus (pars orbitalis) |
| | | 0.03 | | -42 | 26 | -4 | L | Inferior frontal gyrus (pars orbitalis) |
| | | 0.03 | | -52 | 24 | 12 | L | Inferior frontal gyrus (pars triangularis) |
| | | 0.03 | | -32 | 22 | -4 | L | Insula |
| | | 0.03 | | -48 | 20 | -14 | L | Superior temporal pole |
| | | 0.03 | | -48 | 20 | 4 | L | Inferior frontal gyrus (pars triangularis) |
| 2 | 2584 | 0.05 | | -2 | 22 | 50 | L | Supplementary motor area |
| | | 0.02 | | 0 | 8 | 54 | L/R | Supplementary motor area |
| 3 | 696 | 0.03 | | 42 | 26 | -8 | R | Inferior frontal gyrus (pars orbitalis) |
| | | 0.03 | | 40 | 24 | -4 | R | Insula |
| | | 0.03 | | 46 | 28 | -12 | R | Inferior frontal gyrus (pars orbitalis) |
| 4 | 456 | 0.03 | | 54 | 24 | 22 | R | Inferior frontal gyrus (pars triangularis) |
| 5 | 16 | 0.02 | | -54 | -58 | 6 | L | Middle temporal gyrus |
| 6 | 8 | 0.02 | | 38 | 24 | 0 | R | Insula |
| **Empathy Emotions - Semantic Control** | | | | | | | | |
| 1 | 1896 | | 3.54 | 48.1 | -69.5 | -4.5 | R | Inferior temporal gyrus |
| 2 | 808 | | 3.72 | -0.6 | -29.7 | -9.1 | L | Brainstem^ |
| 3 | 736 | | 3.72 | 54.5 | -29.2 | -3.7 | R | Middle temporal gyrus |
| | | | 3.54 | 52 | -40 | -2 | R | Middle temporal gyrus |
| 4 | 544 | | 3.72 | -48 | -72.5 | 4 | L | Middle occipital gyrus |
| | | | 3.54 | -44 | -68 | 6 | L | Inferior occipital gyrus |
| | | | 3.35 | -44 | -70 | -4 | L | Inferior occipital gyrus |
| 5 | 488 | | 3.72 | 2.1 | -55.9 | 29.3 | R | Precuneus |
| 6 | 448 | | 3.72 | 47.3 | 26.8 | -2 | R | Inferior frontal gyrus (pars orbitalis) |
| | | | 3.16 | 52 | 31 | 1 | R | Inferior frontal gyrus (pars triangularis) |

| | | | | | | | | |
|---|---|---|---|---|---|---|---|---|
| 7 | 360 | | 3.72 | -28.3 | 30.7 | -1 | L | Insula |
| **Semantic Control - Empathy Emotions** | | | | | | | | |
| 1 | 688 | | 3.54 | -44 | 29.5 | 8.5 | L | Inferior frontal gyrus (pars triangularis) |
| 2 | 472 | | 3.72 | -41.6 | 19.6 | 21.6 | L | Inferior frontal gyrus (pars triangularis) |
| | | | 3.16 | -54 | 30 | 23 | L | Inferior frontal gyrus (pars triangularis) |
| 3 | 448 | | 3.72 | -55.5 | -40 | -7.5 | L | Middle temporal gyrus |
| | | | 3.54 | -57 | -44 | -8 | L | Middle temporal gyrus |
| | | | 3.35 | -62 | -36 | -2 | L | Middle temporal gyrus |
| 4 | 360 | | 3.72 | -34.9 | -69.6 | 40.4 | L | Middle occipital gyrus |
| | | | 3.35 | -38 | -64 | 46 | L | Angular gyrus |
| | | | 3.16 | -30 | -66 | 40 | L | Middle occipital gyrus |
| 5 | 296 | | 3.54 | -1 | 34 | 34 | L | Medial superior frontal gyrus |
| | | | 3.35 | 2 | 28 | 38.3 | R | Middle cingulate gyrus |

MNI = Montreal Neurological Institute (MNI) standard stereotaxic space
^ The anatomical region was labelled based on visual inspection of the ALE map because the coordinates fall outside the Automated Anatomical Labelling Atlas.

**Table S3.4.** Significant clusters revealed by the conjunction and contrast analyses (p < .001 uncorrected, min. cluster size 200 $mm^3$) between **semantic control** (96) and **empathy for pain** (85) experiments. Anatomical labels are derived from the Automatic Anatomical Labelling Atlas.

| Cluster | Volume (mm3) | ALE value | Z value | Peak MNI coordinates | | | Hemisphere | Anatomical region |
|---|---|---|---|---|---|---|---|---|
| | | | | X | Y | Z | | |
| **Semantic Control & Empathy Pain (conjunction)** | | | | | | | | |
| 1 | 6200 | 0.04 | | -2 | 18 | 48 | L | Supplementary motor area |
| | | 0.04 | | -2 | 26 | 42 | L | Medial superior frontal gyrus |
| 2 | 1776 | 0.03 | | -34 | 20 | -4 | L | Insula |
| | | 0.03 | | -32 | 22 | -8 | L | Insula |
| | | 0.03 | | -42 | 24 | 0 | L | Inferior frontal gyrus (pars triangularis) |
| 3 | 1248 | 0.03 | | -50 | 12 | 22 | L | Inferior frontal gyrus (pars opercularis) |
| | | 0.03 | | -44 | 6 | 26 | L | Inferior frontal gyrus (pars opercularis) |

| | | | | | | | | |
|---|---|---|---|---|---|---|---|---|
| | | 0.03 | | -54 | 16 | 10 | L | Inferior frontal gyrus (pars opercularis) |
| | | 0.02 | | -42 | 2 | 28 | L | Precentral gyrus |
| 4 | 960 | 0.03 | | 36 | 28 | -4 | R | Insula |
| | | 0.03 | | 40 | 30 | -10 | R | Inferior frontal gyrus (pars orbitalis) |
| 5 | 328 | 0.03 | | -50 | -60 | -6 | L | Inferior temporal gyrus |
| | | 0.03 | | -52 | -60 | 0 | L | Middle temporal gyrus |
| 6 | 264 | 0.03 | | -44 | 32 | 14 | L | Inferior frontal gyrus (pars triangularis) |
| | | 0.02 | | -44 | 38 | 6 | L | Inferior frontal gyrus (pars triangularis) |
| 7 | 8 | 0.02 | | -48 | 8 | 32 | L | Precentral gyrus |
| 8 | 8 | 0.02 | | -52 | 8 | 38 | L | Precentral gyrus |
| 9 | 8 | 0.02 | | -52 | 6 | 40 | L | Precentral gyrus |
| **Empathy Pain - Semantic Control** | | | | | | | | |
| 1 | 6400 | | 3.72 | -40.1 | 4.4 | 0.5 | L | Insula |
| | | | 3.54 | -58 | 6 | 16 | L | Inferior frontal gyrus (pars opercularis) |
| 2 | 5376 | | 3.72 | -59.4 | -24.3 | 30.9 | L | Supramarginal gyrus |
| 3 | 4800 | | 3.72 | 59.3 | -22.7 | 34.1 | R | Supramarginal gyrus |
| 4 | 3744 | | 3.72 | 50.3 | -63.2 | -5 | R | Inferior temporal gyrus |
| | | | 3.54 | 46 | -70 | 2 | R | Middle temporal gyrus |
| 5 | 2536 | | 3.72 | 41.4 | 2.2 | -2.9 | R | Insula |
| 6 | 2320 | | 3.72 | -44.5 | -69.4 | -1.7 | L | Inferior occipital gyrus |
| 7 | 888 | | 3.72 | 34.7 | -87.1 | 1 | R | Middle occipital gyrus |
| 8 | 864 | | 3.72 | -37.8 | -47 | 59.9 | L | Superior parietal lobule |
| 9 | 688 | | 3.72 | -56.1 | 5.5 | 29.7 | L | Precentral gyrus |
| 10 | 688 | | 3.72 | -2.1 | 7.7 | 35.9 | L | Middle cingulate gyrus |
| | | | 3.16 | -8 | 13 | 44 | L | Middle cingulate gyrus |
| 11 | 632 | | 3.72 | 35.7 | -51.9 | 59.4 | R | Superior parietal lobule |
| | | | 3.35 | 30 | -56.7 | 55.3 | R | Superior parietal lobule |
| 12 | 464 | | 3.72 | 52.3 | 9.5 | 6.8 | R | Rolandic operculum |
| 13 | 456 | | 3.72 | -24 | -96.5 | -2.5 | L | Middle occipital gyrus |
| 14 | 408 | | 3.72 | 16 | 0.1 | -2.6 | R | Pallidum |
| 15 | 408 | | 3.72 | 5.9 | 14.9 | 61.3 | R | Supplementary motor area |

| | | | | | | | | |
|---|---|---|---|---|---|---|---|---|
| 16 | 280 | | 3.72 | -4.8 | -29.8 | -16.5 | L | Brainstem^ |
| **Semantic Control - Empathy Pain** | | | | | | | | |
| 1 | 3808 | | 3.72 | -45 | 23 | 20.2 | L | Inferior frontal gyrus (pars triangularis) |
| | | | 3.35 | -46 | 24 | 34 | L | Middle frontal gyrus |
| 2 | 2552 | | 3.72 | -57.7 | -40.2 | -2.4 | L | Middle temporal gyrus |
| 3 | 568 | | 3.72 | -49.6 | 28 | -8.7 | L | Inferior frontal gyrus (pars orbitalis) |
| | | | 3.24 | -44.5 | 31 | -6 | L | Inferior frontal gyrus (pars orbitalis) |
| | | | 3.35 | -46 | 30 | -2 | L | Inferior frontal gyrus (pars orbitalis) |
| | | | 3.24 | -48 | 26 | 4 | L | Inferior frontal gyrus (pars triangularis) |
| 4 | 216 | | 3.72 | -34 | -66 | 48 | L | Inferior parietal lobule |

MNI = Montreal Neurological Institute (MNI) standard stereotaxic space
^ The anatomical region was labelled based on visual inspection of the ALE map because the coordinates fall outside the Automated Anatomical Labelling Atlas.

**Table S3.5.** Significant clusters revealed by the conjunction and contrast analyses ($p < .001$ uncorrected, min. cluster size 200 $mm^3$) between **semantic control** (96) and **moral reasoning** (69) experiments. Anatomical labels are derived from the Automatic Anatomical Labelling Atlas.

| **Cluster** | **Volume (mm3)** | **ALE value** | **Z value** | **Peak MNI coordinates** | | | **Hemisphere** | **Anatomical region** |
|---|---|---|---|---|---|---|---|---|
| | | | | **X** | **Y** | **Z** | | |
| **Semantic control & Moral reasoning (conjunction)** | | | | | | | | |
| 1 | 1048 | 0.03 | | -34 | 22 | -8 | L | Insula |
| | | 0.02 | | -36 | 26 | -16 | L | Inferior frontal gyrus (pars orbitalis) |
| 2 | 704 | 0.03 | | -50 | 18 | 16 | L | Inferior frontal gyrus (pars opercularis) |
| | | 0.03 | | -52 | 24 | 14 | L | Inferior frontal gyrus (pars triangularis) |
| 3 | 224 | 0.03 | | -42 | 10 | 46 | L | Precentral gyrus |
| **Moral reasoning - Semantic control** | | | | | | | | |
| 1 | 3448 | | 3.72 | -3 | 57 | 17.1 | L | Medial superior frontal gyrus |
| | | | 3.72 | 3.3 | 59.3 | 22.7 | R | Medial superior frontal gyrus |

| | | | | | | | |
|---|---|---|---|---|---|---|---|
| 2 | 2928 | 3.72 | 1.8 | -56.2 | 31.2 | L | Precuneus |
| 3 | 1728 | 3.72 | -45.8 | -60.1 | 18.8 | L | Middle temporal gyrus |
| | | 3.54 | -54 | -68 | 24 | L | Middle temporal gyrus |
| 4 | 1584 | 3.72 | 52.3 | 4.5 | -28.1 | R | Middle temporal gyrus |
| | | 3.54 | 48 | 2 | -24 | R | Middle temporal gyrus |
| 5 | 928 | 3.72 | 50 | -64.1 | 6.9 | R | Middle temporal gyrus |
| 6 | 472 | 3.72 | -50 | -70.5 | 10.5 | L | Middle temporal gyrus |
| 7 | 440 | 3.54 | 4.6 | 57.4 | -4.4 | R | Medial orbitofrontal cortex |
| | | | | **Semantic control - Moral reasoning** | | | |
| 1 | 7352 | 3.72 | -44.9 | 23.1 | 22.1 | L | Inferior frontal gyrus (pars triangularis) |
| 2 | 3392 | 3.72 | 0.1 | 23.4 | 43.9 | L/R | Medial superior frontal gyrus |
| | | 3.54 | 3.1 | 30.9 | 39.6 | R | Middle cingulate gyrus |
| 3 | 520 | 3.72 | -51.2 | -58 | -17.9 | L | Inferior temporal gyrus |
| | | 3.16 | -44 | -60 | -20 | L | Fusiform gyrus |
| 4 | 256 | 3.54 | 52 | 23 | 32 | R | Inferior frontal gyrus (pars triangularis) |
| | | 3.35 | 49 | 24 | 25 | R | Inferior frontal gyrus (pars triangularis) |
| | | 3.16 | 48 | 24 | 19 | R | Inferior frontal gyrus (pars triangularis) |

MNI = Montreal Neurological Institute (MNI) standard stereotaxic space

**Table S4.1.1.** Significant ALE clusters (cluster extent-based FWE threshold, p< .05) of convergent activation across 46 **explicit empathy for emotions** experiments. Anatomical labels are derived from the Automatic Anatomical Labelling Atlas.

| Cluster | Volume (mm3) | ALE value | Z value | Peak MNI coordinates | | | Hemisphere | Anatomical region |
|---|---|---|---|---|---|---|---|---|
| | | | | X | Y | Z | | |
| 1 | 4080 | 0.04* | 5.45 | -34 | 28 | -4 | L | Inferior frontal gyrus (pars orbitalis) |
| | | 0.03 | 4.73 | -48 | 18 | -18 | L | Superior temporal pole |
| | | 0.02 | 3.62 | -46 | 36 | -8 | L | Inferior frontal gyrus (pars orbitalis) |
| 2 | 3072 | 0.03* | 5.21 | 52 | 30 | -6 | R | Inferior frontal gyrus (pars orbitalis) |
| | | 0.03 | 4.22 | 48 | 28 | 4 | R | Inferior frontal gyrus (pars triangularis) |
| | | 0.03 | 4.18 | 58 | 26 | 14 | R | Inferior frontal gyrus (pars triangularis) |

| | | | | | | | | |
|---|---|---|---|---|---|---|---|---|
| | | 0.02 | 3.44 | 48 | 20 | 0 | R | Inferior frontal gyrus (pars opercularis) |
| 3 | 2328 | 0.03* | 5.14 | -4 | 54 | 22 | L | Medial superior frontal gyrus |
| | | 0.02 | 4.06 | -12 | 48 | 36 | L | Superior frontal gyrus |
| | | 0.02 | 3.68 | -2 | 48 | 36 | L | Medial superior frontal gyrus |
| | | 0.02 | 3.68 | -2 | 52 | 34 | L | Medial superior frontal gyrus |
| 4 | 1968 | 0.04* | 5.91 | -4 | 20 | 48 | L | Supplementary motor area |
| | | 0.02 | 3.66 | -6 | 20 | 62 | L | Supplementary motor area |
| | | 0.02 | 3.42 | -4 | 32 | 50 | L | Medial superior frontal gyrus |
| 5 | 1912 | 0.04* | 5.45 | 0 | -58 | 32 | L/R | Precuneus |
| | | 0.03* | 5.19 | -6 | -50 | 32 | L | Posterior cingulate gyrus |
| | | 0.02 | 4.06 | 8 | -54 | 26 | R | Precuneus |
| 6 | 1496 | 0.03 | 4.94 | 48 | 14 | -36 | R | Middle temporal pole |
| | | 0.03 | 4.44 | 52 | 8 | -34 | R | Inferior temporal gyrus |
| 7 | 1016 | 0.03 | 4.71 | -2 | -28 | -4 | L | Brainstem^ |
| 8 | 920 | 0.05* | 6.51 | -12 | 6 | 12 | L | Caudate |
| 9 | 912 | 0.04* | 5.38 | -48 | 10 | -34 | L | Inferior temporal gyrus |
| 10 | 808 | 0.04* | 6.23 | 0 | -18 | 38 | L/R | Middle cingulate gyrus |

MNI = Montreal Neurological Institute (MNI) standard stereotaxic space

* Indicates the peaks/sub-peaks withstanding the voxel-level FWE $p < .05$ threshold. This analysis revealed one additional cluster in the left thalamus ( [-4, -10, 8], ALE value 0.03, Z value 5.02, cluster size 16 $mm^3$) which did not withstand the cluster extent-based threshold.

^ The anatomical region was labelled based on visual inspection of the ALE map because the coordinates fall outside the Automated Anatomical Labelling Atlas.

**Table S4.1.2.** Significant ALE clusters (cluster extent-based FWE threshold, p< .05) of convergent activation across 23 **implicit empathy for emotions** experiments. Anatomical labels are derived from the Automatic Anatomical Labelling Atlas.

| Cluster | Volume (mm3) | ALE value | Z value | Peak MNI coordinates | | | Hemisphere | Anatomical region |
|---|---|---|---|---|---|---|---|---|
| | | | | X | Y | Z | | |
| 1 | 968 | 0.02 | 4.72 | 46 | 26 | -2 | R | Inferior frontal gyrus (pars orbitalis) |
| 2 | 968 | 0.02 | 4.70 | 48 | -38 | 4 | R | Middle temporal gyrus |
| 3 | 880 | 0.02* | 4.92 | -38 | 24 | -4 | L | Inferior frontal gyrus (pars orbitalis) |
| 4 | 784 | 0.02 | 4.42 | -54 | -64 | 8 | L | Middle temporal gyrus |
| | | 0.02 | 3.65 | -48 | -72 | -2 | L | Inferior occipital gyrus |
| 5 | 680 | 0.02 | 3.98 | 38 | -76 | -18 | R | Cerebellum |
| | | 0.02 | 3.64 | 44 | -74 | -6 | R | Inferior occipital gyrus |

MNI = Montreal Neurological Institute (MNI) standard stereotaxic space
* Indicates the peaks/sub-peaks withstanding the voxel-level FWE p < .05 threshold. This analysis revealed one additional cluster in the right fusiform gyrus ( [44, -48, -22], ALE value 0.03, Z value 4.97, cluster size 8 $mm^3$) which did not withstand the cluster extent-based threshold.

**Table S4.1.3.** Significant ALE clusters (cluster extent-based FWE threshold, p< .05) of convergent activation across 48 **explicit empathy for pain** experiments. Anatomical labels are derived from the Automatic Anatomical Labelling Atlas.

| Cluster | Volume (mm3) | ALE value | Z value | Peak MNI coordinates | | | Hemisphere | Anatomical region |
|---|---|---|---|---|---|---|---|---|
| | | | | X | Y | Z | | |
| 1 | 10336 | 0.05* | 7.02 | -32 | 22 | 6 | L | Insula |
| | | 0.04* | 6.36 | -40 | 12 | -6 | L | Insula |
| | | 0.04* | 5.81 | -52 | 10 | 10 | L | Inferior frontal gyrus (pars opercularis) |
| | | 0.03* | 5.26 | -58 | 10 | 28 | L | Precentral gyrus |
| | | 0.03* | 4.90 | -32 | 24 | -10 | L | Inferior frontal gyrus (pars orbitalis) |
| | | 0.03 | 4.57 | -42 | -4 | 2 | L | Insula |
| | | 0.02 | 3.88 | -38 | -2 | 14 | L | Insula |
| 2 | 9152 | 0.07* | 8.83 | -4 | 22 | 40 | L | Middle cingulate gyrus |
| | | 0.03 | 4.44 | 0 | 14 | 58 | L/R | Supplementary motor area |
| | | 0.02 | 4.24 | -2 | 18 | 26 | L | Anterior cingulate gyrus |

| | | | | | | | | |
|---|---|---|---|---|---|---|---|---|
| | | 0.02 | 4.12 | 0 | 4 | 64 | L/R | Supplementary motor area |
| | | 0.02 | 4.07 | 4 | 8 | 62 | R | Supplementary motor area |
| 3 | 3784 | 0.05* | 7.36 | -60 | -22 | 30 | L | Supramarginal gyrus |
| 4 | 2152 | 0.03 | 4.66 | -44 | 44 | 6 | L | Inferior frontal gyrus (pars triangularis) |
| | | 0.03 | 4.38 | -34 | 50 | 14 | L | Middle frontal gyrus |
| | | 0.02 | 3.31 | -36 | 50 | 24 | L | Middle frontal gyrus |
| 5 | 2144 | 0.05* | 7.10 | 62 | -22 | 34 | R | Supramarginal gyrus |
| 6 | 1488 | 0.03* | 5.02 | -46 | -68 | -2 | L | Inferior occipital gyrus |
| | | 0.02 | 4.30 | -50 | -60 | -2 | L | Middle temporal gyrus |
| 7 | 1104 | 0.03 | 4.44 | 36 | 30 | -2 | R | Insula |
| | | 0.02 | 4.25 | 32 | 22 | 6 | R | Insula |
| | | 0.02 | 3.50 | 50 | 30 | 0 | R | Inferior frontal gyrus (pars triangularis) |
| 8 | 912 | 0.03* | 5.20 | -12 | -12 | 8 | L | Thalamus |
| | | 0.02 | 3.65 | -14 | -20 | 6 | L | Thalamus |
| 9 | 808 | 0.03 | 4.72 | 14 | 4 | 4 | R | Pallidum^ |

MNI = Montreal Neurological Institute (MNI) standard stereotaxic space
* Indicates the peaks/sub-peaks withstanding the voxel-level FWE p < .05 threshold.
^ The anatomical region was labelled based on visual inspection of the ALE map because the coordinates fall outside the Automated Anatomical Labelling Atlas.

**Table S4.1.4.** Significant ALE clusters (cluster extent-based FWE threshold, p< .05) of convergent activation across 37 **implicit empathy for pain** experiments. Anatomical labels are derived from the Automatic Anatomical Labelling Atlas.

| Cluster | Volume (mm3) | ALE value | Z value | Peak MNI coordinates | | | Hemisphere | Anatomical region |
|---|---|---|---|---|---|---|---|---|
| | | | | X | Y | Z | | |
| 1 | 4552 | 0.06* | 8.44 | -58 | -24 | 32 | L | Supramarginal gyrus |
| 2 | 4152 | 0.04* | 6.45 | 42 | 0 | -4 | R | Insula |
| | | 0.03* | 4.95 | 40 | 24 | 0 | R | Insula |
| | | 0.03 | 4.46 | 42 | 32 | 4 | R | Inferior frontal gyrus (pars triangularis) |
| | | 0.02 | 4.07 | 38 | 12 | 2 | R | Insula |
| 3 | 4000 | 0.06* | 7.99 | 62 | -20 | 34 | R | Supramarginal gyrus |

| | | | | | | | | |
|---|---|---|---|---|---|---|---|---|
| 4 | 3672 | 0.04* | 5.97 | 48 | -64 | -8 | R | Inferior temporal gyrus |
| | | 0.03* | 5.62 | 52 | -64 | 0 | R | Middle temporal gyrus |
| 5 | 3232 | 0.04* | 5.70 | -50 | 10 | 18 | L | Inferior frontal gyrus (pars opercularis) |
| | | 0.02 | 4.38 | -50 | 4 | 28 | L | Precentral gyrus |
| | | 0.02 | 4.14 | -52 | 4 | 40 | L | Precentral gyrus |
| | | 0.02 | 3.98 | -44 | 6 | 8 | L | Insula |
| 6 | 2848 | 0.06* | 7.74 | -44 | -70 | -4 | L | Inferior occipital gyrus |
| 7 | 1576 | 0.03* | 5.38 | -30 | 24 | 4 | L | Insula |
| | | 0.02 | 3.71 | -40 | 24 | 0 | L | Inferior frontal gyrus (pars triangularis) |
| 8 | 1536 | 0.03* | 5.24 | 48 | 6 | 28 | R | Precentral gyrus |
| | | 0.02 | 4.05 | 50 | 8 | 44 | R | Precentral gyrus |
| 9 | 1464 | 0.04* | 6.18 | -14 | -96 | -6 | L | Inferior occipital gyrus |
| 10 | 1416 | 0.03 | 4.53 | 2 | 20 | 48 | R | Supplementary motor area |
| | | 0.02 | 3.79 | 8 | 12 | 58 | R | Supplementary motor area |
| | | 0.02 | 3.43 | -2 | 16 | 42 | L | Middle cingulate gyrus |
| 11 | 1288 | 0.04* | 5.96 | 18 | -94 | -4 | R | Calcarine fissure |
| 12 | 1112 | 0.03* | 5.56 | -40 | -2 | -6 | L | Insula |
| | | 0.02 | 3.23 | -40 | 10 | -4 | L | Insula |
| 13 | 928 | 0.04* | 5.90 | 22 | -4 | -14 | R | Amygdala |
| 14 | 792 | 0.03 | 4.45 | 2 | -30 | -8 | R | Brainstem^ |
| | | 0.02 | 4.40 | -2 | -30 | -6 | L | Brainstem^ |
| 15 | 768 | 0.03* | 4.94 | -2 | 4 | 30 | L | Anterior cingulate gyrus |
| 16 | 760 | 0.03 | 4.70 | -40 | -46 | 54 | L | Inferior parietal lobule |
| 17 | 688 | 0.03* | 5.23 | -6 | -16 | -2 | L | Thalamus |

MNI = Montreal Neurological Institute (MNI) standard stereotaxic space
* Indicates the peaks/sub-peaks withstanding the voxel-level FWE p < .05 threshold. This analysis revealed one additional cluster in the left amygdala ( [-22, -6, -14], ALE value 0.03, Z value 4.92, cluster size 8 $mm^3$) which did not withstand the cluster extent-based threshold.
^ The anatomical region was labelled based on visual inspection of the ALE map because the coordinates fall outside the Automated Anatomical Labelling Atlas.

**Table S4.1.5.** Significant ALE clusters (cluster extent-based FWE threshold, p< .05) of convergent activation across 47 **explicit moral reasoning** experiments. Anatomical labels are derived from the Automatic Anatomical Labelling Atlas.

| Cluster | Volume (mm3) | ALE value | Z value | Peak MNI coordinates | | | Hemisphere | Anatomical region |
|---|---|---|---|---|---|---|---|---|
| | | | | X | Y | Z | | |
| 1 | 8760 | 0.04* | 6.63 | -2 | 58 | 28 | L | Medial superior frontal gyrus |
| | | 0.03 | 4.85 | -6 | 52 | 12 | L | Anterior cingulate gyrus |
| | | 0.03 | 4.76 | 8 | 60 | 12 | R | Medial superior frontal gyrus |
| | | 0.02 | 3.55 | -20 | 48 | 32 | L | Middle frontal gyrus |
| 2 | 4144 | 0.05* | 6.92 | -50 | -62 | 24 | L | Middle temporal gyrus |
| 3 | 3512 | 0.06* | 7.81 | -2 | -56 | 30 | L | Precuneus |
| 4 | 1744 | 0.04* | 5.88 | -32 | 20 | -12 | L | Insula |
| 5 | 952 | 0.02 | 4.13 | 48 | -58 | 22 | R | Middle temporal gyrus |
| | | 0.02 | 3.56 | 54 | -54 | 14 | R | Middle temporal gyrus |
| | | 0.02 | 3.50 | 54 | -62 | 30 | R | Angular gyrus |
| | | 0.02 | 3.32 | 58 | -58 | 22 | R | Superior temporal gyrus |
| 6 | 896 | 0.02 | 4.10 | 2 | 56 | -6 | R | Medial orbitofrontal cortex |

MNI = Montreal Neurological Institute (MNI) standard stereotaxic space
* Indicates the peaks/sub-peaks withstanding the voxel-level FWE p < .05 threshold.

**Table S4.1.6.** Significant ALE clusters (cluster extent-based FWE threshold, p< .05) of convergent activation across 22 **implicit moral reasoning** experiments. Anatomical labels are derived from the Automatic Anatomical Labelling Atlas.

| Cluster | Volume (mm3) | ALE value | Z value | Peak MNI coordinates | | | Hemisphere | Anatomical region |
|---|---|---|---|---|---|---|---|---|
| | | | | X | Y | Z | | |
| 1 | 1200 | 0.03* | 5.34 | 54 | 2 | -28 | R | Middle temporal gyrus |
| | | 0.02 | 3.72 | 54 | 6 | -18 | R | Middle temporal pole |
| 2 | 728 | 0.02 | 4.32 | -50 | 8 | 10 | L | Inferior frontal gyrus (pars opercularis) |
| 3 | 720 | 0.02 | 4.62 | 2 | 58 | 18 | R | Medial superior frontal gyrus |

MNI = Montreal Neurological Institute (MNI) standard stereotaxic space
* Indicates the peaks/sub-peaks withstanding the voxel-level FWE p < .05 threshold.

**Table S4.2.1.** Significant clusters revealed by the conjunction and contrast analyses ($p < .001$ uncorrected, min. cluster size 200 $mm^3$) between **explicit** (46) and **implicit** (23) **empathy for emotion** experiments. Anatomical labels are derived from the Automatic Anatomical Labelling Atlas.

| Cluster | Volume (mm3) | ALE value | Z value | Peak MNI coordinates X | Y | Z | Hemisphere | Anatomical region |
|---|---|---|---|---|---|---|---|---|
| **Explicit & Implicit (conjunction)** | | | | | | | | |
| 1 | 720 | 0.02 | | -38 | 24 | -4 | L | Inferior frontal gyrus (pars orbitalis) |
| 2 | 472 | 0.02 | | 48 | 28 | 0 | R | Inferior frontal gyrus (pars triangularis) |
| **Explicit - Implicit** | | | | | | | | |
| 1 | 552 | | 3.54 | -12.5 | 47.3 | 37 | L | Superior frontal gyrus |
| | | | 3.24 | -6 | 52.5 | 35.5 | L | Medial superior frontal gyrus |
| | | | 3.24 | -5.2 | 52.2 | 37.2 | L | Medial superior frontal gyrus |
| **Implicit - Explicit** | | | | | | | | |
| | | | | | - | | | |

MNI = Montreal Neurological Institute (MNI) standard stereotaxic space

**Table S4.2.2.** Significant clusters revealed by the conjunction and contrast analyses ($p < .001$ uncorrected, min. cluster size 200 $mm^3$) between **explicit** (48) and **implicit** (37) **empathy for pain** experiments. Anatomical labels are derived from the Automatic Anatomical Labelling Atlas.

| Cluster | Volume (mm3) | ALE value | Z value | Peak MNI coordinates X | Y | Z | Hemisphere | Anatomical region |
|---|---|---|---|---|---|---|---|---|
| **Explicit & Implicit (conjunction)** | | | | | | | | |
| 1 | 3072 | 0.05 | | -60 | -22 | 30 | L | Supramarginal gyrus |
| 2 | 1936 | 0.05 | | 62 | -22 | 34 | R | Supramarginal gyrus |
| 3 | 1760 | 0.03 | | -52 | 10 | 14 | L | Inferior frontal gyrus (pars opercularis) |
| | | 0.02 | | -52 | 6 | 28 | L | Precentral gyrus |
| | | 0.02 | | -56 | 8 | 32 | L | Precentral gyrus |
| 4 | 976 | 0.03 | | -30 | 24 | 4 | L | Insula |
| 5 | 904 | 0.03 | | -46 | -68 | -2 | L | Inferior occipital gyrus |

| | | | | | | | | |
|---|---|---|---|---|---|---|---|---|
| 6 | 864 | 0.02 | | 0 | 18 | 48 | L/R | Supplementary motor area |
| | | 0.02 | | 4 | 18 | 52 | R | Supplementary motor area |
| | | 0.02 | | 2 | 18 | 44 | R | Middle cingulate gyrus |
| | | 0.02 | | 6 | 12 | 60 | R | Supplementary motor area |
| | | 0.02 | | -2 | 16 | 42 | L | Middle cingulate gyrus |
| 7 | 392 | 0.02 | | 36 | 26 | 2 | R | Insula |
| 8 | 328 | 0.02 | | -40 | -2 | -2 | L | Insula |
| 9 | 72 | 0.02 | | -40 | 6 | -4 | L | Insula |
| | | 0.02 | | -40 | 10 | -4 | L | Insula |
| 10 | 8 | 0.02 | | 4 | 20 | 42 | R | Middle cingulate gyrus |
| 11 | 8 | 0.02 | | 4 | 22 | 44 | R | Medial superior frontal gyrus |
| **Explicit - Implicit** | | | | | | | | |
| 1 | 488 | | 3.72 | -7.1 | 22.4 | 40.8 | L | Medial superior frontal gyrus |
| **Implicit - Explicit** | | | | | | | | |
| 1 | 752 | | 3.72 | -12.5 | -94.9 | -9.2 | L | Inferior occipital gyrus |
| | | | 3.54 | -8 | -94 | -5 | L | Calcarine fissure |
| | | | 3.35 | -13.7 | -98.7 | -4 | L | Calcarine fissure |
| 2 | 256 | | 3.35 | 20 | -5 | -18.5 | R | Hippocampus |
| | | | 3.24 | 25.3 | -4.3 | -18.3 | R | Hippocampus |

MNI = Montreal Neurological Institute (MNI) standard stereotaxic space

**Table S4.2.3.** Significant clusters revealed by the conjunction and contrast analyses (p < .001 uncorrected, min. cluster size 200 $mm^3$) between **explicit** (47) and **implicit** (22) **moral reasoning** experiments. Anatomical labels are derived from the Automatic Anatomical Labelling Atlas.

| Cluster | Volume (mm3) | ALE value | Peak MNI coordinates X | Y | Z | Hemisphere | Anatomical region |
|---|---|---|---|---|---|---|---|
| | | | **Explicit & Implicit (conjunction)** | | | | |
| 1 | 416 | 0.02 | 0 | 58 | 18 | L/R | Medial superior frontal gyrus |
| | | 0.02 | 4 | 60 | 16 | R | Medial superior frontal gyrus |
| | | | **Explicit - Implicit** | | | | |
| | | | - | | | | |
| | | | **Implicit - Explicit** | | | | |
| | | | - | | | | |

MNI = Montreal Neurological Institute (MNI) standard stereotaxic space

**Table S5.1.** Significant ALE clusters (cluster extent-based FWE threshold, p < .05) of convergent activation across a subset of 26 **ToM** experiments limited to those for which the experimental condition was harder than the control condition (**E>C**). Anatomical labels are derived from the Automatic Anatomical Labelling Atlas.

| Cluster | Volume (mm3) | ALE value | Z value | Peak MNI coordinates X | Y | Z | Hemisphere | Anatomical region |
|---|---|---|---|---|---|---|---|---|
| 1 | 3904 | 0.05* | 7.28 | -6 | 54 | 36 | L | Medial superior frontal gyrus |
| | | 0.02 | 4.18 | 2 | 56 | 16 | R | Medial superior frontal gyrus |
| | | 0.02 | 4.09 | 4 | 62 | 24 | R | Medial superior frontal gyrus |
| 2 | 3488 | 0.04* | 6.60 | -44 | 26 | -12 | L | Inferior frontal gyrus (pars orbitalis) |
| | | 0.03* | 4.90 | -52 | 24 | 6 | L | Inferior frontal gyrus (pars triangularis) |
| | | 0.02 | 4.10 | -52 | 18 | 18 | L | Inferior frontal gyrus (pars opercularis) |
| 3 | 3360 | 0.03* | 5.52 | -58 | -10 | -16 | L | Middle temporal gyrus |
| | | 0.02 | 4.77 | -54 | -2 | -24 | L | Middle temporal gyrus |
| | | 0.02 | 3.90 | -46 | 2 | -36 | L | Inferior temporal gyrus |

| | | | | | | | | |
|---|---|---|---|---|---|---|---|---|
| | | 0.02 | 3.87 | -44 | 14 | -36 | L | Middle temporal pole |
| | | 0.02 | 3.76 | -46 | 6 | -34 | L | Inferior temporal gyrus |
| 4 | 3272 | 0.05* | 7.37 | -54 | -58 | 20 | L | Middle temporal gyrus |
| 5 | 2480 | 0.03* | 6.02 | 54 | 0 | -22 | R | Middle temporal gyrus |
| | | 0.02 | 4.68 | 52 | -18 | -14 | R | Middle temporal gyrus |
| 6 | 2040 | 0.04* | 6.13 | -58 | -40 | 2 | L | Middle temporal gyrus |
| 7 | 2040 | 0.03* | 5.49 | 54 | -52 | 22 | R | Middle temporal gyrus |
| | | 0.03* | 5.33 | 58 | -54 | 16 | R | Middle temporal gyrus |
| | | 0.02 | 3.91 | 54 | -64 | 24 | R | Middle temporal gyrus |
| 8 | 1888 | 0.03* | 5.42 | -8 | 14 | 64 | L | Supplementary motor area |
| 9 | 1784 | 0.03* | 6.03 | -4 | -52 | 36 | L | Precuneus |
| | | 0.02 | 3.99 | -2 | -62 | 30 | L | Precuneus |
| 10 | 976 | 0.02 | 4.65 | 56 | 26 | 8 | R | Inferior frontal gyrus (pars triangularis) |
| | | 0.02 | 4.13 | 56 | 30 | 0 | R | Inferior frontal gyrus (pars triangularis) |
| 11 | 728 | 0.03* | 5.54 | 28 | -78 | -34 | R | Cerebellum |

MNI = Montreal Neurological Institute (MNI) standard stereotaxic space
* Indicates the peaks/sub-peaks withstanding the voxel-level FWE p < .05 threshold.

**Table S5.2.** Significant ALE clusters (cluster extent-based FWE threshold, p < .05) of convergent activation across a subset of 25 **ToM** experiments limited to those for which the experimental and control conditions were equally hard (**E=C**). Anatomical labels are derived from the Automatic Anatomical Labelling Atlas.

| **Cluster** | **Volume (mm3)** | **ALE value** | **Z value** | **Peak MNI coordinates** | | | **Hemisphere** | **Anatomical region** |
|---|---|---|---|---|---|---|---|---|
| | | | | **X** | **Y** | **Z** | | |
| 1 | 5584 | 0.05* | 7.54 | -6 | -52 | 32 | L | Posterior cingulate gyrus |
| | | 0.02 | 4.79 | 6 | -54 | 28 | R | Precuneus |
| 2 | 4424 | 0.04* | 6.61 | -50 | -58 | 24 | L | Middle temporal gyrus |
| 3 | 4240 | 0.03* | 5.59 | -4 | 56 | 20 | L | Medial superior frontal gyrus |
| | | 0.03* | 5.41 | 6 | 56 | 20 | R | Medial superior frontal gyrus |
| | | 0.03* | 5.30 | -8 | 52 | 34 | L | Medial superior frontal gyrus |
| 4 | 4016 | 0.03* | 5.42 | 54 | -48 | 18 | R | Superior temporal gyrus |

| | | | | | | | | |
|---|---|---|---|---|---|---|---|---|
| | | 0.02 | 4.67 | 54 | -64 | 12 | R | Middle temporal gyrus |
| | | 0.02 | 4.59 | 64 | -48 | 24 | R | Superior temporal gyrus |
| | | 0.02 | 3.97 | 46 | -58 | 10 | R | Middle temporal gyrus |
| | | 0.02 | 3.81 | 46 | -54 | 12 | R | Middle temporal gyrus |
| | | 0.01 | 3.37 | 54 | -58 | 30 | R | Angular gyrus |
| 5 | 2936 | 0.03* | 6.17 | -62 | -14 | -14 | L | Middle temporal gyrus |
| | | 0.02 | 4.79 | -58 | -22 | -6 | L | Middle temporal gyrus |
| | | 0.02 | 4.77 | -52 | -28 | -4 | L | Middle temporal gyrus |
| 6 | 1824 | 0.03* | 6.13 | -48 | 28 | -10 | L | Inferior frontal gyrus (pars orbitalis) |
| | | 0.02 | 4.71 | -52 | 24 | 6 | L | Inferior frontal gyrus (pars triangularis) |
| 7 | 1720 | 0.04* | 6.93 | 50 | 10 | -28 | R | Middle temporal pole |
| 8 | 1248 | 0.03* | 5.36 | 58 | -8 | -16 | R | Middle temporal gyrus |
| 9 | 976 | 0.03* | 6.19 | -26 | -80 | -36 | L | Cerebellum |
| 10 | 832 | 0.03* | 6.23 | 0 | -16 | 38 | L/R | Middle cingulate gyrus |

MNI = Montreal Neurological Institute (MNI) standard stereotaxic space
* Indicates the peaks/sub-peaks withstanding the voxel-level FWE p < .05 threshold.

**Table S5.3.** Significant clusters revealed by the conjunction and contrast analyses ($p < .001$ uncorrected, min. cluster size 200 $mm^3$) between **E>C** (26) and **E=C** (25) **ToM** experiments. Anatomical labels are derived from the Automatic Anatomical Labelling Atlas.

| Cluster | Volume (mm3) | ALE value | Z value | Peak MNI coordinates | | | Hemisphere | Anatomical region |
|---|---|---|---|---|---|---|---|---|
| | | | | X | Y | Z | | |
| | | | | **[E>C] & [E=C] (conjunction)** | | | | |
| 1 | 2208 | 0.04 | | -50 | -58 | 24 | L | Middle temporal gyrus |
| 2 | 2104 | 0.03 | | -8 | 52 | 34 | L | Medial superior frontal gyrus |
| | | 0.02 | | 2 | 56 | 16 | R | Medial superior frontal gyrus |
| | | 0.02 | | -2 | 56 | 24 | L | Medial superior frontal gyrus |
| | | 0.02 | | -6 | 56 | 26 | L | Medial superior frontal gyrus |
| 3 | 1592 | 0.03 | | -4 | -52 | 36 | L | Precuneus |
| 4 | 1344 | 0.03 | | -46 | 28 | -10 | L | Inferior frontal gyrus (pars orbitalis) |

| | | | | | | | | |
|---|---|---|---|---|---|---|---|---|
| | | 0.02 | | -52 | 24 | 6 | L | Inferior frontal gyrus (pars triangularis) |
| 5 | 1120 | 0.03 | | 54 | -50 | 20 | R | Middle temporal gyrus |
| | | 0.01 | | 52 | -58 | 26 | R | Angular gyrus |
| 6 | 632 | 0.03 | | -60 | -10 | -14 | L | Middle temporal gyrus |
| 7 | 416 | 0.02 | | 58 | -6 | -20 | R | Middle temporal gyrus |
| 8 | 352 | 0.02 | | 52 | 6 | -26 | R | Middle temporal gyrus |
| | | | | | **[E>C] – [E=C]** | | | |
| 1 | 736 | | 3.54 | -56 | -40.8 | 0.1 | L | Middle temporal gyrus |
| | | | | | **[E=C] – [E>C]** | | | |
| | | | | | - | | | |

MNI = Montreal Neurological Institute (MNI) standard stereotaxic space

**Table S5.4.** Significant ALE clusters (cluster extent-based FWE threshold, $p < .05$) of convergent activation across a subset of 14 **ToM** experiments limited to those for which the experimental condition was easier than the control condition (**C>E**). Anatomical labels are derived from the Automatic Anatomical Labelling Atlas.

| **Cluster** | **Volume (mm3)** | **ALE value** | **Z value** | **Peak MNI coordinates** | | | **Hemisphere** | **Anatomical region** |
|---|---|---|---|---|---|---|---|---|
| | | | | **X** | **Y** | **Z** | | |
| 1 | 3088 | 0.02 | 4.51 | 54 | -52 | 26 | R | Angular gyrus |
| | | 0.02 | 4.40 | 56 | -46 | 22 | R | Superior temporal gyrus |
| | | 0.02 | 4.30 | 60 | -46 | 22 | R | Superior temporal gyrus |
| | | 0.02 | 4.12 | 48 | -58 | 18 | R | Middle temporal gyrus |
| | | 0.01 | 3.31 | 46 | -46 | 22 | R | Superior temporal gyrus |
| 2 | 2840 | 0.03* | 6.78 | 6 | -54 | 32 | R | Posterior cingulate gyrus |
| 3 | 1552 | 0.03* | 6.11 | -48 | -58 | 26 | L | Angular gyrus |
| 4 | 1496 | 0.02 | 4.49 | 6 | 62 | 24 | R | Medial superior frontal gyrus |
| 5 | 696 | 0.02 | 4.10 | 46 | 18 | -28 | R | Middle temporal pole |
| | | 0.01 | 3.57 | 50 | 10 | -34 | R | Inferior temporal gyrus |

MNI = Montreal Neurological Institute (MNI) standard stereotaxic space

**Table S5.5.** Significant ALE clusters (cluster extent-based FWE threshold, p < .05) of convergent activation across a subset of 60 **ToM** experiments limited to those for which the behavioural information was known. These results represented the basis of the cluster analysis. Anatomical labels are derived from the Automatic Anatomical Labelling Atlas.

| Cluster | Volume (mm3) | ALE value | Z value | Peak MNI coordinates | | | Hemisphere | Anatomical region |
|---|---|---|---|---|---|---|---|---|
| | | | | X | Y | Z | | |
| 1 | 8552 | 0.07* | 8.78 | -8 | 54 | 34 | L | Medial superior frontal gyrus |
| | | 0.06* | 7.24 | 4 | 58 | 20 | R | Medial superior frontal gyrus |
| | | 0.05* | 6.50 | -6 | 58 | 22 | L | Medial superior frontal gyrus |
| 2 | 7936 | 0.08* | 9.26 | -6 | -52 | 34 | L | Posterior cingulate gyrus |
| | | 0.06* | 7.45 | 4 | -54 | 32 | R | Posterior cingulate gyrus |
| 3 | 7880 | 0.06* | 7.53 | 50 | 10 | -28 | R | Middle temporal pole |
| | | 0.05* | 6.92 | 54 | -2 | -22 | R | Middle temporal gyrus |
| | | 0.04* | 5.61 | 52 | -18 | -16 | R | Inferior temporal gyrus |
| | | 0.03 | 4.78 | 48 | -28 | -6 | R | Middle temporal gyrus |
| | | 0.03 | 4.12 | 52 | -36 | 0 | R | Middle temporal gyrus |
| 4 | 7880 | 0.05* | 7.08 | -60 | -12 | -14 | L | Middle temporal gyrus |
| | | 0.04* | 5.64 | -56 | -40 | 2 | L | Middle temporal gyrus |
| | | 0.04* | 5.23 | -52 | 2 | -28 | L | Middle temporal gyrus |
| | | 0.03 | 4.60 | -52 | -28 | -4 | L | Middle temporal gyrus |
| | | 0.03 | 4.12 | -46 | 10 | -34 | L | Inferior temporal gyrus |
| 5 | 7104 | 0.09* | 10.27 | -50 | -58 | 24 | L | Middle temporal gyrus |
| 6 | 6784 | 0.06* | 8.06 | 54 | -50 | 20 | R | Middle temporal gyrus |
| 7 | 3352 | 0.06* | 7.79 | -46 | 28 | -10 | L | Inferior frontal gyrus (pars orbitalis) |
| | | 0.05* | 6.25 | -52 | 24 | 6 | L | Inferior frontal gyrus (pars triangularis) |
| 8 | 1632 | 0.05* | 6.66 | -26 | -78 | -36 | L | Cerebellum |
| 9 | 1560 | 0.03 | 4.78 | -2 | 46 | -16 | L | Gyrus rectus |
| | | 0.03 | 4.07 | 2 | 56 | -18 | R | Gyrus rectus |
| | | 0.03 | 4.01 | -8 | 52 | -2 | L | Middle orbitofrontal cortex |
| 10 | 1480 | 0.04* | 5.96 | 56 | 28 | 2 | R | Inferior frontal gyrus (pars triangularis) |
| 11 | 1176 | 0.03 | 4.53 | -6 | 12 | 62 | L | Supplementary motor area |
| 12 | 1144 | 0.05* | 7.04 | 28 | -78 | -34 | R | Cerebellum |

MNI = Montreal Neurological Institute (MNI) standard stereotaxic space
* Indicates the peaks/sub-peaks withstanding the voxel-level FWE $p < .05$ threshold. This analysis revealed one additional cluster in the middle cingulate gyrus ( [0, -16, 40], ALE value 0.04, Z value 5.88, cluster size 136 $mm^3$) which did not withstand the cluster extent-based threshold.
